# A targetable dependency on nonsense-mediated decay for proteostasis and immune control in small cell lung cancer

**DOI:** 10.64898/2026.03.31.715503

**Authors:** Lucia A. Torres Fernández, Volker Boehm, Joel Kaufmann, Agnieszka Rumińska, Jonas P. Becker, Maria Garcia Marquez, Beaunelle de Bruijn, Christian Müller, Nazanin Alavinejad, Laura Lovric, Johanna Bihler, Kian R. Weihrauch, Laura Kaiser, Marcel Schmiel, Lukas Maas, Filippo Beleggia, Olta Ibruli, Bianca Göbel, Vignesh Sakthivelu, Graziella Bosco, Liu Fanyu, Ariadne Androulidaki, Silvia von Karstedt, Maria Cartolano, Johannes Brägelmann, Justinas Valiulis, Joshua D’Rozario, Nina Wobst, Katja Höpker, Felix John, Jürgen Wolf, Alexander Quaas, Stefan Knapp, Poorya Davoodi, Anna Schöllhorn, Cécile Gouttefangeas, Hans-Georg Rammensee, Angelika B. Riemer, Hans C. Reinhardt, Hans A. Schlößer, Niels H. Gehring, Martin Peifer, Roman K. Thomas, Julie George

**Affiliations:** Department of Translational Genomics, University of Cologne, Faculty of Medicine and University Hospital Cologne, Weyertal 115b, 50931 Cologne, Germany; Institute for Genetics, Department of RNA biology, University of Cologne, 50674 Cologne, Germany; Center for Molecular Medicine Cologne (CMMC), University of Cologne, 50931 Cologne, Germany; Division of Immunotherapy and Immunoprevention, German Cancer Research Center (DKFZ), 69120 Heidelberg, Germany; Immunopeptidomics Unit, National Center for Tumor Diseases (NCT), NCT Heidelberg, a partnership between DKFZ and University Hospital, 69120 Heidelberg, Germany; Department of General, Visceral, Thoracic and Transplantation Surgery, University Hospital Cologne, 50931 Cologne, Germany; Institute of Pathology, Faculty of Medicine and University Hospital Cologne, University of Cologne, 50931 Cologne, Germany; Mildred School of Oncology Aachen-Bonn-Cologne-Düsseldorf (MSSO ABCD), Faculty of Medicine and University Hospital Cologne, University of Cologne, 50937 Cologne, Germany; Department I of Internal Medicine, Lung Cancer Group Cologne, Center for Integrated Oncology, University Hospital Cologne, 50931 Cologne, Germany; Department of Hematology and Stem Cell Transplantation, University Hospital Essen, 45147 Essen, Germany; CECAD Cluster of Excellence, Faculty of Medicine and University Hospital Cologne, 50931 Cologne, Germany; Department III of Internal Medicine, Lung Cancer Group Cologne, University Hospital Cologne, 50931 Cologne, Germany; Structural Genomics Consortium (SGC) and Institute of Pharmaceutical Chemistry, Goethe-University Frankfurt, 60438 Frankfurt am Main, Germany; Institute for Immunology, University of Tübingen, 72076 Tübingen, Germany; Cluster of Excellence iFIT (EXC2180) "Image-Guided and Functionally Instructed Tumor Therapies", 72076 Tübingen, Germany; German Cancer Consortium (DKTK) and German Cancer Research Center (DKFZ) partner site Tübingen, 72076 Tübingen, Germany; Molecular Vaccine Design, German Center for Infection Research (DZIF), partner site Heidelberg, 69120 Heidelberg, Germany; Department of Otorhinolaryngology, Head and Neck Surgery, University Hospital Cologne, 50937 Cologne, Germany

## Abstract

Small cell lung cancer (SCLC) is among the deadliest cancers with excessive somatic alterations, expectedly resulting in immunogenic epitopes. However, patient benefit from immunotherapy is limited. We found abundant frameshift mutations in SCLC, regarded as highly immunogenic, counterbalanced by a hyperactive nonsense-mediated decay (NMD) pathway, responsible for frameshift-mRNA degradation. NMD activity correlated with tumor mutational burden (TMB) across cancers, suggesting that SCLC depends on NMD for cellular homeostasis and immune evasion. NMD inhibition impaired SCLC proliferation and induced ER stress-dependent apoptosis in a TMB-dependent manner, thereby enabling pharmacological inhibition *in vivo* which effectively controlled tumor growth. By integrating genome and transcriptome sequencing with MHC-I immunopeptidomics and functional *in vitro* and *in vivo* assays, we identified that NMD inhibition boosted neoantigen expression and presentation by tumor cells and increased T cell recognition, thus enhancing overall tumor immunogenicity and further improving immunotherapy efficacy *in vivo*. Our work shows that SCLC – as a TMB^high^ cancer – relies on NMD for survival and immune escape, uncovering a novel TMB-dependent tractable vulnerability for this devastating disease.

## Introduction

Lung cancer is the leading cause of cancer-related death worldwide^1^. Small cell lung cancer (SCLC) is the most aggressive form of lung cancer accounting for approximately 15-20% of all patients. SCLC is characterized by rapid metastatic spread and an extremely poor 5-year survival rate of less than 7%^1–5^. While most patients with SCLC respond well to initial treatment with platinum-based chemotherapy, relapse almost inevitably occurs within a few months^6^. Combinations of immune-checkpoint blockade (ICB) with chemotherapy show improved outcomes, but overall survival benefits remain moderate at best^7,8^. Furthermore, given as a single agent after relapse from first-line chemotherapy, ICB showed poor responses with less than 30% of the patients responding to the treatment^9^. This limited efficacy is intriguing, as SCLC is one of the cancers with the highest frequency of somatic mutations, which are widely expected to generate neoantigens, thereby facilitating immune cell recognition^10–12^. Most recently, DLL3-targeted T-cell engagers showed exceptional durable responses in patients with relapsed SCLC^13,14^, supporting immunotherapeutic approaches in this deadly disease.

We previously performed comprehensive large-scale genome sequencing studies of SCLC^2–5^ and identified key somatic drivers and patterns of genome evolution during treatment resistance^3^. The underlying data have yielded a catalogue of SCLC genome alterations at unprecedented resolution. A key question arising from these studies is how tumor cells can tolerate extremely large numbers of mutated proteins, which can be cytotoxic and thus affect cellular homeostasis. Furthermore, a high somatic mutational burden leads to a large number of mutated proteins, which can be processed into neoantigens and confer immunogenicity. Specifically, frameshift and intron-retention mutations can generate novel peptides of various lengths and are thus considered particularly immunogenic^15,16^. Consequently, they have been strongly associated with the clinical response of cancer patients to ICB^12,16–18^. In this context, the nonsense-mediated decay (NMD) pathway plays an important role as an evolutionary conserved RNA surveillance mechanism that detects and subsequently degrades transcripts with premature termination codons (PTCs), including those arising from frameshifts and intron retentions^19–24^. PTC-containing transcripts are typically marked by a downstream exon junction complex, which interacts with two core NMD pathway factors, the phosphatidylinositol 3-kinase-related kinase SMG1 and the RNA helicase UPF1. Such interactions lead to the activation of SMG1 kinase activity and the subsequent phosphorylation of UPF1, which in turn recruits several downstream NMD factors to promote mRNA degradation via endonucleolytic cleavage, deadenylation and decapping^19–24^. The NMD pathway thus prevents the accumulation of mutant misfolded proteins with not only immunogenic, but also cytotoxic potential^10,25^. In cancers with a large number of somatic mutations, inhibition of NMD might therefore have a particularly profound impact on both tumor cell viability and tumor immunogenicity. Supporting the latter, previous studies demonstrated an immune-dependent control of tumor growth upon NMD inhibition^15,26–28^. Furthermore, NMD-escaping frameshifts^18^, as well as mutations in NMD pathway components^27^, have been associated with response to ICB, and reduced NMD activity was found to correlate with increased tumor immune infiltration^29^.

In recent years, it has become apparent that the NMD pathway has roles beyond canonical quality control of aberrant transcripts, participating in the homeostatic regulation of up to 10-25% of the transcriptome^19,20,30,31^. Consequently, NMD inhibition might not only lead to the upregulation of neoantigen-encoding transcripts, but also impacts a multitude of functional mRNAs participating in the regulation of transcription, splicing, translation, proliferation, differentiation, survival and response to several types of stress responses^19,23,30,32–38^. Thus, elucidating the role of the NMD pathway in cancer, and how NMD perturbation affects tumor progression in specific cancer types will advance our mechanistic understanding of tumorigenesis and inform novel strategies for therapeutic intervention.

We therefore sought to determine how SCLC cells cope with cell-autonomous and immunological implications of a high TMB, under the hypothesis that this tumor type might be particularly dependent on the NMD pathway.

## Results

### High TMB correlates with elevated NMD pathway activity

We first performed a comprehensive analysis of the number and types of somatic mutations in SCLC compared to other cancer entities by integrating genomic data from our previous studies^2–5^ and from publicly available sources^39^. Our analysis confirmed that SCLC has one of the highest mutational burdens - known to be caused by tobacco smoke toxicity^3,40^ -, with a remarkably high number of frameshift and nonsense mutations (**Fig. 1A** and **Table S1A-B**). These two types of mutations generate altered transcripts which, upon expression, lead to truncated and/or misfolded proteins that can perturb cellular homeostasis^41^. Furthermore, frameshift mutations may serve as a source for highly immunogenic peptides^12,17,18^. Transcripts with nonsense and frameshift mutations are typically degraded by the nonsense-mediated decay (NMD) pathway. Since altered NMD activity has been reported in some cancer types^29,42^, we aimed at quantifying the levels of NMD pathway activity in SCLC.

**Figure 1.**
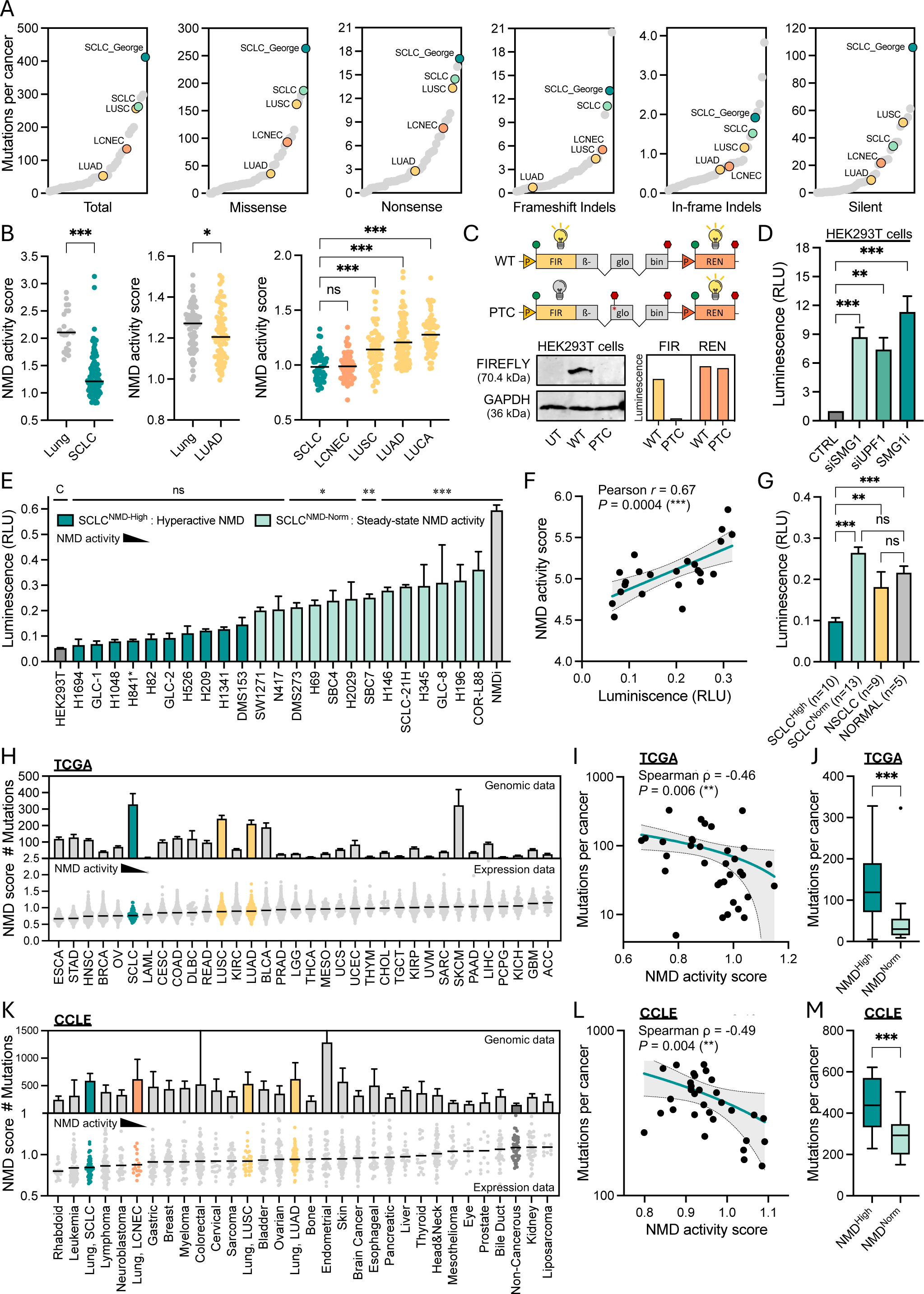
High TMB correlates with elevated NMD pathway activity (see also **Fig. S1** and **Table S1**). **A)** Average number of mutations per cancer for the indicated mutation types, obtained from COSMIC database and our own SCLC cohort (SCLC_George). NSCLC cancer subtypes are also highlighted: LUAD = lung adenocarcinoma; LUSC = lung squamous cell carcinoma; LCNEC = large cell neuroendocrine carcinoma. **B)** NMD activity score calculated for lung cancer datasets. NMD activity scores represent the expression of 50 NMD-sensitive genes ^43^ and are thereby inversely proportional to NMD activity (see **Methods**). Left panel: normal lung (n=22) *vs* SCLC (n=109)^54,5^, compared by Mann-Whitney test. Middle panel: normal lung (n=72) *vs* LUAD (n=72)^55^, compared by Mann-Whitney test. Right panel: SCLC (n=50) *vs* LCNEC (n=66), LUSC (n=60), LUAD (n=101) and lung carcinoid = LUCA (n=61)^2^ compared by one-way ANOVA. **C)** Schematic representation of the luciferase reporter assay using a wild-type (WT) NMD-insensitive or a premature termination codon (PTC)-containing NMD-sensitive dual luciferase reporter (upper panel)^56–58^; western blot for HEK293T cells untransfected (UT) or transfected with WT- and PTC-reporter constructs (bottom left); quantified Firefly (FIR) and Renilla (REN) luciferase signals in adjacent HEK293T cells (bottom right). **D)** Luciferase reporter assay for CTRL *vs* NMD-inhibited HEK293T cells via siSMG1, siUPF1 and SMG1i treatment (see **Methods** and Fig. 2A), compared by one-way ANOVA (n≥3). RLU = relative light units, calculated as (FIR/REN)_PTC_/(FIR/REN)_WT_. Luminescence thus represent the expression of the PTC-reporter and is thereby inversely proportional to NMD activity. **E)** Luciferase reporter assay for human SCLC (n=21) and SMARCA-UT (n=2, name marked with*) cell lines, with untreated and SMG1i-treated HEK293T serving as controls for high and low NMD activity, respectively (grey bars). Comparisons to control (C) HEK293T cells by one-way ANOVA (n≥3). Population frequency analysis enabled the definition of high (SCLC^NMD-High^) and steady-state NMD activity (SCLC^NMD-Norm^) groups (see **Fig. S1E**). **F)** Pearson correlation of luciferase-based NMD activity (from **E**) and transcriptome-based NMD activity scores in SCLC cell lines (see **Fig. S1C**). **G)** Luciferase reporter assay for human SCLC (NMD^High^ n=10; NMD^Norm^ n=13; from **E**), NSCLC (n=9) and non-neoplastic cells (NORMAL, n=5) (see cell lines list in **Table S1C)** compared by one-way ANOVA (n≥5). **H)** Mutation load (upper panel) and transcriptome-based NMD activity scores (lower panel) for all TCGA cancers and SCLC data^5^. Other lung cancers (LUSC and LUAD) are highlighted. **I)** Spearman correlation for median data from **H**. **J)** Median number of mutations per cancer for NMD^High^ (n=15) *vs* NMD^Norm^ (n=19) cancers from **H**, compared by Mann-Whitney test. NMD^High^ and NMD^Norm^ groups were defined by NMD activity scores below or above the TCGA average, respectively. **K)** Mutational load (upper panel) and transcriptome-based NMD activity scores (lower panel) for all CCLE cancers. Other lung cancers (LCNEC, LUSC and LUAD) are highlighted. **L)** Spearman correlation for median data from **K**. **M)** Median number of mutations per cancer for NMD^High^ (n=19) and NMD^Norm^ (n=13) cancers from **K**, compared by Mann-Whitney test. NMD^High^ and NMD^Norm^ groups were defined by NMD activity scores below or above the CCLE average, respectively. For all graphs, error bars represent the SEM, and P-values are specified as ***P<0.001; **P <0.01; *P<0.05; ns=non-significant.

To this end, we first applied a transcriptome-based metric for NMD activity^43^, here termed the *NMD activity score*. This score estimates NMD pathway activity based on the repression of 50 previously validated NMD-sensitive genes^44–47^, with high cellular NMD activity corresponding to stronger repression of such genes and consequently lower scores (see **Methods**). We assessed the validity of these 50 NMD-sensitive genes by referring to data from other studies^48–53^ and confirmed consistent upregulation of these 50 genes upon NMD inhibition across multiple human cell types (**Fig. S1A**). We next applied this metric to determine NMD activity scores for several published lung cancer datasets^2,5,54,55^ and found that both SCLC and lung adenocarcinoma (LUAD) exhibited higher NMD pathway activities (lower NMD activity scores) than normal lung, with SCLC displaying the highest NMD activity across all lung cancer types (**Fig. 1B**, *P* < 0.001). Accordingly, patient-derived SCLC specimens exhibited higher NMD activity levels than most normal healthy tissues within the GTEx database, including normal lung (**Fig. S1B**).

To further validate the transcriptome-based NMD activity score, we next sought to functionally assess the activity of the NMD pathway in SCLC cell lines. We analyzed the transcriptomes of 45 human SCLC cell lines (**Table S1C**) to determine their NMD activity score, exhibiting a wide range of activity that enabled us defining groups of presumably hyperactive (*NMD^High^*) and normal/basal (*NMD^Norm^*) NMD pathway activity (**Figs. S1C-D**). We then employed a previously established and well-validated luciferase-based cellular reporter system^56–58^ (see **Methods**) to experimentally measure NMD activity in SCLC cell lines (**Figs. 1C-E** and **S1E-F**). Since embryonic tissues are known to exhibit high NMD activity^36,59^, HEK293T cells were used as a reference for high NMD activity, whereas inhibition of NMD in HEK293T cells served as a control for low NMD activity (**Figs. 1C-E**). The elevated NMD activity in HEK293T cells was demonstrated by strong repression of a β-Globin reporter with a premature termination codon (PTC) – but not of a wild-type β-Globin reporter (WT) (**Fig. 1C**, see **Methods**) – which was de-repressed upon NMD inhibition (**Figs. 1D** and **S1F**). Luciferase reporter assays across 23 human cell lines again revealed groups of distinct NMD activity (**Figs. 1E** and **S1E**), with multiple SCLC cell lines showing a high NMD activity, comparable to that quantified in HEK293T cells. Importantly, the cellular NMD activity measured by luciferase reporter assays strongly correlated with the transcriptome-based NMD activity scores (**Fig. 1F**, *r* = 0.67; *P* < 0.001). Furthermore, luciferase reporter assays confirmed a higher NMD activity (stronger PTC-reporter repression) in NMD^High^ SCLC than in NSCLC cell lines and several non-neoplastic *NORMAL* cell lines, including blood-derived samples and primary lung fibroblasts (**Fig. 1G** and **Table S1C**), further distinguishing cells with hyperactive (*NMD^High^*) *versus* normal/basal (*NMD^Norm^*) levels of NMD activity. Collectively, these results validate the reliability of the transcriptome-based NMD activity score and confirm a high cellular NMD pathway activity in SCLC.

To contextualize such a high NMD activity, we calculated NMD activity scores for SCLC and all cancer types profiled within the TCGA database (**Figs. 1H-J** and **Table S1D**) and correlated this with matched genomic data (see **Methods**). NMD pathway activity was moderately but significantly correlated with the TMB across different cancer types, with SCLC exhibiting the highest mutational load and among the highest NMD pathway activities (reflected by low NMD scores, **Fig. 1I**, ρ = -0.46; *P* < 0.01). While overall NMD efficiency may be shaped by multiple cancer type–specific factors, these data reveal a contribution of TMB to the observed variation in the levels of NMD activity across cancers. Consistently, somatic mutations were significantly higher in NMD^High^ *vs* NMD^Norm^ cancer types (**Fig. 1J**). Similar results were observed when analyzing data from the *Cancer Cell Line Encyclopedia* (CCLE) database (**Figs. 1K-M**).

Collectively, we identify hyperactive NMD as a hallmark of SCLC and observe a significant correlation between TMB and cellular NMD pathway activity across different cancer types. We therefore hypothesized that cancers harboring a high number of disruptive mutations, such as SCLC, may upregulate their NMD activity – via yet unknown mechanisms – to maintain cellular homeostasis and/or as a mechanism to evade immune recognition. This prompted us to investigate next whether SCLC, given its high TMB, depends on NMD for survival.

### NMD inhibition causes a specific and reproducible accumulation of PTC-transcripts

To further investigate a dependency on the NMD pathway in SCLC, we next sought to inhibit NMD by siRNA-mediated knockdown of the two key NMD components SMG1 and UPF1, as well as by pharmacological inhibition of SMG1 kinase activity using the well-described small molecule inhibitor SMG1i-11j^60,61^, hereafter referred to as SMG1i (**Fig. 2A**). The use of these orthogonal approaches enabled us ruling out off-target effects while distinguishing overall NMD-dependent effects from merely SMG1-dependent effects. To consider the previously reported role of proposed SCLC subtypes^62^, we included several neuroendocrine-high (NE^High^) and neuroendocrine-low (NE^Low^) human cell lines representing SCLC subtypes based on the expression of key lineage transcription factors *ASCL1* (NE^High^), *NEUROD1* (NE^High^), *POU2F3* (NE^Low^) and *YAP1* (NE^Low^)^62^ (**Fig. S2**). Our analyses also included the cell line H841, which was formerly considered a YAP1+ SCLC line, but has recently been reclassified as a SMARCA4-undifferentiated (UT) tumor^63^. Despite this histological reclassification, H841 remains comparable to other TMB^High^SCLC cell lines in terms of elevated NMD activity (see again **Fig. 1E**). Furthermore, it serves as a valuable model for a TMB^High^ cancer due to its high transfection efficiency which is typically lower in most SCLC cell lines^64^.

**Figure 2.**
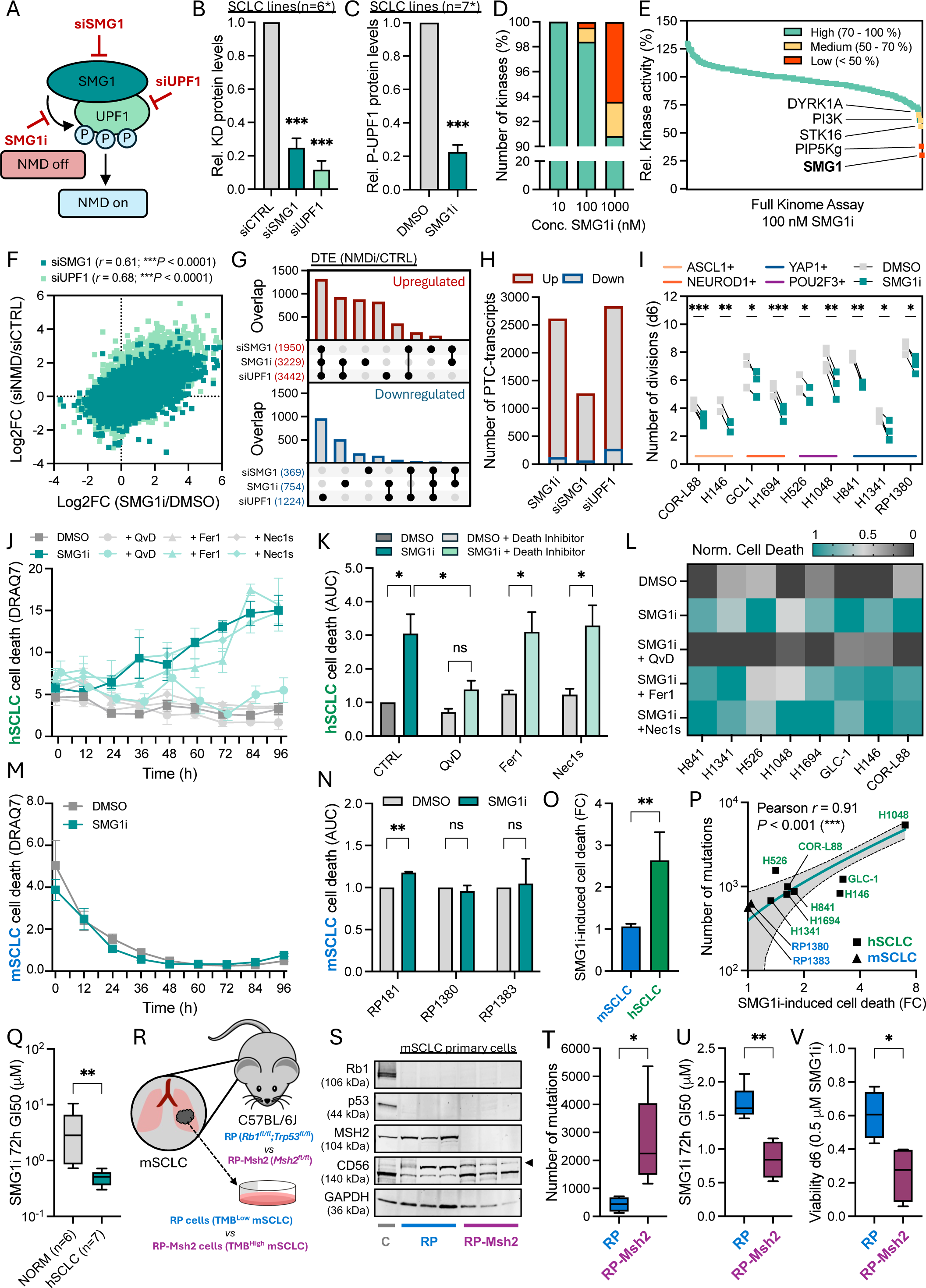
NMD inhibition impairs tumor cell proliferation and triggers apoptotic cell death in TMB^High^ tumor cells (see also **Figs. S2-S6** and **Table S2**). **A)** Schematic representation of the NMD pathway initiation and approaches for NMD inhibition by siRNA-knockdown (KD) of *SMG1* (siSMG1) or *UPF1* (siUPF1), or chemical inhibition of the SMG1 kinase (SMG1i). **B)** Quantification of SMG1 and UPF1 KD protein levels (relative to GAPDH) in n=6 TMB^high^ cell lines 72h post-transfection with siSMG1 and siUPF1, respectively. Band densitometry analysis was done on western blots from **Fig. S2B**, with comparisons by unpaired t-tests. **C)** Quantification of P-UPF1 protein levels (relative to UPF1/HSP90) in n=7 TMB^high^ cell lines 24h post-treatment with SMG1i in western blots from **Fig. S2D**, with comparisons by unpaired t-tests. \***B** and **C** include not only SCLC cell lines but also H841 and H1341, recently reclassified as SMARCA4-UT and small cell carcinoma of the cervix, respectively^125^. **D-E)** Kinome assay for SMG1i treatment on 436 kinases showing the number of kinases (%) affected and the degree of inhibition (**D**), highlighting affected kinases at 100 nM (**E**). Colors in **D-E** indicate high (>70%), medium (50-70%) and low (<50%) relative kinase activity (see **Table S2A** and **Fig. S3**). **F)** Pearson correlation of transcriptomic changes (bulk RNA-Seq, DESeq2 analysis for differential transcript expression (DTE)) in H841 cells following chemical NMD inhibition (0.5µM SMG1i for 24h) *vs* genetic KD (siRNA-mediated for 72h). Each dot represents the Log2FC expression of a single transcript in SMG1i *vs* DMSO (n=3, *x*-axis) against the Log2FC for siUPF1 or siSMG1 *vs* siCTRL (n=3, *y*-axis). **G)** UpSet plots for differential transcript expression (DTE) overlaps in H841 following chemical NMD inhibition with SMG1i and siRNA-mediated KD. For each treatment (siSMG1, siUPF1, SMG1i), numbers of differentially up/downregulated transcripts (Log2FC>|1|, Padj<0.05, n=3) are indicated by colored numbers in brackets (see **Table S2D-E**). **H)** Number of PTC-containing transcripts (annotated as *nonsense mediated decay* or *retained intron* by GENCODE) significantly up/downregulated upon NMD inhibition (see **Table S2B-C**). **I)** Proliferation assays for CTRL *vs* SMG1i-treated cell lines showing the number of cell divisions after 6 days based on the loss of the proliferation dye eFluor670 fluorescence intensity over time (see **Methods** and **Fig. S4**). Comparisons by paired t-tests (n≥3). **J-L)** Cell death assays in human cell lines (hSCLC) monitored by live-cell imaging (DRAQ7 staining) for 4 days. Representative death curves shown for H146 cells treated with DMSO *vs* SMG1i and additional cell death inhibitors (QvD=apoptosis inhibitor; Fer1=ferroptosis inhibitor; Nec1s=necroptosis inhibitor) **(J)**; the area under the curve (AUC) was quantified and compared by unpaired t-tests (n≥3), as shown exemplified for H146 **(K)**, with average normalized cell death plotted as a heatmap for n=7 human TMB^High^ cell lines **(L)**. **M-N)** Cell death assays in DMSO *vs* SMG1i-treated murine TMB^Low^ cell lines (mSCLC) monitored by live-cell imaging (DRAQ7 staining) for 4 days. Representative death curves shown for RP1380 cells **(M)** with AUC quantification compared by unpaired t-tests (n≥3) for n=3 mSCLC RP cell lines (see **Fig. S5**) **(N). O)** SMG1i-induced cell death in n=3 TMB^Low^ mSCLC RP cell lines *vs* n=7 TMB^High^ hSCLC cell lines (from **N** and **L**, respectively) compared by Mann-Whitney test. **P)** Pearson correlation of SMG1i-induced cell death (from **N** and **L**) with total number of exome mutations (see **Table S5A**). **Q)** Sensitivity to SMG1i treatment of n=6 non-neoplastic (NORM) cell lines *vs* n=7 hSCLC cell lines compared by Mann-Whitney test (see **Fig. S6A-B**). **R)** Schematic representation for mSCLC cell lines derived from lung tumors arising in RP (*Rb1^fl/fl^;Trp53^fl/fl^*, TMB^low^) and RP-Msh2 mice (*Rb1^fl/fl^;Trp53^fl/fl^;Msh2^fl/fl^,* TMB^high^) following Adeno-CMV-Cre inhalation. **S)** Western blot for n=3 RP and n=3 RP-Msh2 cell lines representative cell lines compared to MEFs as control (C) probing for Rb1, p53 and MSH2, and confirming expression of the SCLC-specific NCAM1/CD56 140KDa-isoform^126^. **T-V)** RP *vs* RP-Msh2 cell lines comparing total number of exome mutations (**T**), sensitivity to SMG1i treatment (**U**) (see **Fig. S6C-D**), and CTG-viability assays following 6 days of SMG1i treatment (**V**), by Mann-Whitney tests. For all graphs, error bars represent the SEM, and P-values are specified as ***P<0.001; **P <0.01; *P<0.05; ns=non-significant.

Knockdown of SMG1 and UPF1 (siSMG1 and siUPF1) led to efficient downregulation of both proteins in all human cell lines tested (n=6, **Figs. 2B** and **S2B-C**). Furthermore, treatment with SMG1i reduced UPF1 phosphorylation in all human cell lines tested (n=7, **Figs. 2C** and **S2D-E**). Both, genetic and chemical inhibition of NMD effector proteins reduced NMD activity, indicated by de-repression of the artificial PTC-reporter (**Figs. S2F-G**), as well as upregulation of the NMD target *TBL2*^65–67^ (**Fig. S2H**), but not the non-target *GAPDH^67^*(**Fig. S2I**). Transcriptome sequencing of NMD inhibited samples, including cell lines (n=5) and primary SCLC specimens (n=4), also confirmed upregulation of all 50 NMD-sensitive genes previously used to calculate NMD activity scores following NMD inhibition (see **Fig. S1A**), thus further validating the employed NMD inhibition approaches as well as the transcriptome-based NMD activity metric.

In comparison to other widely used NMD inhibitors^68,69^, SMG1i induced stronger suppression of NMD activity^60,61^ (see again **Fig. S2F**). We confirmed the on-target effect of the SMG1i compound on the purified full length SMG1 kinase by an *in vitro* phosphorylation assay (see **Methods** and **Fig. S3A**) and further evaluated the selectivity of SMG1i via an enzymatic full kinome assay on 436 recombinant kinases (**Figs. 2D-E**, **S3B** and **Table S2A**), confirming SMG1 as the top-target at 100nM, followed by inhibition of the lipid kinase PIP5Kg (Relative kinase activity (RKA) < 50%) and only moderate inhibition (RKA = 50-70%) of three other human kinases (PI3K, STK16 and DYRK1A); even at 1 μM of SMG1i, – a rather high concentration for a biochemical assay on purified components – fewer than 10% of all tested kinases were affected (RKA < 70 %) (**Figs. 2D-E**, **S3B** and **Table S2A**), underscoring this compound’s remarkable selectivity. We further evaluated the off-target inhibition of selected kinases in cellular assays, and found that the kinase activity of neither DYRK1A, PI3K nor STK16 were affected at 0.5 μM SMG1i, as quantified by phosphorylation of their respective substrates^70–72^ (**Fig. S3C**). Furthermore, a nanoBRET reporter assay revealed a 10-fold higher IC50 for inhibition of PIP5Kg over the IC50 reported for the on-target inhibition of NMD via luciferase reporter assays (**Figs. S3D-F**). Collectively, these data establish SMG1i as a potent and highly selective SMG1 kinase inhibitor.

We next compared chemical with genetic NMD inhibition using transcriptome sequencing data of the SMARCA4-UT cell line H841. Transcriptomic changes induced by chemical NMD inhibition with SMG1i treatment tightly correlated with those induced by siRNA-mediated knockdown of both SMG1 (siSMG1i; *r* = 0.61; *P* < 0.0001) and UPF1 (siUPF1; *r* = 0.68; *P* < 0.0001) (**Fig. 2F** and **Tables S2B-C**), with major overlaps observed in differentially expressed transcripts especially affecting the upregulated transcriptome (**Fig. 2G** and **Tables S2D-E**). As expected, PTC-containing transcripts were mostly upregulated upon NMD inhibition (**Fig. 2H** and **Tables S2B-C**), thus highlighting the reliability and on-target effect of the distinct approaches employed to inhibit the NMD pathway. Altogether, our data show that targeting key components of the NMD pathway using orthogonal strategies induces reproducible, NMD-specific transcriptomic changes that lead to a massive accumulation of aberrant transcripts in SCLC.

### NMD inhibition impairs tumor cell proliferation and triggers apoptotic cell death in TMB^High^ tumor cells

The accumulation of thousands of altered transcripts could disrupt cellular homeostasis due to the production of truncated and misfolded proteins with cytotoxic potential^25,41,73,74^. Proteotoxic stress is typically mitigated by the NMD pathway at the transcript level and at the protein level through downstream proteostatic mechanisms triggered by the ER stress response pathway. We hypothesized that TMB^High^ cancers such as SCLC may depend on enhanced NMD activity to protect against such cytotoxicity. We therefore evaluated tumor cell-autonomous effects of NMD inhibition on SCLC proliferation and survival. To this end we employed several human SCLC cell lines (hSCLC) as well as murine cancer cells (mSCLC) derived from a genetically engineered SCLC mouse model with conditional biallelic inactivation of *Rb1* and *Trp53* (*Rb1*^fl/fl^;*Trp53*^fl/fl^), abbreviated as RP mice^75^.

All cell lines tested exhibited reduced proliferation (see **Methods**) upon SMG1i treatment (**Fig. 2I and Fig. S4**). Employing live-cell imaging and dead-cell staining (see **Methods**), we observed increased cell death upon SMG1i treatment in multiple human TMB^High^ cell lines, which was rescued by inhibiting apoptosis with the pan-caspase inhibitor QvD^76^, but not when using the ferroptosis inhibitor Fer1^77^ or the necroptosis inhibitor Nec1s^78^ (**Figs. 2J-L**). We thus confirmed that NMD inhibition triggers cell death predominantly through apoptosis. This effect was observed for all included TMB^High^ cell lines despite their different levels of basal NMD activity (see again **Fig. 1E**). Furthermore, we did not noted differences in the vulnerability to NMD inhibition across SCLC subtypes.

Most strikingly, human and murine SCLC cell lines exhibited differential vulnerability to NMD inhibition (**Figs. 2M-O** and **Figs. S5**). Murine SCLC (mSCLC) tumor cells derived from RP mice (*Rb1*^fl/fl^;*Trp53*^fl/fl^, in short RP cells) mirror SCLC morphology and biology, but do not yield large numbers of somatic alterations - a defining hallmark of hSCLC - and therefore are classified as TMB^Low^ ^75,79^. While proliferation of RP cells such as RP1380 and RP1383 was reduced upon NMD inhibition (**Fig. 2I** and **Figs. S5A-B**), cell death was absent in these TMB^Low^ mSCLC RP models (**Figs. 2M-O** and **Figs. S5C-D**). Furthermore, the degree of SMG1i-induced cell death observed in different SCLC cellular models correlated tightly with their mutational load (**Fig. 2P,** *r* = 0.91; *P* < 0.001). In line with this observation, TMB^High^ hSCLC cell lines were more sensitive to NMD inhibition than several non-neoplastic normal (*NORM*) cell lines considered as TMB^Low^, including human peripheral blood mononuclear cells (PBMCs), human pancreatic epithelial duct cells (HPDE), human bronchial epithelial cells (HBEC), human lung fibroblasts (MRC-9) and murine fibroblasts (MEF and NIH3T3)) (**Figs. 2Q** and **S6A-B**). These observations collectively suggest that cells with a high burden of somatic alterations – such as SCLC - are particularly vulnerable to NMD inhibition likely due to the cytotoxic effects of somatically altered proteins. To further investigate a potential impact of TMB on the vulnerability to NMD inhibition, and to rule out that different vulnerability in murine and human cells is species-specific, we tested mSCLC cells derived from a mismatch repair-deficient SCLC mouse model termed RP-Msh2 mice (*Rb1^fl/fl^;Trp53^fl/fl^;Msh2^fl/fl^*, in short RP-Mh2 cells), characterized by somatic hypermutation^80^. We thus compared the vulnerability to NMD inhibition in TMB^Low^ RP cells and TMB^High^ RP-Mhs2 cells derived from RP and RP-Msh2 mice lung tumors, respectively (**Figs. 2R-S, S2J-K** and **Table S1C**). Indeed, RP-Msh2 cells with a higher load of somatic mutations (**Figs. 2T and S2L**) were more sensitive to NMD inhibition than RP cells (**Figs. 2U-V** and **S6C-D**), thus further confirming a TMB-dependent vulnerability to NMD inhibition in SCLC.

### NMD inhibition triggers ER stress-related cell death in TMB^High^ tumor cells

Our data show that NMD inhibition triggers apoptotic tumor cell death in a TMB-dependent manner. We therefore hypothesized that in highly mutated tumor cells, cell death upon NMD inhibition was caused by cytotoxic effects of accumulated aberrant proteins, thus causing ER stress-related cell death. To further determine the cause of cell death we examined transcriptome sequencing data of several SMG1i-treated TMB^High^ human cancer cells, including cell lines and primary cells from patients with SCLC (**Tables S1C** and **S3**). We observed consistent transcriptional changes induced by SMG1i treatment across all samples, with substantial overlaps again largely affecting the upregulated transcriptome (**Fig. 3A** and **Tables S4A-B**). In agreement with our previous observations, pathway enrichment analysis after NMD inhibition (see **Methods**) pointed to significantly deregulated critical cellular processes including the cell cycle, intrinsic apoptosis and ER stress-related pathways (**Figs. 3B** and **S7A-B,** and **Tables S3C-D**). The ER stress signature was characterized by upregulated genes, which participate in multiple cellular pathways to control proteotoxic stress caused by the accumulation of misfolded proteins, including protein folding via chaperones, protein degradation via autophagy and the ubiquitin-proteasome system, interference with global protein translation, and ultimately, in case of unresolved ER stress, activation of apoptosis^74^ (**Figs. S7A-B** and **Table S4C**). Confirming our hypothesis, NMD inhibition caused a massive accumulation of PTC-transcripts in all TMB^High^ samples (**Fig. 3C** and **Table S3**), paralleled by upregulation of chaperones (BiP/GRP78), autophagy (p62 degradation) and ER stress-dependent apoptosis effectors (ATF4-CHOP-BAK1 axis) (**Figs. 3D** and **S7C**).

**Figure 3.**
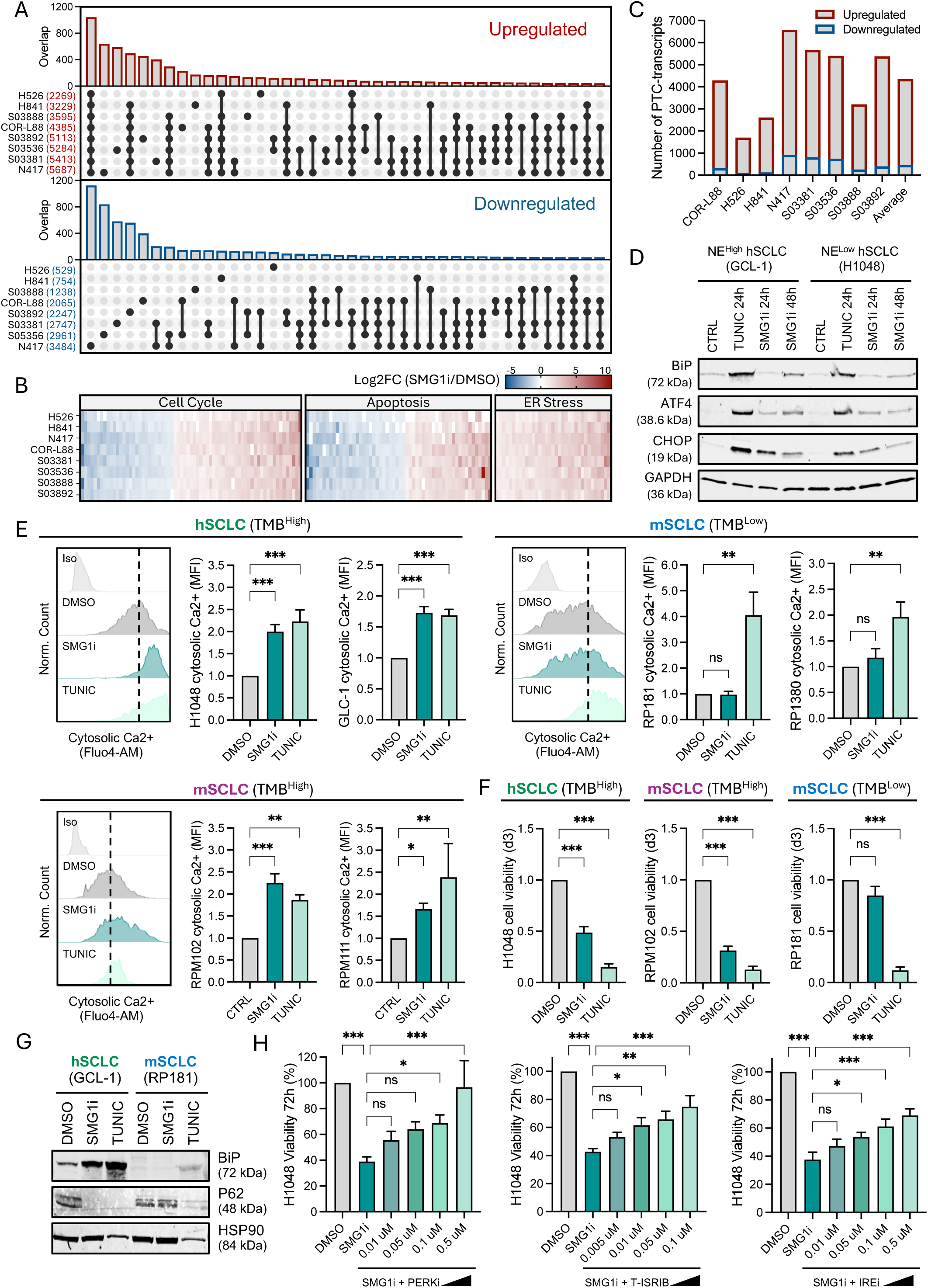
NMD inhibition triggers ER stress-related death in TMB^High^ tumor cells (see also **Fig. S7** and **Tables S3-4**). **A-C)** Transcriptome sequencing data for indicated n=4 SCLC cell lines and n=4 patient-derived SCLC tumor cells treated with DMSO *vs* SMG1i for 24h (**Table S3**). UpSet plots for differential transcript expression (DTE) overlaps (**Tables S4A-B**) with the number of differentially up/downregulated transcripts (Log2FC>|1|, Padj<0.05, n=3) indicated by colored numbers within brackets **(A)**. Heatmap for differentially expressed transcripts involved in cell cycle, apoptosis and ER stress-related processes identified through pathway enrichment analysis (**Figs. S7A-B** and **Tables S4C-D**) **(B)**; number of PTC-transcripts (GENCODE annotation as *nonsense-mediated decay* or *retained intron,* **Table S3**) significantly up/downregulated per sample **(C). D)** Western blot for ER stress response factors in hSCLC cell lines with NE^Low^ and NE^High^ phenotypes treated with SMG1i or tunicamycin (TUNIC; positive control for ER stress) **(Figs. S7C)**. **E)** Cytosolic Ca2+ levels in hSCLC (TMB^High^), mSCLC RP (TMB^Low^) and mSCLC RP-Msh2 (TMB^High^) cell lines measured by flow cytometry (Fluo-4AM staining) 72h post-treatment with SMG1i or TUNIC (positive control for ER stress-related death), compared by one-way ANOVA (n≥3), and accompanied by representative Fluo-4AM histograms (MFI=mean fluorescence intensity; Iso=Isotype control). **F)** CTG-viability assays of representative hSCLC (TMB^High^), mSCLC RP (TMB^Low^) and mSCLC RP-Msh2 (TMB^High^) cell lines 72h post-treatment with SMG1i or TUNIC, compared by one-way ANOVA (n≥4). **G)** Western blot for BiP chaperone accumulation and p62 degradation as readouts for protein folding and autophagy activation, respectively, after SMG1i or TUNIC treatment for 72h in representative hSCLC and mSCLC cell lines. **H)** CTG-viability assays of DMSO *vs* SMG1i-treated H1048 cells rescuing SMG1i-induced death with increasing concentrations of the indicated ER stress response inhibitors (see **Methods**), analyzed by one-way ANOVA (n≥4). For all graphs, error bars represent the SEM, and P-values are specified as ***P<0.001; **P <0.01; *P<0.05; ns=non-significant.

Activation of ER stress-related cell death induces apoptosis through calcium release from the ER, which causes mitochondrial membrane permeabilization^81,82^. Further confirming the activation of ER-related cell death, NMD inhibition in TMB^High^ hSCLC and mSCLC RP-Mhs2 cell lines caused increased cytoplasmic calcium levels to the same extent as treatment with tunicamycin (**Fig. 3E-F**), which served as a positive control for massive protein misfolding and ER stress-induced death^83,84^. By contrast, TMB^Low^ mSCLC RP cell lines, which did not undergo cell death upon NMD inhibition (**Figs. 2M-O**, **Fig. 3F and S5C-D**), did also not show altered cytoplasmic calcium levels (**Fig. 3E**), or activation of protein folding or autophagy following NMD inhibition (**Fig. 3G**), thus supporting the view that the number of detrimental mutations in murine RP cells is not sufficient to trigger ER stress-related cell death. Finally, inhibition of several ER stress sensors rescued SMG1i-induced cell death in TMB^High^ hSCLC in a concentration-dependent manner (**Fig. 3H**). Our data thus indicate that NMD inhibition in a TMB^High^ cellular context likely causes an intolerable accumulation of misfolded proteins, thereby triggering ER stress-dependent apoptosis in hypermutated cancer cells.

### NMD inhibition *in vivo* results in a TMB-dependent control of tumor growth

We have uncovered a TMB-dependent vulnerability to NMD inhibition in cancer cells, thus pointing to a potential therapeutic window for *in vivo* treatment. We therefore implanted human and murine SCLC cells subcutaneously in immunocompromised NSG (NOD scid gamma) mice and inhibited NMD using the small molecule KVS0001, a derivative of SMG1i with similar potency (**Figs. 4A** and **S8A**) but enhanced solubility^28^. Treatment was initiated once tumors reached 100 mm^3^ in a 5-days-on-2-days-off regimen with 50 mpk (mg compound per kg body weight) KVS0001 for a maximum of three weeks. The mice did not show any signs of toxicity, with only an initial weight drop followed by a fast recovery (**Fig. 4B**). As suggested by our previous experiments (see again **Figs. 2P-Q**), these *in vivo* results thus confirm a high tolerability to systemic NMD inhibition.

**Figure 4.**
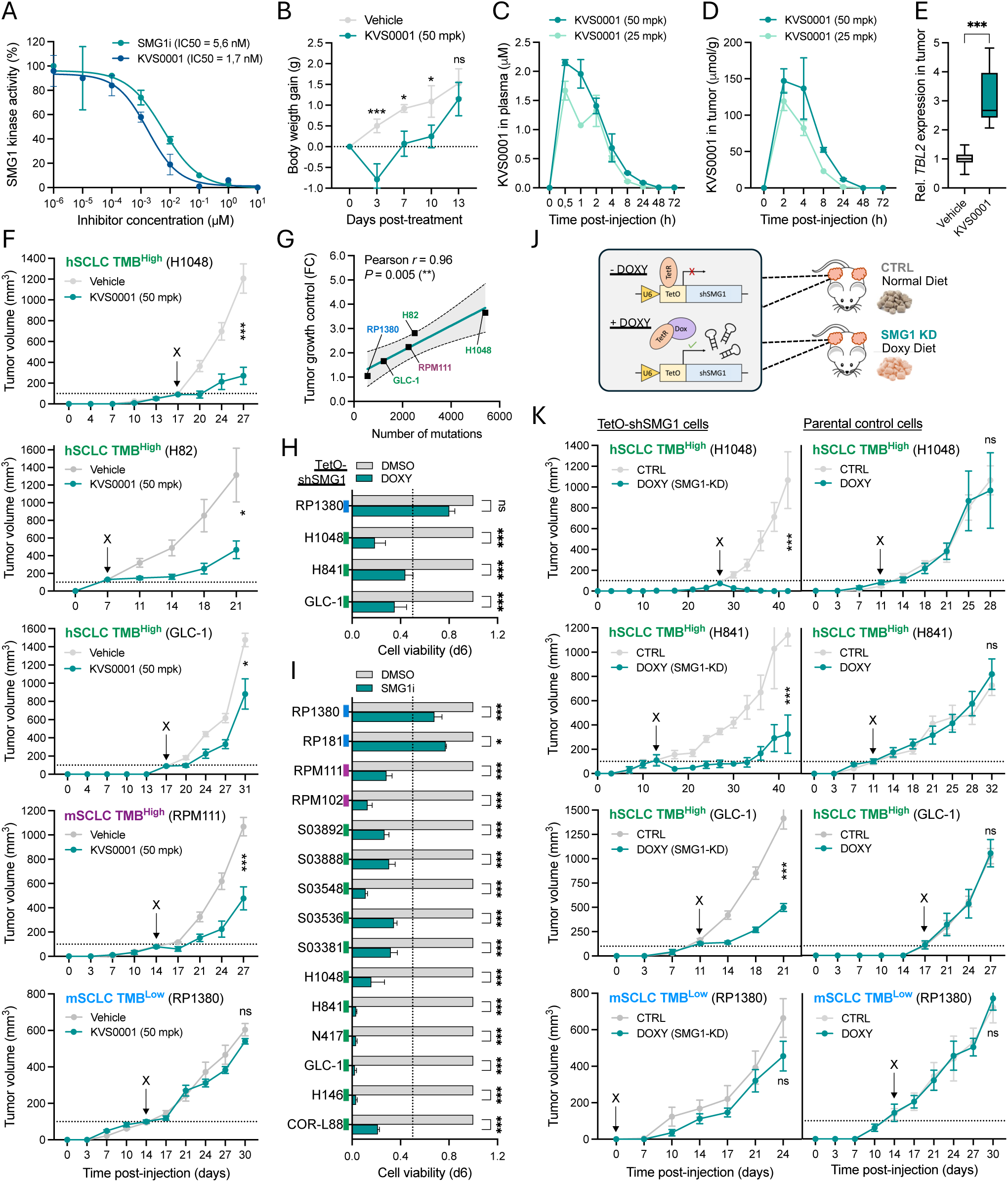
NMD inhibition *in vivo* results in a TMB-dependent control of tumor growth (see also **Fig. S8**). **A)** Enzymatic *in vitro* phosphorylation assay for full-length recombinant SMG1 protein using p53-Ser15 as a phosphorylation substrate^127^ in the presence of increasing SMG1i and KVS0001 concentrations (see **Fig. S8A**). Absolute IC50 was calculated by non-linear adjustment of the inhibitory curve. **B-F)** *In vivo* pharmacological NMD inhibition treatment of NSG mice with KVS0001 monitoring **(B)** systemic toxicity through weight changes at 50mpk KVS0001 applied in a 5-days-on-2-day-off regimen, with each time point compared by unpaired t-tests (n=16); **(C-D)** pharmacokinetics (PK) after a single shot intraperitoneal (ip) injection of KVS0001 at the indicated concentrations (n=3 mice per timepoint), quantifying KVS0001 levels by mass spectrometry in plasma **(C)** and inside subcutaneous tumors **(D)** (see **Methods** and **Fig. S8B)**; **(E)** intratumor NMD inhibition shown by qPCR for the NMD target *TBL2* in control *vs* 50mpk KVS0001-treated tumors 2h post-injection, compared by Mann-Whitney test (n=6); **(F)** tumor growth for NSG mice treated with control *vs* KVS0001 in n=3 hSCLC and n=2 mSCLC xenograft models. Treatment was initiated at tumor sizes of 100mm^3^ (indicated by “X”) and conducted on a 5-days-on-2-day-off regimen, Statistics represent unpaired t-tests on AUC (n≥4). **G)** Pearson correlation between total number of exome mutations and tumor growth control (fold change in end tumor size of control *vs* KVS0001 treatment from **F**). **H-I)** *In vitro* CTG-viability assays for (**H**) TetO-SMG1 n=1 mSCLC RP and n=3 hSCLC cell lines following doxycycline-induced SMG1-KD for 6 days (**I**) and for chemical NMD inhibition with SMG1i for 6 days in n=2 mSCLC RP, n=2 mSCLC RP-Msh2, n=6 hSCLC cell lines and n=5 patient-derived tumor cells, with comparisons by Two-way ANOVA (n≥3). **J)** Schematic representation of tumor-targeted genetic NMD inhibition *in vivo* employing doxycycline-inducible SMG1-KD models (TetO-shSMG1 cell lines) grown as subcutaneous tumors in NSG mice fed with either normal or doxycycline-diet (see **Fig. S8C-J**). **K)** Tumor growth for NSG mice xenograft models for TetO-shSMG1 cell lines (**left**) and parental control cell lines (**right**) fed with normal or doxycycline-diet initiated at tumor sizes of 100mm^3^ (indicated by “X”; note that for RP1380-TetO-shSMG1 model, doxycycline treatment started directly after tumor cell injection, to act as a proper control for experiments shown in Fig. 6E and 6G). Statistics represent unpaired t-tests on AUC (n≥4). For all graphs, error bars represent the SEM, and P-values are specified as ***P<0.005; **P <0.01; *P<0.05; ns=non-significant.

Despite rapid clearance of the KVS0001compound, micromolar concentrations were detected in the plasma (**Figs. 4C** and **S8B**), and intra-tumoral concentrations (**Figs. 4D** and **S8B**) were sufficient to elicit NMD inhibition *in vivo* as shown by de-repression of *TBL2* mRNA (**Fig. 4E**), a *bona fide* NMD target. NMD inhibition *in vivo* led to decreased tumor growth in TMB^High^ cancer cells, including three hSCLC cell lines (H1048, H82, GLC-1) and a hypermutated mSCLC RP-Msh2 cell line (RPM111), but not in a TMB^Low^ mSCLC RP cell line (RP1380) (**Fig. 4F**). The degree of tumor growth control significantly correlated with the number of mutations (**Fig. 4G**), which confirmed a TMB-dependent vulnerability of cancer cells to NMD inhibition *in vivo*.

To rule out any possible off-target effects of the employed chemical NMD inhibitor KVS0001 and to specifically assess tumor-specific consequences of NMD pathway inhibition, we also created genetic NMD inhibition models based on doxycycline-shRNA-inducible knockdown (KD) of SMG1 (in short TetO-shSMG1 cells) (**Fig. S8C**). Murine and human TetO-shSMG1 models showed *in vitro* detectable SMG1 knockdown and reduced phosphorylation of UPF1 after 72h of doxycycline treatment (**Figs. S8D-E**), resulting in effective inhibition of NMD activity as shown by de-repression of *TBL2* mRNA (**Fig. S8F**). Genetic NMD inhibition via SMG1-KD strongly reduced the viability of all TMB^High^ human TetO-shSMG1 cell lines, but not of the TMB^Low^ murine TetO-shSMG1 RP cell line (**Fig. 4H**). Similar results were obtained following chemical NMD inhibition with SMG1i, which caused pronounced reduction in cell viability (<50%) in human and murine TMB^High^cell lines as well as in patient-derived primary SCLC cells, while only mild effects were observed in murine TMB^Low^ RP cells after six days of treatment (**Fig. 4I**).

We then implanted human and murine TetO-shSMG1 cells subcutaneously in immunocompromised NSG mice, which either received a normal or a doxycycline-containing diet once growing tumors reached a size of 100 mm^3^ (**Figs. 4J** and **S8G**). Doxycycline treatment led to effective tumor-targeted SMG1 protein KD *in vivo* (**Figs. S8H-I**) without causing any notable diet-related adverse effects (**Fig. S8J**). Doxycycline treatment effectively reduced tumor growth in all TMB^High^ human TetO-shSMG1 tumors, but not in the TMB^Low^ TetO-shSMG1 murine RP cell line or in any parental control cells (**Fig. 4K**). Thus, both systemic pharmacological NMD inhibition and tumor-specific genetic NMD inhibition strongly attenuate the growth of established TMB^High^ tumors *in vivo*.

Together, our data show that highly mutated cells require the NMD pathway to maintain cellular homeostasis, thus exposing a novel TMB-dependent, tumor cell-intrinsic vulnerability to NMD inhibition. Consequently, NMD inhibition in SCLC causes a TMB-related unresolved ER stress, which triggers apoptotic cell death. Through complementary approaches we demonstrate a cell-autonomous NMD-dependent control of tumor cell survival in TMB^High^ cancer cells *in vitro* and *in vivo*.

### NMD inhibition upregulates multiple neoantigen-encoding transcripts

In addition to the newly described TMB-dependent tumor cell-autonomous vulnerability to NMD inhibition, interference with the NMD pathway has been shown to enhance immunogenicity in several cancer types^18,26–28^. This mechanism has not yet been explored in SCLC, where it may be particularly relevant given the limited efficacy of immunotherapy^7,8^. In light of the massive increase of altered transcripts caused by NMD inhibition in SCLC (**Figs. 3A-C** and **Table S3**), we next aimed at dissecting expression and upregulation of tumor-specific mutated transcripts following NMD inhibition, which could contribute to the generation of neoantigens and thereby enhance immune recognition.

We therefore first analyzed exome sequencing data of primary SCLC specimens and cell lines including TMB^High^ SCLC and SMARCA4-UT cells (**Fig. S9A** and **Table S5A**) to identify candidate neoantigens in each sample. Beyond the somatic alterations characteristic of this tumor type^2–5^ (**Fig S9A** and **Table S5A**, e.g., loss of the tumor suppressors *TP53* and *RB1*), potentially immunogenic neoantigen mutations (in short *NeoAg*) constituted 67% of all mutations – including missense substitutions, in-frame indels, frameshift indels and splice mutations (**Figs. 5A left panel, 5B** and **S9B-E** and **Table S5B**). We used the information for missense, in-frame and frameshift mutations to generate sample-specific *in-silico* peptide libraries (**Table S5C,** see **Methods**). Such libraries consisted of short mutant peptide sequences that were then used to predict MHC-I-binders within all possible overlapping 8-14mers (Xmers) using the online MHC-I-binding prediction tool *NetMHCpan EL 4.1* of the immune epitope database (IEDB)^85,86^. In agreement with previous MHC-I binder predictions^87–89^, around 1.1% (>60000) of all generated Xmers were predicted mutant MHC-I-binders (predicted NeoAg) composed of strong (24%) and weak binders (76%) (**Figs. 5A left panel, 5B** and **S9F-I, Table S5D**). The absolute number of predicted MHC-I binders per sample (**Figs. S10A-D** and **Table S5B**) was highly influenced by the number of mutations and the number of different MHC-I alleles employed for the prediction (**Fig. S10E**). We thus estimated the *neoantigen yield* as the number of MHC-I binders per mutation and MHC-I allele and determined frameshift mutations to yield the highest number of MHC-I binders (**Fig. S10F**). Notably, in-frame mutations were more frequent in cell lines than in primary tumor specimens (**Figs. S9B-C** and **S10C**), likely due to the absence of matched normal controls, which can cause germline variants to be misclassified as somatic (see **Methods** and **Table S5B**). However, when assessing proportions of mutations for Xmers and predicted MHC-I binders, no differences were observed between cell lines and primary tumor specimens (**Figs. S9D-I**), thus supporting the validity of using cell lines to study immunogenic mutations.

**Figure 5.**
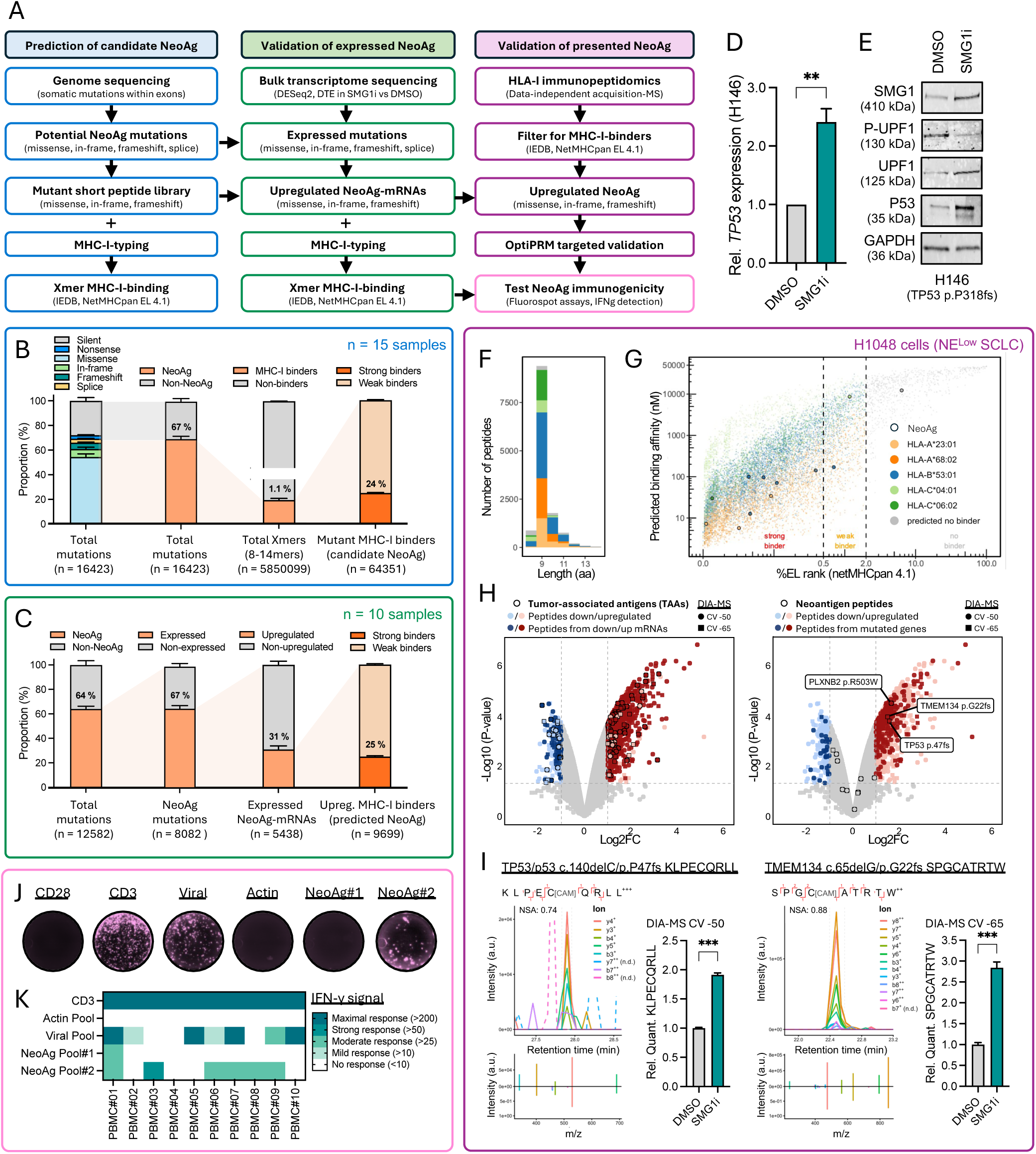
NMD inhibition upregulates expression and presentation of immunogenic neoantigens (see also **Figs. S9-14** and **Table S5**). **A)** Flowchart depicting the steps for the prediction of candidate neoantigens (NeoAg) from exome sequencing data (**left**, results shown in **B**), the expression of NeoAg-encoding genes based on bulk-RNAseq data (**middle**, results shown in **C**) and the validation of NeoAg presentation via MHC-I immunopeptidomics (**right**, results shown in **F-I**) followed by NeoAg recognition by T cells tested by FluoroSpot assays (results shown in **J-K**). **B)** Prediction of candidate NeoAg in SCLC tumors and cell lines based on exome sequencing data and MHC-I-binding modeling in n=15 samples (see **Fig. S9-10** and **Tables S5A-D**). **C)** Validation of NeoAg expression and upregulation based on exome and transcriptome sequencing of DMSO *vs* SMG1i-treated SCLC samples in n=10 samples (see **Fig. S11-13** and **Tables S5B** and **S5E**). **D-E)** Upregulation of an example frameshift *TP53*/p53 NeoAg mRNA (qPCR, n=3) **(D)** and protein (p.318fs, truncated protein) **(E)** upon NMD inhibition in H146 cells. **F-I)** MHC-I immunopeptidomics by data-independent acquisition mass spectrometry (DIA-MS) in the hypermutated SCLC cell line H1048 (n=3). **(F)** Peptide length distribution for identified MHC-I binders. **(G)** MHC-I binding predictions by NetMHCpan EL 4.1 with identified *bona fide* NeoAg highlighted (see also **Fig. S14A**). **(H)** Volcano plots showing all significantly up/downregulated MHC-I binders identified by DIA-MS using two different compensation voltages (CV-50 and CV-65) which enabled capturing different peptide repertoires; **(left)** highlights peptides derived from de-regulated mRNAs and de-regulated tumor associated antigens (TAAs); (**right**) highlights peptides derived from mutated genes and presented NeoAg (see **Fig. S14B-D** and **Table S5F**). **(I)** DIA-MS ion chromatograms for presented frameshift neoantigens (**top left**) and comparison to *in silico* predicted fragment spectra (**bottom left**) with relative peptide amounts in DMSO *vs* SMG1i-treated H1048 cells compared by unpaired t-test (n=3) (**right**) (see targeted validation in **Fig. S14E-F**). Intensity shown as arbitrary units (a.u). For the TP53 frameshift epitope in the DMSO samples, data was imputed as it was below the DIA-MS detection limit. **J-K)** FluoroSpot showing human TCR reactivities in 10 healthy-donor PBMCs for two NeoAg peptide test pools: Pool#1 contains n=3 upregulated NeoAg captured by MHC-I immunopeptidomics in H1048 and Pool#2 contains n=136 predicted frameshift NeoAg from integrating H1048 RNA-seq, MHC-I immunopeptidomics and MHC-I binding modeling data (see **Methods** and **Table S5G**). Representative reactions based on IFN-γ detection are shown for PBMC#9 **(J)** and quantified for all donors by subtracting background fluorescence of the CD28 negative control from all other conditions **(K),** with positive responses considered for IFN-γ signals with ≥10 spots after background subtraction^128^. For all graphs, error bars represent the SEM, and P-values are specified as ***P<0.001; **P <0.01; *P<0.05; ns=non-significant.

Within the predicted potential NeoAg repertoire, many mutated genes may not be expressed due to epigenetic silencing. We therefore sought to validate the expression and NMD-dependent regulation of sample-specific NeoAg upon NMD inhibition (see **Methods** and **Tables S5B** and **S5E**). By examining matched mutational and transcriptional data (**Figs. 5A middle panel** and **5C**), we observed that around 67% of all NeoAg-encoding genes were expressed at mRNA levels (43% of total mutations) (**Figs. 5C** and **S11A-B**, **Tables S5B** and **S5E**), and of these, 31% were significantly upregulated upon NMD inhibition (13% of total mutations) (**Fig. 5C** and **S11C-F, Tables S5B** and **S5E**).

We observed that not only frameshift and splice mutant transcripts were upregulated upon NMD inhibition, but also a high number of missense and in-frame mutant transcripts were found to be NMD-sensitive (**Figs. S11E-F** and **Table S5E**). It is known that NMD can regulate up to 25% of the transcriptome^30,31^, and some of those genes may, in this context, contain missense or in-frame mutations, as shown by a 20% overlap between genes mutated and genes regulated by the NMD pathway (i.e. *GENCODE*-annotated as NMD targets) in the same sample (**Figs. S12A-B**). In support of this notion, we found multiple transcripts consistently upregulated after NMD inhibition in all samples tested (**Table S4A**), many of which contained either a missense or an in-frame mutation in only one of the samples (**Fig. S12C-D**), again indicating that these genes are generally regulated by the NMD pathway, regardless of the presence of any mutation. Indeed, split analysis of wildtype (WT) and mutant (MUT) allele expression counts for upregulated transcripts with missense mutations showed a comparable regulation of both alleles, thus confirming that NMD regulates those missense transcripts independently of the harbored mutation, whereas – as expected – a preferential regulation of the MUT allele was apparent for nonsense mutations (**Fig. S12E-F**).

This concurrent regulation of transcripts with missense and in-frame mutations by the NMD pathway may provide an additional advantage in promoting tumor immune recognition. We thus next predicted NeoAg derived from upregulated missense, in-frame and frameshift mutated transcripts, and found an average of 970 predicted mutant MHC-I binders upregulated per sample (**Figs. 5C** and **S11G-I,** and **Table S5B**). These predicted NeoAg may be presented on MHC-I in all included SCLC samples given that MHC-I binding predictions are 90% accurate^86,90,91^ and all included samples express *MHC-I (HLA-A/B/C)* transcripts (**Figs. S11J-K**). Although previous studies reported low MHC-I expression specifically in NE^High^ SCLC tumors^92,93^, our analysis in a large panel of human cell lines shows substantial expression of *MHC-I* mRNA in most tested SCLC samples - including multiple NE^High^ cell lines - which correlated with protein expression levels (**Figs. S13A-E**) and effectively reflected cell surface protein expression (**Figs. S13F**). Moreover, *MHC-I* expression levels in SCLC patient samples is comparable to that observed across other cancer types in the TCGA dataset (**Figs. S13G-H**). Although immune evasion in SCLC has been attributed to low MHC-I expression^94–97^ and loss of heterozygosity (LOH) at the human *HLA-I* locus indeed occurs in 13% of SCLC cases^55^ – yet lying below the pan-cancer *HLA-I* LOH average (17%)^98^ – our results reveal a substantial MHC-I expression in multiple human SCLC cell lines and tumor specimens and support a potentially functional MHC-I antigen presentation in these samples.

Last, we did not observe any relevant association of different transcriptional SCLC subtypes with TMB levels, the levels of NeoAg-encoding mRNAs upregulated upon NMD inhibition or the proportion of predicted NeoAg (**Figs. S9-11**). Altogether, our experiments show that, in addition to cell-intrinsic effects on viability, NMD inhibition also leads to upregulation of NeoAg-encoding transcripts, thereby boosting the predicted NeoAg landscape across all four SCLC transcriptional subtypes as well as all tested cell lines and SCLC patient-derived tumor samples.

### NMD inhibition shapes the MHC-I-presented immunopeptidome and upregulates presentation of highly immunogenic neoantigens

We next examined how upregulation of mutated transcripts affects neoantigen presentation in SCLC following NMD inhibition. Although mRNA and protein levels for a given gene do not always correlate, upregulation of neoantigen-encoding mRNAs is expected to result in upregulated protein levels^99,100^. Consistently, inhibition of NMD in the SCLC cell line H146, which harbors a PTC-mutation in *TP53 (c.953_971del; p.P318fs)*, led to upregulation of mutant mRNA (**Fig. 5D**) resulting in increased truncated p53 protein levels (**Fig. 5E**). However, even upregulated mutant proteins still require highly orchestrated complex mechanisms to result in presentation by MHC-I and to ultimately elicit recognition by specific T cell receptors (TCRs). We therefore investigated the presentation and recognition of upregulated neoantigens following NMD inhibition using MHC-I immunopeptidomics and FluoroSpot assays (**Fig. 5A, right panel**).

To increase the chances of capturing *bona fide* neoantigens, we employed the human SCLC cell line H1048, which harbors high levels of somatic alterations and naturally expresses high levels of MHC-I (see again **Figs. S11K** and **S13A-F**). We conducted MHC-I-immunoprecipitations in DMSO *vs* SMG1i treated cells followed by data-independent acquisition mass spectrometry (DIA-MS) identification of eluted peptides (see **Methods**). As expected, captured peptides sizes ranged from 8-14 aa, with a clear preference for 9mer binders (**Fig. 5F**). Most MHC-I-bound peptides (93.4%) were predicted binders for at least one HLA-I allele present in H1048 cells (**Fig. 5G**) and such binders were enriched for amino acid sequence motifs expectedly corresponding to their binding HLA-I allele (**Fig. S14A**).

The number of distinct pulled down peptides was comparable for DMSO and SMG1i treated cells (**Fig. S14B**), but their presentation levels were quantitatively different, with 1004 significantly upregulated and 208 significantly downregulated peptides presented in SMG1i-treated H1048 cells as compared to DMSO controls (**Fig. 5H** and **Table S5F**). Intersection of MHC-I immunopeptidomics data with transcriptome sequencing data revealed that 76.3% of all upregulated peptides presented in SMG1i-treated samples were already upregulated at mRNA levels (**Fig. 5H, left,** highlighted in **dark red, Table S5F**), further demonstrating that transcriptome changes induced upon NMD inhibition indeed shape the immunopeptidome. Of note, several tumor-associated antigens (TAAs), rarely expressed in normal adult healthy tissues (see **Methods**), were differentially presented in H1048 cells and may contribute to the overall immunogenicity of this cell line (**Fig. 5H, left,** highlighted with **black borders, Table S5F**).

Intersection of MHC-I immunopeptidomics and exome sequencing data in H1048 cells revealed 1329 presented peptides which were derived from mutated genes. 24.3% of all significantly upregulated MHC-I peptides originated from these mutated genes (**Fig. 5H, right,** highlighted in **dark red, Table S5F**), with their allelic tumor fraction reflecting the probability for such peptides to derive from the mutant allele (**Table S5A**). Among those, a total of 13 different *bona fide* neoantigens derived from 11 missense and 2 frameshift mutations were identified by MHC-I immunopeptidomics (**Figs. S14C-D**), 3 of which were found significantly upregulated in SMG1i-treated cells, including two mutant peptides derived from frameshift mutations in *TP53* (p.P47fs; AF=0.47) and *TMEM134* (p.G22fs; AF=0.07) and one mutant peptide derived from a missense mutation in *PLXNB2* (p.R503W; AF=0.24) (**Figs. 5H right** and **5I**). We next aimed at validating presented neoantigens using targeted proteomics (see **Methods**) including these 13 identified peptides (Targeted Pool#1, **Table S5G**) as well as 58 additional frameshift neoantigens (Targeted Pool#2, **Table S5G**) which were selected based on predictions from RNA-sequencing and MHC-I immunopeptidomics data (see **Methods**). Using this approach, we validated 12 of the 13 presented neoantigens (note that the one neoantigen that could not be validated was a predicted non-binder), and identified two additional mutant peptides upregulated in SMG1i-treated H1048 cells, both originating from a frameshift mutation in *HNRNPL* (p.P337fs; AF=0.36) (**Fig. S14E-F**).

Finally, we conducted FluoroSpot assays to evaluate the immunogenicity of the 3 identified upregulated presented neoantigens (NeoAg Pool#1, **Table S5G**) as well as of 136 predicted upregulated frameshift neoantigens (NeoAg Pool#2, **Table S5G**). To this end, human TCR reactivities against neoantigen peptide pools were determined for 10 healthy PBMC donors based on T cell-secreted IFN-γ detection (see **Methods**). TCR reactivities were identified for NeoAg Pool#1 in 1/10 donors and for NeoAg Pool #2 in 6/10 donors (**Fig. 5J-K**). These results thus confirm T cell recognition of NMD-regulated neoantigens and highlight the remarkably high immunogenicity of frameshift peptides.

Altogether, our data demonstrate that, in addition to a TMB-dependent cell-autonomous vulnerability, NMD inhibition upregulates mutant transcripts in SCLC, which in turn leads to increased presentation of MHC-I-ligated neoantigens that can elicit potent immune responses.

### NMD inhibition increases tumor immunogenicity eliciting T cell-mediated responses

We next sought to evaluate whether changes in the neoantigen landscape induced by NMD inhibition lead to increased tumor immunogenicity through T cell-mediated recognition of tumor cells. To this end, we first established *in vitro* co-cultures of the MHC-I^High^ and TMB^High^ tumor cell line H841 with allogeneic PBMCs from healthy donors. Like autologous T cell responses against infected or malignant cells, alloreactive T cells can recognize a broad range of peptide-MHC combinations on foreign cells requiring three activation signals: TCR signaling (CD3), co-stimulatory signaling (CD28) and cytokine signaling (IL-2)^101–103^. Therefore, co-cultures were conducted under either naive condition (unstimulated, negative control), primed with CD28 and IL-2 (tumor killing takes place only upon antigen recognition) or artificially activated with CD28, IL-2 and CD3 (positive control for tumor killing). Successful T cell activation was apparent by T cell cluster formation for co-cultures with primed and artificially activated T cells (**Fig. 6A**). We confirmed a high MHC-I protein expression in H841 cells and created antigen presentation-deficient H841 cells via CRISPR/Cas9-based knockout (KO) of the *beta-2-microglobulin* (*B2M*) gene resulting in a lack of MHC-I expression on the cell surface which thus allowed for assessing the MHC-I-dependency of the observed effects (**Fig. 6B**, **Fig S15A**). Importantly, pharmacological inhibition of NMD with either SMG1i or KVS0001 on PBMCs alone did not affect T cell survival. Upon full artificial activation, we noted a difference in the proliferation of CD4+ T cells, but CD8^+^ T cells remained unaffected (**Fig S15B-C**). In light of this observation, co-cultures were conducted after tumor-targeted depletion of NMD pathway components via siRNA-KD (**Fig. 6C**).

**Figure 6.**
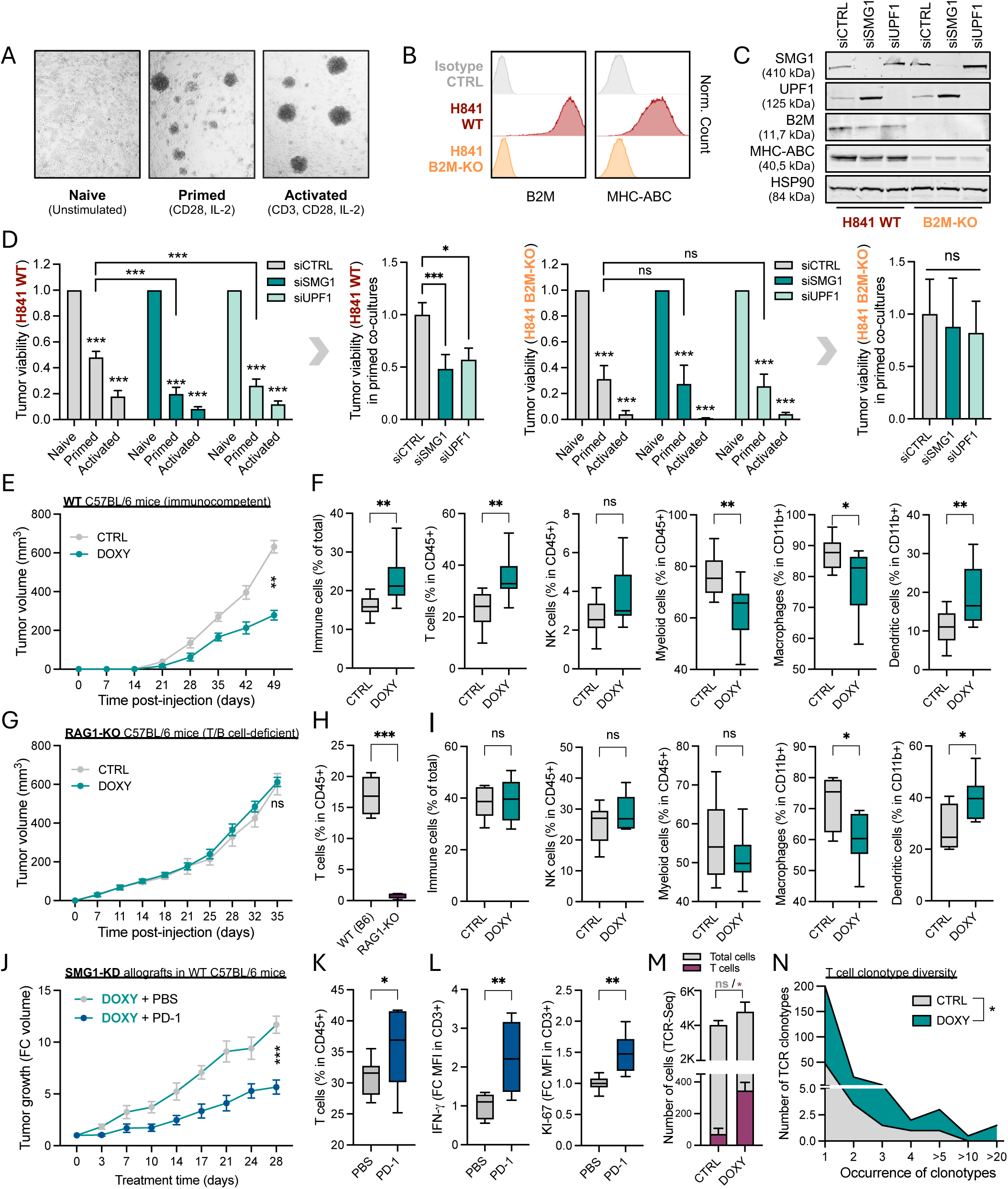
NMD inhibition enhances tumor immunogenicity (see also Fig. S15 and Table S5). **A-D)** *In vitro*co-cultures of the TMB^High^ human cell line H841 with PBMCs (>80% T cells, **Fig. S15D-E**). **(A)** Representative images 4 days post-stimulation with indicated stimuli promoting naive, primed or activated T cell status; **(B)** flow cytometry histograms showing B2M and MHC-ABC surface expression in WT *vs B2M*-KO H841 cells (**Fig. S15A**); **(C)** Western blot for WT and *B2M*-KO H841 cells showing SMG1-KD and UPF1-KD 72 hpt. **(D)** Tumor cell viability represented by the fraction of living single H841 cells (CD45-) measured by flow cytometry in 4 day-co-cultures with naive, primed or activated T cells. WT (**left**) or *B2M*-KO (**right**) H841 cells were transfected with siCTRL/siSMG1/siUPF1 and tumor killing under primed/activated conditions was normalized internally to the naive condition to subtract possible effects of NMD inhibition on cell-intrinsic viability. A subsequent re-normalization to the siCTRL condition is shown for primed co-cultures (n≥3). **E)** Tumor growth of RP1380 TetO-shSMG1 cell allografts growing in wild-type immunocompetent C57BL/6J mice fed with normal or doxycycline-diet (n=13). **F)** Flow cytometry immunophenotyping of endpoint tumors from **E** (n≥7) (see **Fig. S15F-G**). **G)** Tumor growth of RP1380 TetO-shSMG1 cells allografts growing in partially immunocompromised, T/B cell-deficient RAG1-KO C57BL/6J mice fed with normal or doxycycline-diet (n≥6). **H)** T cell quantification in WT C57BL/6J *vs* RAG1-KO C57BL/6J mice (n=4) (see **Fig. S15N**). **I)** Flow cytometry immunophenotyping of endpoint tumors from **G** (n≥6)**. J)** Tumor growth of RP1380 TetO-shSMG1 cell allografts grown in C57BL/6J mice fed with doxycycline-diet (SMG1-KD models) and treated with PBS or anti-PD-1 (see **Mehods**). **K-L)** Flow cytometry showing infiltrating T cell numbers (**K**) and activation markers (**L**) in endpoint tumors from **J** (n≥6). **M-N)** Single cell TCR sequencing of sorted CD45+ cells derived from digested enpoint tumors from **E**, displaying number of identified T cells within all CD45+ cells **(M)** and number and distribution of T cell clonotypes in control *vs* NMD-inhibited tumors **(N)** (n=2, see **Table S5H**). All statistics by unpaired t-tests (for **E**, **G** and **K** unpaired t-tests for AUC). For all graphs, error bars represent the SEM, and P-values are specified as ***P<0.001; **P <0.01; *P<0.05; ns=non-significant.

SMG1- and UPF1-KD in H841 tumor cells resulted in enhanced T cell-mediated tumor cell killing (*P* < 0.001) only in co-cultures with MHC-I-proficient H841 but not with H841 *B2M*-KO cells (**Fig. 6D**). Thus, NMD pathway inhibition not only created an altered repertoire of neoantigens (as shown in **Fig. 5**), possibly implicating a higher diversity of peptide/MHC combinations, but also led to an increase in MHC-I-dependent tumor cell killing. We note that NK cells (CD45+/CD56+), myeloid cells (CD45+/CD11b+) and B cells (CD45+/CD19+) were initially present in healthy PBMC donors (d-1 and d0). However, after 4 days of *in vitro* culture these populations markedly declined with NK cells decreasing to approximately 1% and T cells predominating, constituting >85% of all CD45+ cells (**Fig. S15D**). Consequently, no NK cells were detected in our co-cultures (**Fig. S15E**), excluding the possibility of NK cell-mediated cytotoxicity. Our results thus demonstrate that NMD inhibition enhances an MHC-I-dependent tumor cell recognition by T-cells *in vitro*, resulting in increased T cell cytotoxicity and tumor cell killing.

We next sought to evaluate tumor immunogenicity upon NMD inhibition *in vivo* using syngeneic (haplotype-matched) allografts of murine SCLC cell lines grown subcutaneously in immunocompetent C57BL/6 mice. In order to achieve tumor-specific targeting of the NMD pathway *in vivo,* we used the murine RP1380 TetO-shSMG1 cells (see again **Figs. 4J** and **S8C-J**). Since TMB^Low^ mSCLC RP cell lines did not reveal any cell-intrinsic lethality upon NMD inhibition (as shown *in vitro* in **Figs. 2-4** and *in vivo* upon engraftment in full immunocompromised NSG mice in **Fig. 4**), any decrease observed in tumor growth in immunocompetent C57BL/6 mice can be attributed to immune intervention. Despite its low TMB, we identified tumor-specific mutations in RP1380 cells and confirmed the upregulation of predicted neoantigens following SMG1i treatment (**Figs. S9-11**).

Employing the TMB^Low^ RP1380 TetO-shSMG1 allograft model, we observed a significant delay in tumor growth in immunocompetent C57BL/6 mice (WT) upon doxycycline-induced SMG1-KD (**Figs. 6E** and **S8G-J**) which was not evident in full immunocompromised NSG mice (as shown in **Fig. 4K**), thus demonstrating a relevant contribution of the immune system in controlling tumor growth upon tumor-targeted NMD inhibition. We quantified the levels of tumor infiltrates (**Fig. 6F**) and found a significant increase of CD45+ immune cells in SMG1-depleted tumors (DOXY), characterized by higher proportions of T cells (CD3+) and dendritic cells (CD11b+/CD14-; *P*<0.01), and a significant decrease in tumor-associated monocytes/macrophages (CD11b+/CD14+; *P*<0.05) which have well-known immunosuppressive roles^104–109^. On the contrary, mice engrafted with RP1380 TetO-shCTRL cells and subjected to a doxycycline diet exhibited neither altered tumor growth dynamics nor enhanced immune infiltration (**Fig. S15F-G**), ruling out immunogenic effects associated with the doxycycline treatment or the TetO system (see **Methods**). Our data therefore show that, while TMB^Low^ tumors such as mSCLC-RP models may upregulate a limited number of altered transcripts upon NMD inhibition, increased immune responses were nonetheless observed in our model system. This supports previous findings that even a single NMD-escape frameshift mutation can be sufficient to increase immune cell recognition^18^.

In order to further assess the role of T cells in controlling tumor growth upon NMD inhibition, we also engrafted RP1380 TetO-shSMG1 cells onto partially immunocompromised *RAG1*-KO C57BL/6 mice, which lack B and T cells, but not NK cells nor myeloid cells^110^. We observed no differences in tumor growth upon doxycycline-inducible NMD inhibition, thus further confirming a T cell-dependent control of tumor growth (**Figs. 6G-H**). These analyses also excluded any contribution of NK cells in controlling tumor growth. Notably, NMD inhibition-induced changes in dendritic cells and macrophages observed in WT C57BL/6 mice were also evident in *RAG1*-KO C57BL/6 mice (**Fig. 6I**), although their causes and contribution to immunogenicity remain to be investigated.

We further note that NMD inhibition did not alter levels of MHC-I surface expression on tumor cells *in vitro*, as neither siRNA-mediated SMG1/UPF1 knockdown nor doxycycline-inducible SMG1 knockdown *in vitro* resulted in enhanced MHC-I surface expression (**Fig. S15H-I**). This is in line with our MHC-I-immunopeptidome data which did not show changes in the overall number of peptides presented in control *versus* NMD inhibited samples (see again **Figs. 5** and **S14**). However, we observed *in vivo* increased MHC-I protein levels in NMD-inhibited tumors growing in fully immunocompetent wildtype C57BL/6 mice (**Fig. S15J**), which significantly correlated with the levels of immune cell infiltration (**Fig. S15K**). By contrast, this was not the case in NMD-inhibited tumors grown in *RAG1*-KO mice (**Figs. S14L-N**), thus indicating that increased MHC-I protein levels observed following NMD inhibition *in vivo* can be attributed to enhanced T cell infiltration. Collectively, our data show that NMD inhibition caused upregulation and presentation of neoantigens in cancer cells leading to enhanced immune cell infiltration and T cell activation in SCLC models; this in turn may modulate MHC-I surface expression on tumor cells *in vivo* (e.g. through IFNg secretion^111^) further boosting neoantigen presentation.

Since NMD inhibition enhanced neoantigen exposure in SCLC, we next sought to evaluate the therapeutic potential of combining NMD inhibition with immune checkpoint blockade (ICB), given the limited response to immunotherapy in SCLC patients. We previously showed that anti-PD-1 treatment alone does not reduce the growth of mSCLC RP tumors growing in C57BL/6 mice^80^. In contrast, combining anti-PD-1 treatment with NMD inhibition, compared with tumor-targeted NMD inhibition alone, resulted in further significant reduction in tumor growth (**Fig. 6J**), as well as enhanced tumor T cell infiltration (**Fig. 6K**) and activation (**Fig. 6L**). These results therefore confirm that NMD inhibition promotes T cell recognition and killing of tumor cells *in vivo*, and emphasize its potential to augment immunotherapy efficacy.

To further characterize T cell responses *in vivo*, we performed single cell TCR sequencing of sorted intra-tumoral CD45+ cells extracted from RP1380 TetO-shSMG1 allograft tumors growing in immunocompetent C57BL/6 mice (endpoint tumors from **Fig. 6E**). TCR sequencing again confirmed increased T cell numbers infiltrating NMD-inhibited tumors (**Fig. 6M**) and revealed a greater clonotype diversity and expansion among infiltrating T cells (**Fig. 6N**). These results indicate that increased tumor neoantigen diversity as a result of NMD inhibition (as shown in **Fig.5**) promotes the recruitment of T cells with broader TCR reactivities which may aid in eradicating heterogeneous tumors^112^.

Altogether, our work demonstrates that NMD inhibition in SCLC induces tumor cell-intrinsic apoptosis while additionally expanding both the expression and diversity of the neoantigen landscape in cancer cells, thereby reshaping tumors toward a more immunogenic state with increased DCs, fewer macrophages and elevated T cell infiltration and activation with a broader epitope repertoire - ultimately driving more effective tumor growth control and potentially enhancing the efficacy of immunotherapy.

## Discussion

Our work identifies a previously unrecognized and therapeutically tractable dependency of SCLC - and potentially other TMB^High^ cancers (e.g., as shown for tobacco-associated SMARCA4-deficient UT) - on the NMD pathway to mitigate mutation-induced proteotoxic stress and sustain tumor cell survival. While prior studies have linked the NMD pathway to limiting altered transcripts thereby shaping tumor immunogenicity^15,18,26–29^, we show for the first time that TMB^High^cancers such as SCLC exhibit a hyperactive NMD pathway. The mechanisms by which highly mutated cancer cells may upregulate NMD activity remain to be elucidated. Our work, however, demonstrates that the NMD pathway not only impacts the accumulation of aberrant transcripts to facilitate immune evasion, but also is critical for maintaining cellular homeostasis and thus supporting cell-intrinsic survival in hypermutated tumors. To our knowledge, this study is the first to reveal that NMD inhibition leads to both cell-intrinsic and immune-mediated anti-tumor effects, and to connect this dual therapeutic vulnerability to TMB.

Although the NMD pathway has attracted increasing attention in oncology in the recent years - particularly in cancer immunotherapy – our study is the first to systematically explore its therapeutic potential in SCLC, one of the most aggressive and heavily mutated tumor types. Specifically, we showed that NMD inhibition impairs SCLC cell proliferation and triggers ER stress-dependent cell-intrinsic apoptosis, effectively leading to a TMB-dependent control of cell survival *in vitro* and of tumor growth *in vivo*, even in the absence of an immune system.

Furthermore, we have shown that NMD suppression upregulates multiple neoantigen-encoding transcripts thereby altering the repertoire of presented neoantigens and enhancing T cell recognition of human and murine tumor cells *in vitro* and *in vivo*. Our data collectively confirm an immunogenicity of NMD-regulated mutations and demonstrate that NMD-dependent control of neoantigen presentation contributes to immune escape. Importantly, we showed that NMD-regulated frameshift peptides are remarkably immunogenic, which may have profound implications in the future design of neoantigen vaccines. Combining NMD inhibition with anti-PD-1 treatment further increased immunogenicity of SCLC models, as also recently shown in NSCLC^27,113^, thereby raising the prospect of improving the currently limited responses of SCLC patients to ICB treatment^7-9^. Despite previous reports suggesting low immunogenicity in SCLC due to reduced MHC-I expression^94–97^, our data support the presence of a functional antigen presentation machinery across multiple SCLC models. Notably, in tumors with impaired MHC-I expression or defects in the antigen presentation pathway, the immunogenic effects of NMD inhibition may be limited. However, in the context of sufficiently high TMB, such tumors may still benefit from the cell-intrinsic cytotoxic effects of NMD inhibition.

Our findings, based on both immunocompetent and immunodeficient mouse models, demonstrate that immune-mediated as well as cell-autonomous anti-tumor mechanisms are operative *in vivo*. Importantly, these observed effects were consistent across different methods of NMD perturbation and reproducible in multiple preclinical cellular and *in vivo* models. We anticipate that the combined effects of cell-autonomous mechanisms and enhanced immunogenicity can exert a robust control over tumor growth in SCLC and possibly other TMB^High^tumors. We are aware that TMB^High^ mSCLC RP-Msh2 cells would be the optimal model to investigate the extend of such combined effects *in vivo,* but unfortunately these cells did not grow as subcutaneous models in wildtype C57BL/6 mice due to immune intervention (data not shown). Our study warrants further investigation using autochthonous SCLC mouse models harboring TMB-low and TMB-high tumors, combined with conditional, tumor-specific targeting of NMD pathway components, to further elucidate the combined contribution of tumor cell-intrinsic death and T cell-mediated tumor killing in the overall control of tumor growth following NMD inhibition.

Our work importantly shifts the paradigm regarding the targetability of the NMD pathway. We show that the high number of somatic mutations is a liability for SCLC cell homeostasis. We confirmed that, upon NMD inhibition, intrinsic apoptosis in SCLC results from unresolved ER stress caused by the accumulation of misfolded proteins in this high TMB context. Accordingly, cell death induced by NMD inhibition significantly correlated with the samples’ TMB, with hypermutated mSCLC RP-Msh2 cells exhibiting greater sensitivity compared to murine TMB^Low^ mSCLC RP cells. Furthermore, non-neoplastic cells were more resistant to NMD inhibition than SCLC cells, thus suggesting a potential therapeutic index for NMD pathway inhibition in patients. These results should alleviate concerns of an unviable NMD inhibition-based therapy due to systemic toxicity. Although NMD is an essential mechanism ubiquitously expressed in all tissues and developmental stages, and total ablation of NMD components *in vivo* is often embryonically lethal^114–116^, NMD activity withstands fluctuations during stress and homeostasis^32,117,118^ as well as proliferation and differentiation^33,36,59,119^, and it is known to decline during aging^120^. Therefore, although sustained depletion of NMD may be universally lethal – as supported by publicly available DepMap data (www.depmap.org)^121^ showing that core NMD factors such as SMG1 and UPF1 are broadly essential across cancer cell lines (data not shown) –, healthy non-neoplastic cells are expected to tolerate therapeutic NMD off-cycles and be less sensitive to NMD inhibition than hypermutated cancer cells. Given the low number of somatic mutations in healthy cells, NMD inhibition should neither render these cells immunogenic, as confirmed in two recent parallel studies^27,113^, nor set them under unbearable ER stress, as shown by our study. In agreement with these arguments^27,28,113^ our data showed that pharmacological NMD inhibition *in vivo* results in reduced tumor growth without overt toxicity, thus demonstrating targetability of a pathway largely considered undruggable, in agreement with recent studies^27,28,113^.

Our work has thus uncovered a TMB-dependent dual vulnerability in SCLC and paves the way for the development of novel targeted NMD inhibition therapies with combined cell-intrinsic and immune-mediated anti-tumor effects. Despite the recent approval of the DLL3 T cell engager tarlatamab for treatment of chemotherapy-refractory disease^122^ and the ongoing clinical testing of several new targeted therapies^123,124^, SCLC remains one of the most deadly and hard-to-treat cancer types. Collectively our data suggest that NMD inhibition may represent a novel strategy for the treatment of patients with SCLC, and possibly other cancers with a high TMB, particularly when combined with immune checkpoint blockade.

## Acknowledgements

We thank all the patients in Cologne University Hospital who decided to support research in their most vulnerable moment, as well as their family and friends. We thank also the clinics personnel taking care of those patients and involved in sample collection. This work was supported by the German Research Foundation (DFG) through funding provided by a collaborative research center grant on small cell lung cancer (CRC1399, project-ID 413326622) to JG, RKT, MP, HCT, MC, FB, JB, SvK, KH, SK, AQ and HGR. JG, RKT, HCR, MP and SvK are supported by the German Federal Ministry of Education and Research (BMBF, e:Med consortium InCa, grant 01ZX1901A and 01ZX2201A). RKT is supported by the German state of North Rhine-Westphalia EFRE initiative (EFRE-0800397). Additional funding was received from "Netzwerke 2021", an initiative of the Ministry of Culture and Science of the German state of North Rhine-Westphalia for the CANTAR project to JG, RKT, SvK HCR and MP. Additional support has been received from the DFG (project ID: 497777992 to JG, MP and AQ; project ID: 418074181 to MP and HCR, and as part of the SFB1530 – project ID 455784452 to MP), the German Cancer Aid (Mildred-Scheel professorship to MP and project ID: 70116707 to FB), the Bruno-Helene-Joester foundation (JG and MP), and the Jean Uhrmacher Foundation (JG), the German Cancer Aid TACTIC grant (SK and HCR), the Mildred Scheel Nachwuchszentrum Grant (JB and FB, project-ID 70113307) by the German Cancer Aid (Deutsche Krebshilfe). We thank the Cystic Fibrosis Foundation for donating the SMG1 kinase inhibitor for our research. We appreciate plasmids donations of Prof. Fernando Pastor (University of Navarra, Spain) and Prof. Waldemar Kolanus (University of Bonn, Germany) and antibody donations, as well as advice on ER stress signaling from Dr. Christina M. Bebber, Prof. Elena Rugarli and Giada Di Pietro (University of Cologne, Germany) and Dr. Jan Eickhoff (Lead Discovery Center GmbH). We thank the FACS facility of the Max Planck Institute for Biology of Ageing (University Hospital of Cologne) for their contributions to this work, Eurofins-CEREP and Stephane Paris for the enzymatic phosphorylation assay on purified SMG1 kinase and the full kinome assay and KYINNO biotechnology for the pharmacokinetics study of KVS0001. We also thank Prof. Alessandro Annibaldi for the frequent use of his lab’s Incucyte device. We thank Dr. Anna Pavlova and Dr. Behnaz Pezeshkpoor from the Institute of Experimental Haematology and Transfusion Medicine (University of Bonn) for the MHC-I genotyping of our samples. We thank the Proteomics Core Facility at the German Cancer Research Center (DKFZ) for their excellent support and services. We thank the Cologne Center for Genomics as part of the West German Genome Center for support in processing sequencing data. We also thank the computing center of the University of Cologne (RRZK) for providing the CPU time on the DFG-funded supercomputer ’CHEOPS’, as well as for the constant support provided.

## Author Contributions

LATF designed and performed all experiments. RKT and JG supervised experimental design and performance. LATF, RKT and JG wrote the manuscript. VB and NG analyzed RNA sequencing data. JK and MP integrated SCLC data with TCGA genomic and expression data for NMD cancer score calculations. AR, LM and MP analyzed whole exome sequencing. JPB and ABR performed and analyzed MHC-I immunopeptidomics experiments. MGM and HAS assisted with FluoroSpot assays. CM assisted with sample preparation for exome, RNA and TCR sequencing. BB, NA, and LK assisted with mouse work. JB assisted with qPCR, TCGA data analysis and immunopeptidomics data analysis. LL assisted with proliferation and cell death assays. KW retrieved COSMIC data for all cancer types. FB and VS generated RP cells. FB analyzed GTEx data. OI and HCR generated the RPM mouse model. BG and VS assisted with generation of RPM cells. GB assisted with MTAs and ethical and safe handling of clinical specimens. SvK and AA advised on cell death assays, and together with FL assisted with RAG1-KO mouse work. KH, FJ and JW were the clinicians responsible for patient care and sampling. AQ and MS assisted with pathological characterization of tumor specimens. JDR assisted with the generation of H841 *B2M*-KO cells. MC, MS and JV analyzed TCR sequencing data. JV, PD and CG assisted on the identification of TAAs. NW advised on routinary flow cytometry experiments. SK advised on SMG1i inhibitor selectivity tests. AS, CG and HGR contributed with HLA-I and T cell expertise.

## Supplementary Figure Legends

**Figure S1 – related to Figure 1.**
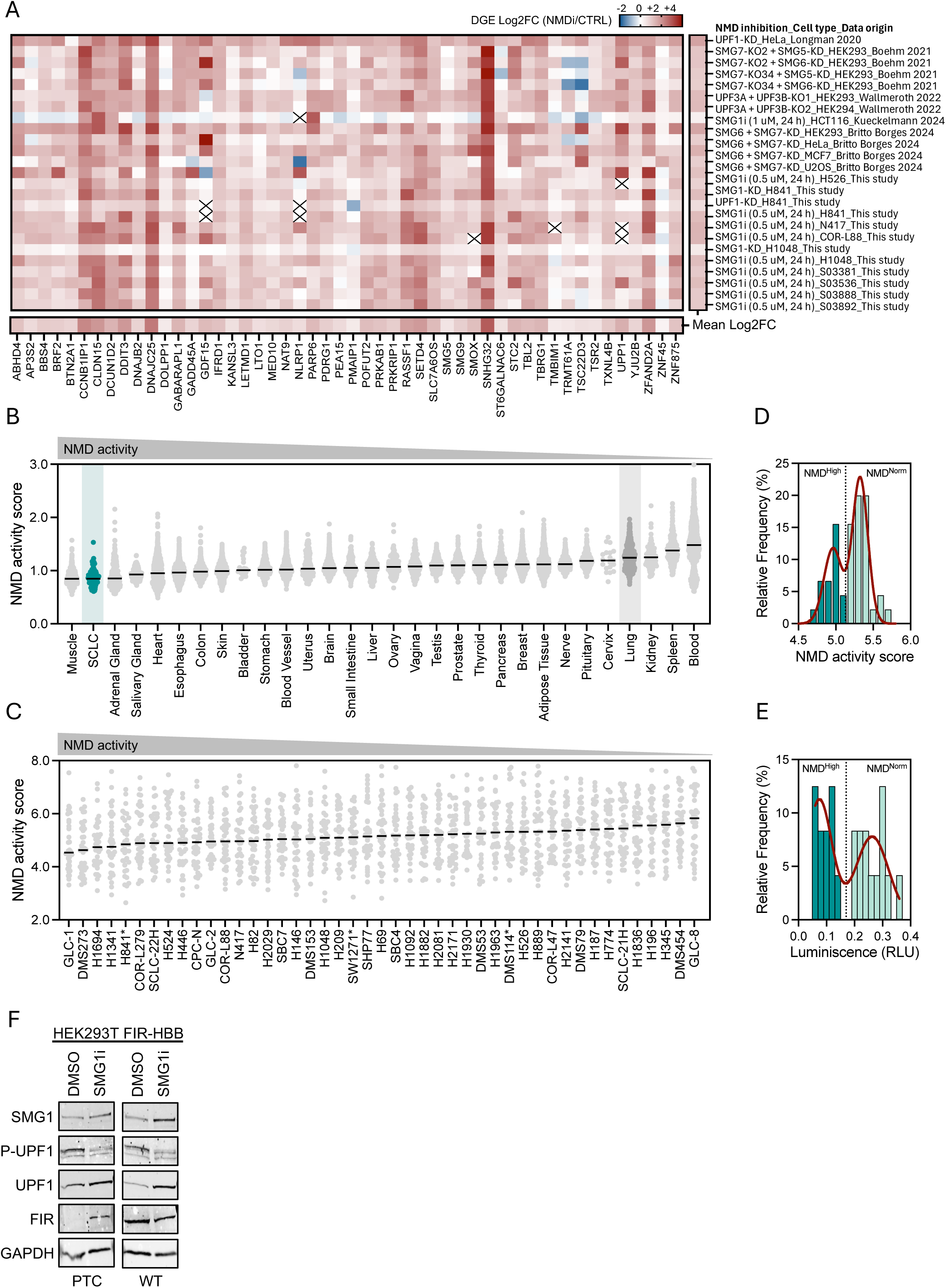
Transcriptome-based NMD activity scores reliably estimate NMD pathway activities across multiple human tissues and cancers. **A)** Heatmap for log2FC expression of NMD-inhibited (NMDi) *vs* control (CTRL) samples referring to 50 NMD-sensitive genes employed to calculate transcriptome-based NMD activity scores, as proposed by Wang et al^43^. Six independent NMD inhibition studies^48–53^ were included – different from the four studies previously employed by Wang et al.^44–47^ –, with the reference, cell system and NMD inhibition method indicated at the right side of the heatmap. This analysis additionally includes data from this study referring to NMD inhibition by SMG1i treatment in SCLC cell lines (n=5), patient-derived tumor cells (n=4), and SMG1-KD or UPF1-KD in the SCLC cell line H1048 and the SMARCA4-UT cell line H841. **B)** Transcriptome-based NMD activity scores calculated for all adult healthy tissues within the GTEx database harmonized with published SCLC data^5^, with SCLC and lung tissue highlighted. **C)** Transcriptome-based NMD activity scores calculated from expression data for n = 45 human SCLC cell lines, some of which have been recently reclassified as SMARCA4-UT^63^ (marked by *, see **Table S1C**). **D)** Frequency distribution for NMD activity score median values shown in **C**, with resultant histograms adjusting to two gaussian curves which reveal two different NMD activity groups. **E)** Frequency distribution for Luminescence (RLU) values derived from luciferase reporter assays (see Fig. 1E), with resultant histograms adjusting to two gaussian curves which reveal two different NMD activity groups. **F)** Western blot of HEK293T treated with DMSO vs SMG1i and transfected for luciferase assays shown in Fig. 1D with FIR-HBB-WT (WT) or FIR-HBB-PTC reporters (PTC). **G)** Somatic mutation load and transcriptome-based NMD activity scores calculated for all cancer cell lines within the CCLE database, supporting a high NMD activity in TMB-High tumors.

**Figure S2 – related to Figure 2.**
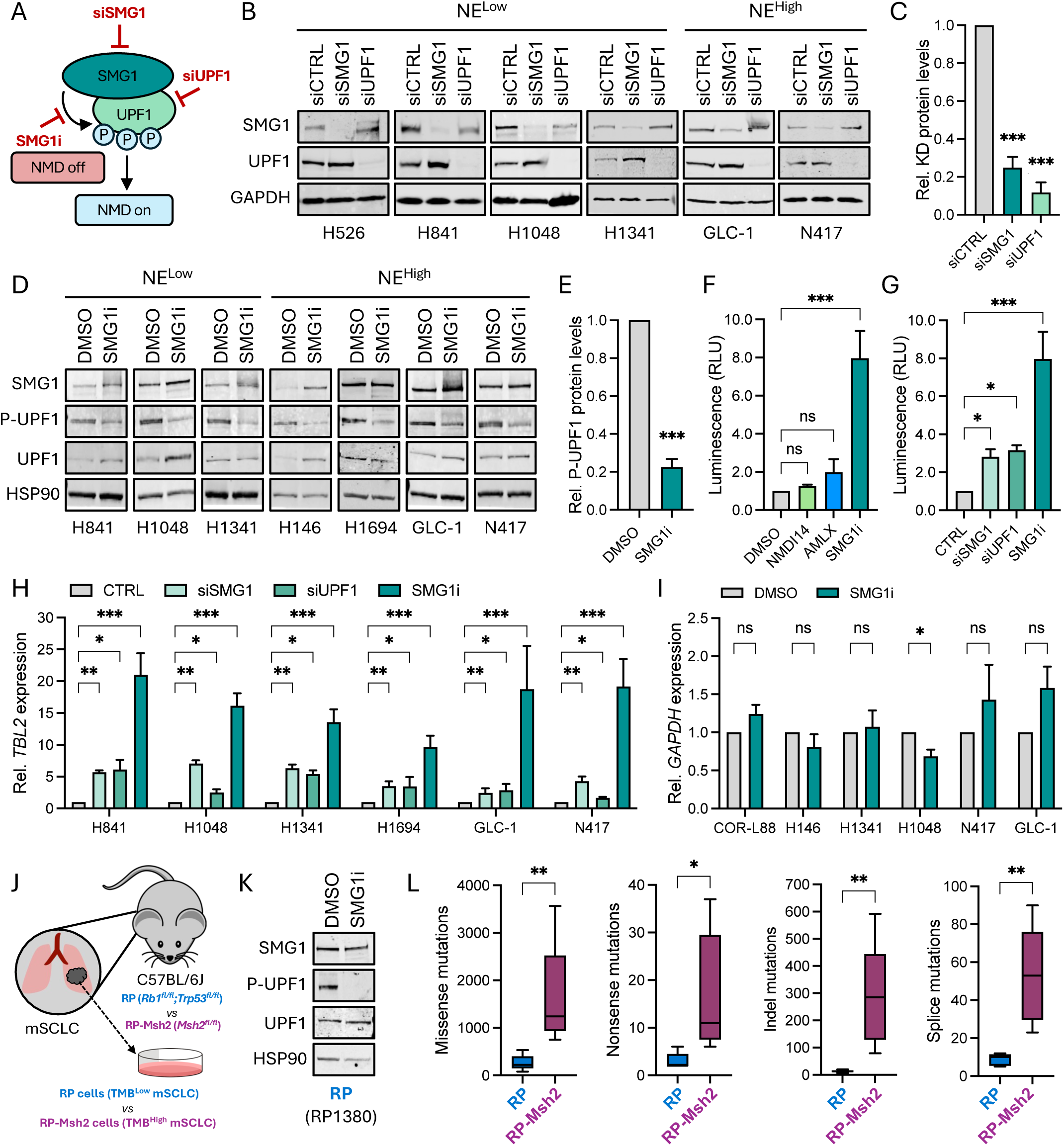
Transient inhibition of the NMD pathway through siRNA mediated knockdown and chemical inhibition. **A)** Schematic representation of the NMD pathway and approaches for NMD inhibition by siRNA-mediated knockdown (KD) of *SMG1* (siSMG1) or *UPF1* (siUPF1), or chemical inhibition of the SMG1 kinase (SMG1i). **B)** Representative western blot for n=4 SCLC cell lines and the SMARCA4-undifferentiated (UT) cell line H841^125^ and the small cell carcinoma of the cervix cell line H1341^125^ 72 hours post-transfection (hpt) with siCTRL/siSMG1/siUPF1. Neuroendocrine phenotypes (NE^High^ and NE^Low^) are indicated. **C)** Band densitometry quantification of western blots depicted in panel B for the SMG1-KD and UPF1-KD protein levels (relative to GAPDH) in siSMG1- and siUPF1-transfected cells, respectively. **D)** Representative western blot for n=5 SCLC cell lines, the SMARCA4-undifferentiated (UT) cell line H841^125^ and the small cell carcinoma of the cervix cell line H1341^125^ 24 hours post-treatment with 0.5 µM SMG1i. NE^High^ and NE^Low^ phenotypes are indicated. **E)** Band densitometry quantification of western blots depicted in panel D for the levels of P-UPF1 (relative to UPF1/HSP90). **F-G)** NMD inhibition quantified by dual luciferase reporter assays comparing SMG1i treatment (0.5 µM, 24 h) with **(F)** previously reported NMD inhibitors NMD14^69^ (50 µM, 48 h) and Amlexanox^68^ (AMLX, 100 µM, 48 h) (n = 3), and **(G)** with siRNA-mediated KD of SMG1 and UPF1 (n = 4). **H-I**) qPCR analysis in SCLC and H841 cancer cell lines following NMD inhibition through siSMG1/siUPF1/SMG1i treatment quantifying levels of (**H**) the *bona fide* NMD target *TBL2* (n ≥ 3) and **I)** a non-target of NMD, *GAPDH*. **J)** Schematic representation for mSCLC cell lines derived from lung tumors arising in RP (*Rb1^fl/fl^;Trp53^fl/fl^*, TMB^low^) and RP-Msh2 mice (*Rb1^fl/fl^;Trp53^fl/fl^;Msh2^fl/fl^,* TMB^high^) following Adeno-CMV-Cre inhalation. **K)** Western blot showing the SMG1i-mediated inhibition of UPF1 phosphorylation in a representative murine cell line. **L)** Comparison of exome mutations in RP *vs* RP-Msh2 cell lines. Bars represent mean + SEM. ***P < 0.001; **P < 0.01; *P < 0.05; ns = non-significant (One-sample t-test for C and E; One-way ANOVA for F and G, Two-way ANOVA for H; unpaired t-tests for I).

**Figure S3 – related to Figure 2.**
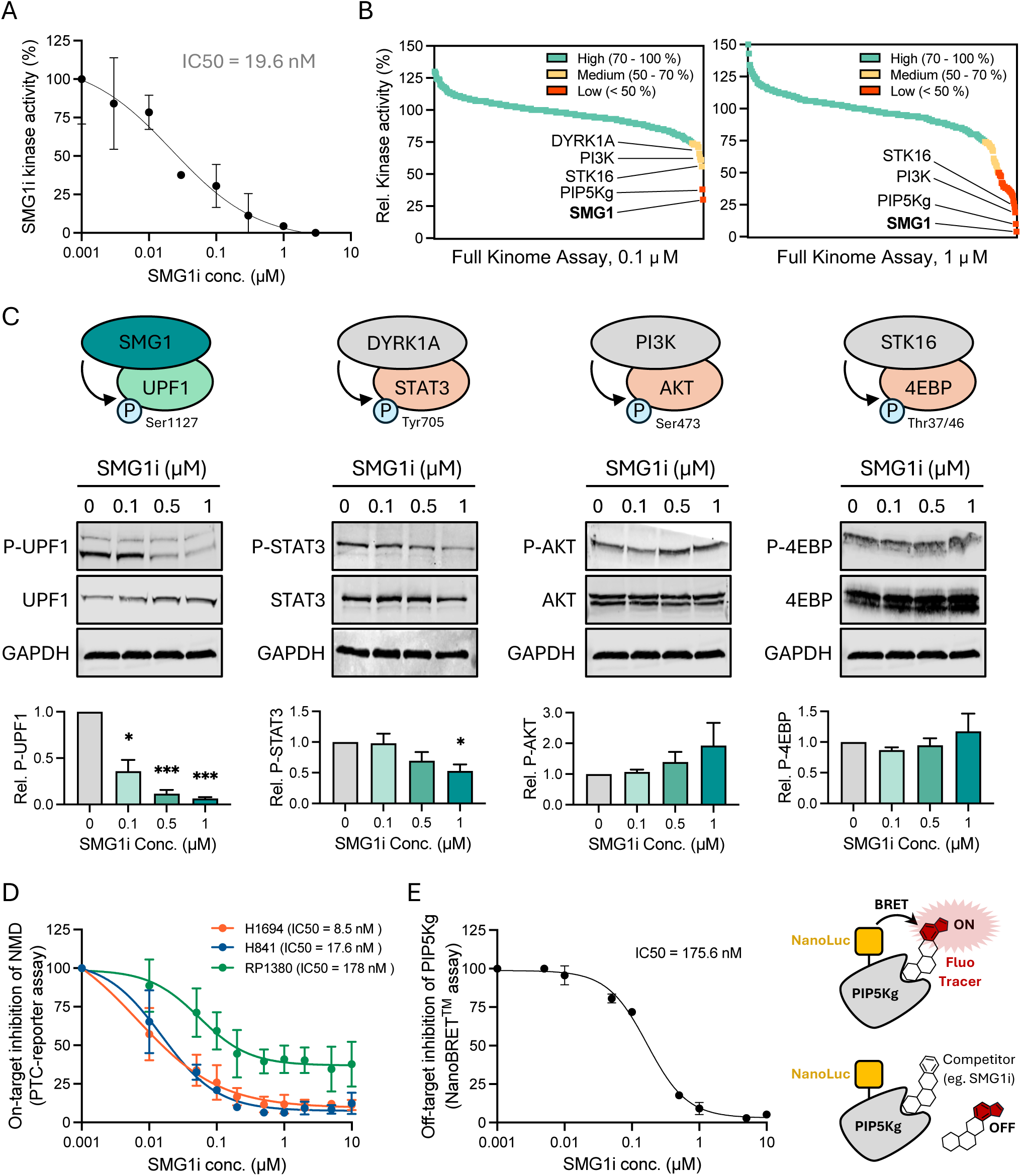
The SMG1i compound 11j is a potent and highly selective SMG1 kinase inhibitor. **A)** Enzymatic *in vitro* phosphorylation assay for full-length recombinant SMG1 protein using p53-Ser15 as a phosphorylation substrate in the presence of increasing SMG1i concentrations. IC50 was calculated by non-linear adjustment of the kinase activity inhibitory curve. **B-C)** Kinome assay on 436 kinases for selected SMG1i concentrations showing the relative kinase activity (%) for each individual kinase upon NMD inhibition with 100 nM **(B)** and 1000 nM SMG1i **(C)**, with kinases showing medium (yellow) to low (red) activity highlighted (see **Table S2A**). **D)** Western blot evaluation of the off-target inhibition of selected kinases in COR-L88 cells (NE^High^ SCLC) treated with increasing concentrations of SMG1i, assessing for the respective kinase activity by probing for the phosphorylated downstream substrate as indicated in the scheme (upper panel). Quantifications based on band densitometry are provided (lower panels). Bars represent mean + SEM (n = 3); ***P<0.001; *P<0.05 (Unpaired t-tests *vs* untreated sample). **E)** On-target inhibition of NMD by SMG1i as shown by cell-based luciferase reporter assays in human SCLC (H1694), human SMARCA4-UT (H841) and murine SCLC (RP1380) cell lines transfected with the NMD-sensitive PTC-reporter and treated with increasing SMG1i concentrations. IC50 values were calculated by non-linear adjustment of NMD inhibitory curves (n = 3). **F)** Off-target inhibition of PIP5Kg by SMG1i as shown by NanoBRET assays, with IC50 value calculated by non-linear adjustment of PIP5Kg inhibitory curve, accompanied by a schematic representation of the NanoBRET assay principle.

**Figure S4 – related to Figure 2.**
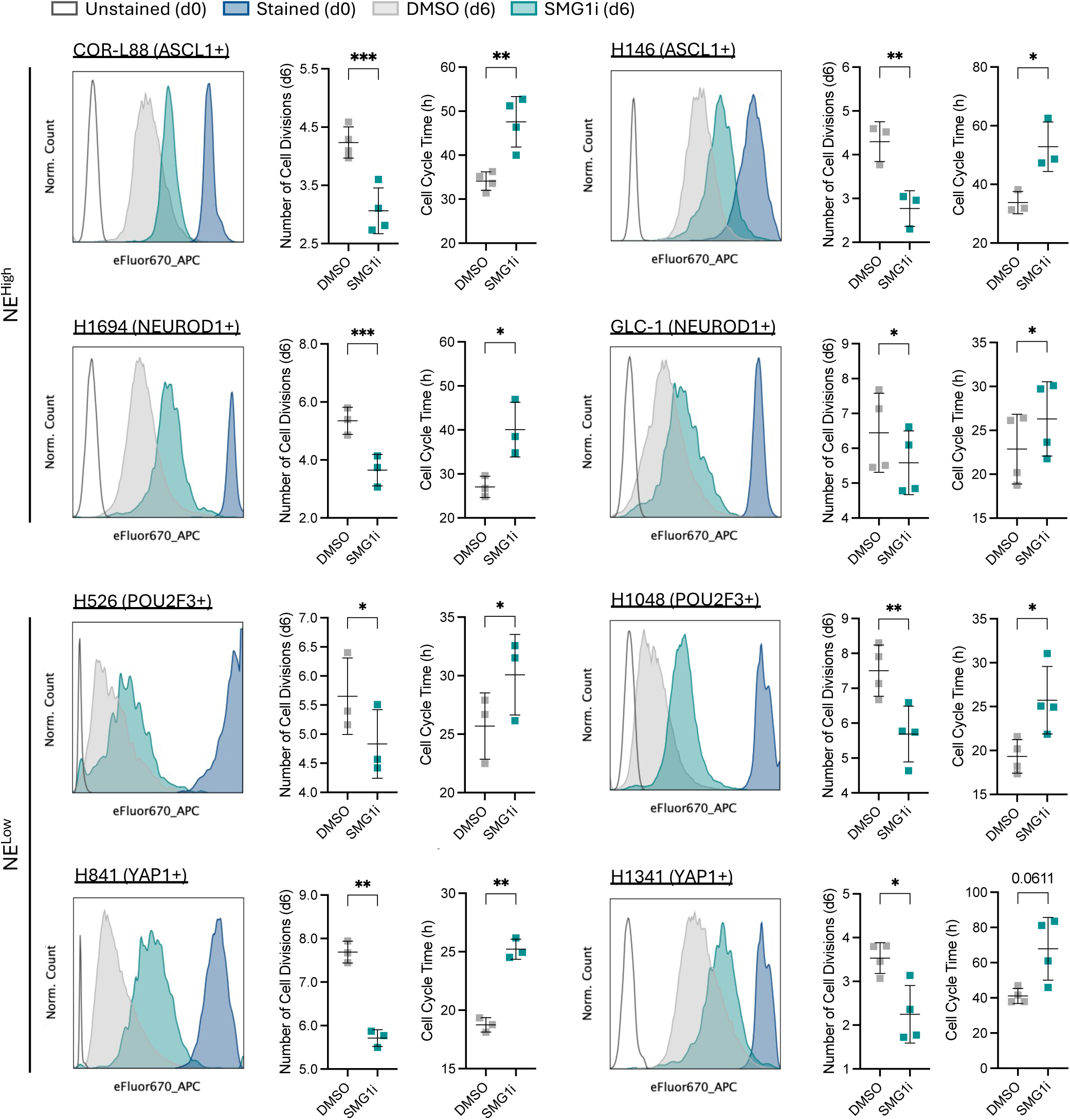
Inhibition of NMD restricts cell proliferation in SCLC. **A)** Proliferation assays for n=8 human SCLC cell lines (two of each SCLC subtype: COR-L88 and H146 as ASCL1+; N417 and GLC-1 as NEUROD1+; H526 and H1048 as POU2F3+; H841 and H1341 as YAP1+, with H841 recently reclassified as a SMARCA4-UT cell line). Neuroendocrine phenotype is indicated (NE^High^ and NE^Low^). For each cell line, a representative flow cytometry eFluor670 histogram for DMSO *vs* SMG1i-treated cells 6 days post-staining/treatment is shown (left), along with a quantification of the number of cell divisions after 6 days based on the loss of eFluor670 fluorescence intensity over time (middle) and the estimated average duration of cell cycle time in hours (right).

**Figure S5 – related to Figure 2.**
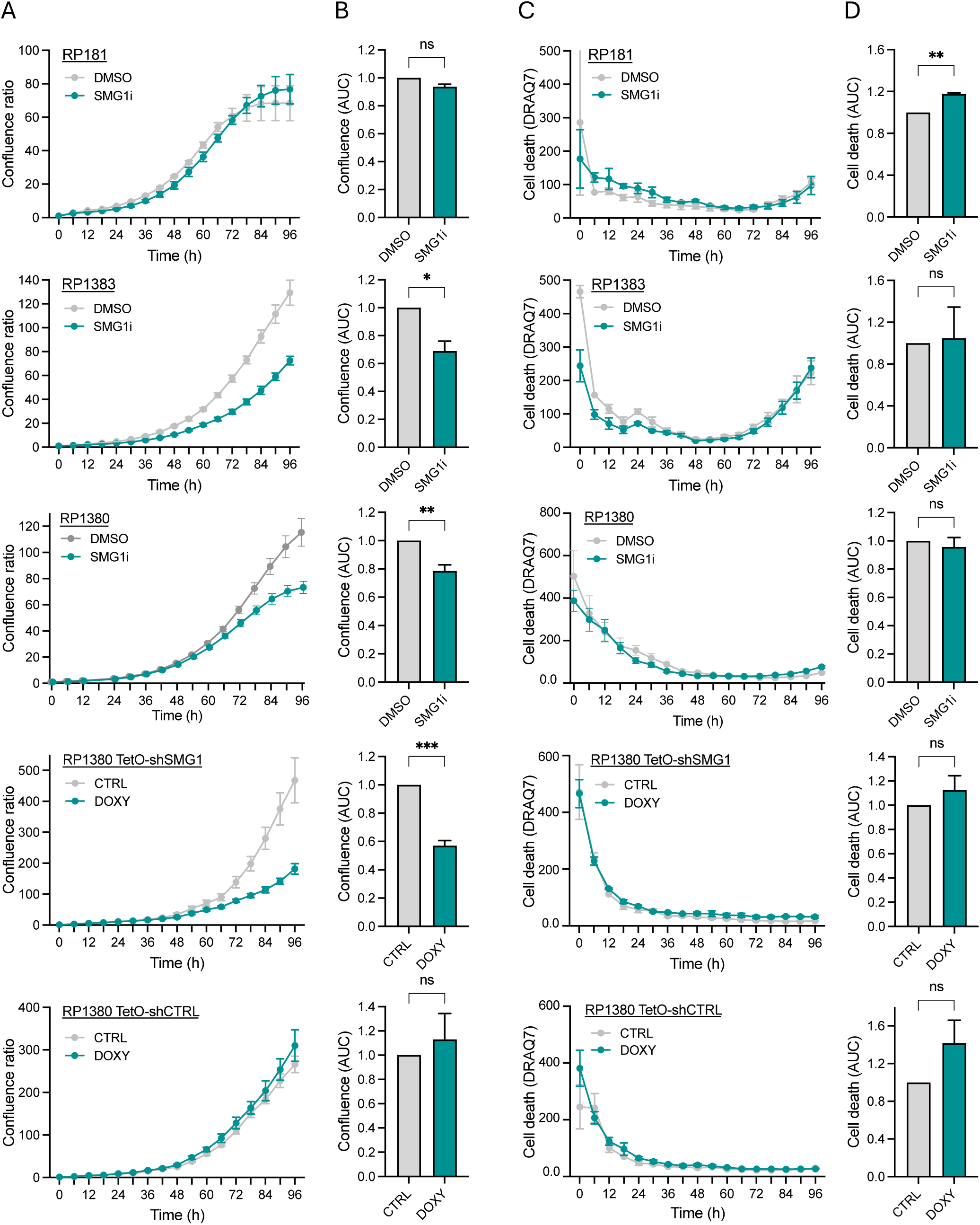
Inhibition of NMD does not induce cell death in murine TMB^low^ SCLC cell lines. **A-D)** Representative curves for cell growth (confluence ratio, **A**) and cell death (DRAQ7 staining, **C**) determined by live-cell imaging accompanied by respective AUC quantifications (**B** and **D**, respectively) for individual experiments (n = 3) in several murine TMB^low^SCLC cell lines. All graphs represent mean + SEM. ***P < 0.001; **P < 0.01; *P < 0.05; ns = non-significant (Unpaired t-tests on AUC quantification). The three upper panels refer to chemical NMD inhibition treatment with DMSO *vs* SMG1i, and the two lower panels refer to DMSO vs doxycycline (DOXY) treatment of RP1380 TetO-shSMG1 cells for genetic NMD inhibition via SMG1-KD, and in RP1380 TetO-shCTRL cells as a control to account for doxycycline-associated effects. Doxycycline-inducible SMG1-KD cellular models are described in **Fig. S8C-E**.

**Figure S6 – related to Figure 2.**
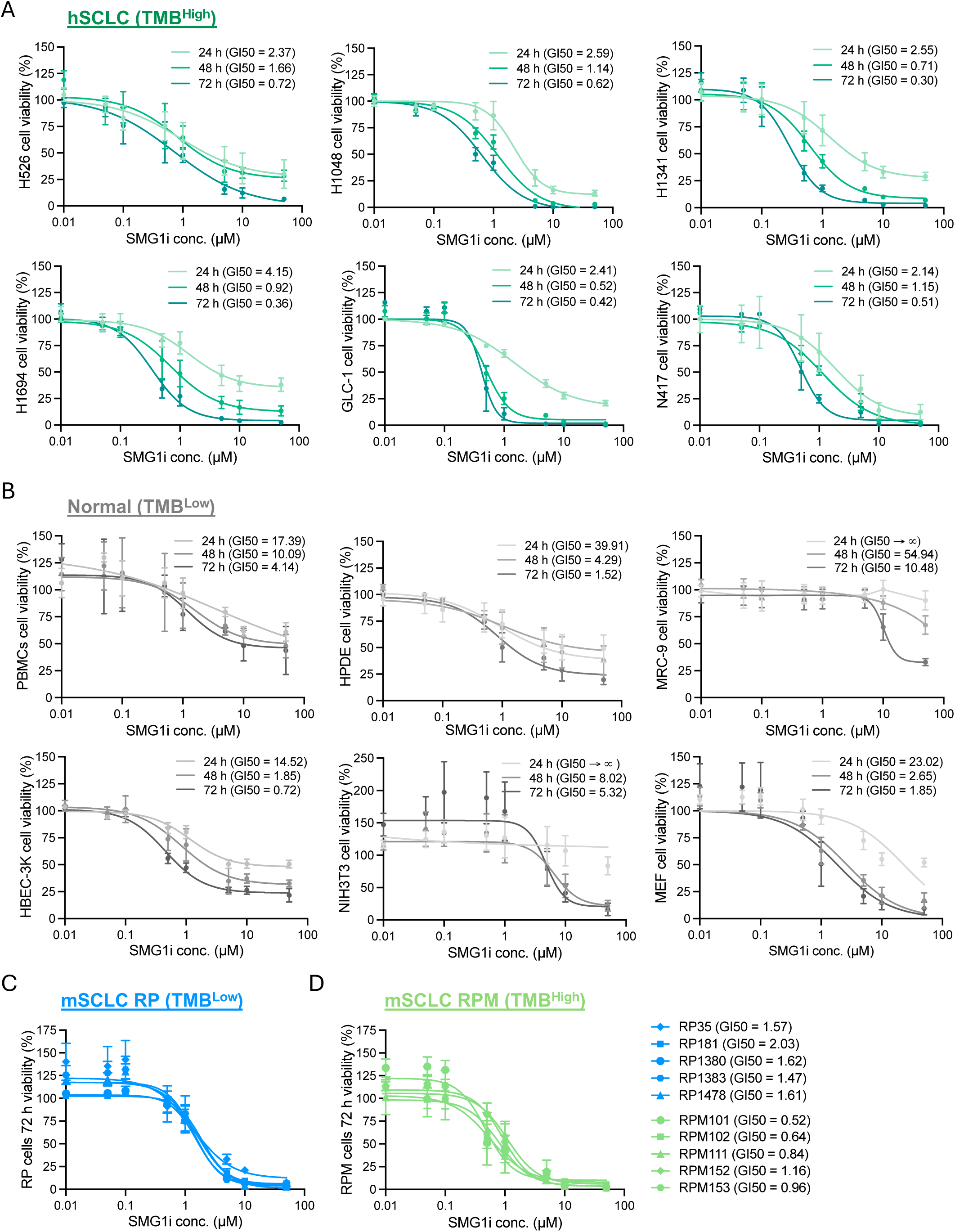
Cell-intrinsic vulnerability to NMD inhibition correlates with mutational load implicating a therapeutic index. CTG-viability assays following treatment at the indicated times with increasing concentrations of SMG1i for (**A**) the indicated human TMB^High^ SCLC cell lines, (**B**) TMB^Low^ non-neoplastic (normal/healthy) cells, including PMBCs = peripheral blood mononuclear cells (primary); HPDE = human pancreatic duct epithelial cell line; MRC-9 = lung human fibroblast cell line; HBEC-3K = human bronchial epithelial cells; MEFs = mouse embryonic fibroblasts; NIH3T3= mouse embryonic fibroblasts; and (**C**) TMB^Low^ murine SCLC RP (*Rb1-/-; Trp53-/-*) primary cells and (**D**) TMB^High^ murine SCLC RPM (*Rb1-/-; Trp53-/-; Msh2-/-*) primary cells. The growth inhibitory concentration GI50 was estimated from non-linear adjustments of viability curves. Graphs represent mean + SEM (n ≥ 3).

**Figure S7 – related to Figure 3.**
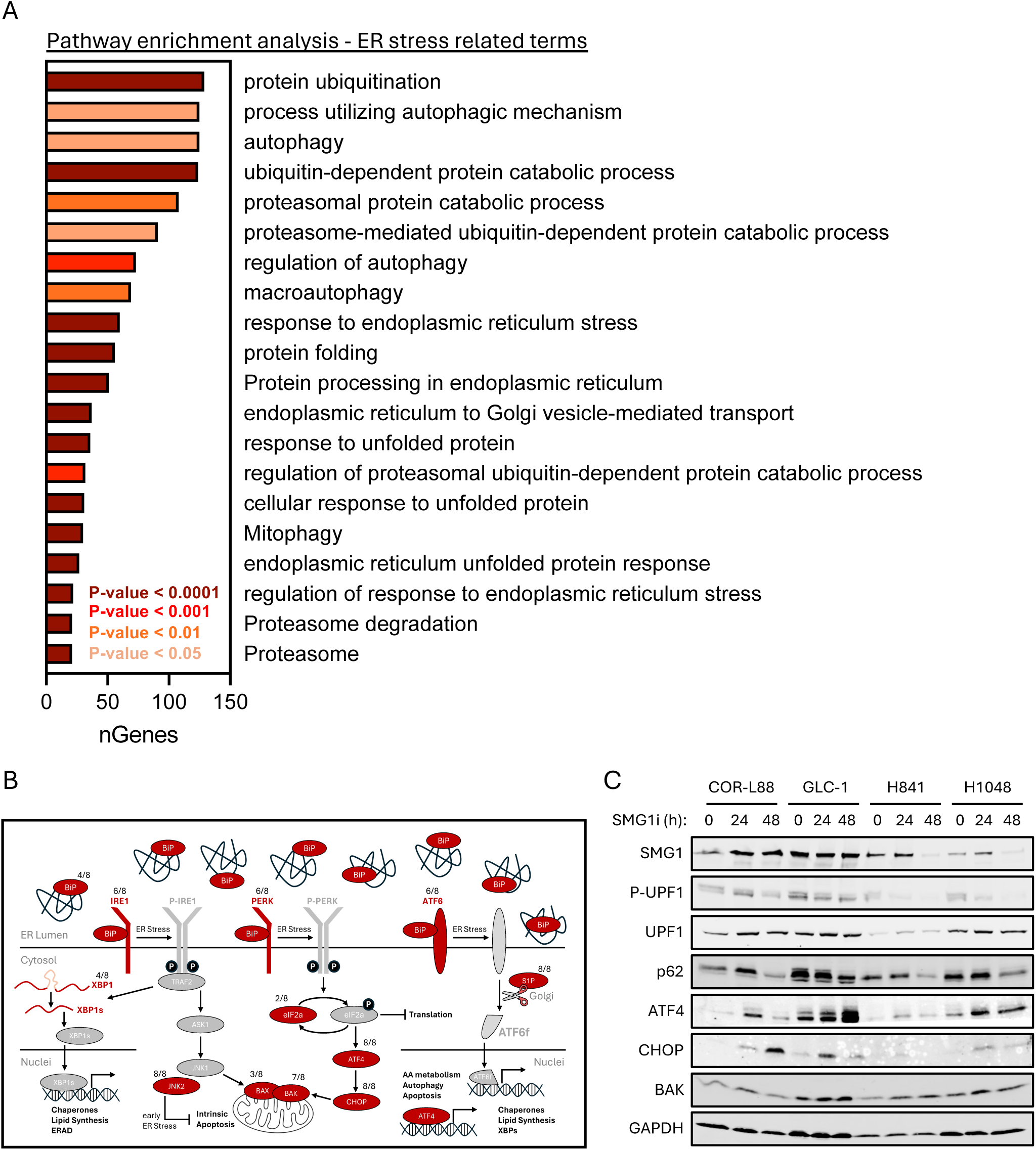
NMD inhibition triggers ER stress and upregulates the unfolded protein response (UPR) pathway. **A)** Pathway enrichment analysis based on transcriptome sequencing data of n=8 SCLC samples treated with DMSO *vs* SMG1i (See **Tables S3** and **S4**). Transcripts commonly upregulated (Log2FC > 1 and Padj < 0.05) upon SMG1i treatment in at least 6 out of 8 SCLC samples revealed an enrichment of multiple ER stress-related terms. **B)** Schematic representation of the ER stress response pathway also known as the unfolded protein response (UPR) pathway highlighting in red those proteins for which transcripts were found significantly upregulated in at least n=2 SCLC samples. The number of samples for which such an upregulation was observed is indicated as a fraction (e.g. 2/8) next to the respective protein. **C)** Representative western blot showing levels of selected proteins of the UPR pathway (apoptosis effector ATF4-CHOP-BAK1 axis) and the autophagy substrate p62 upon SMG1i treatment in SCLC cell lines, including one cell line per SCLC subtype (COR-L88 ASCL1+, GLC-1 as NEUROD1+, H1048 as POU2F3+ and H841 as YAP1+ which has been reclassified as SMARCA4-UT cell line).

**Figure S8 – related to Figure 4.**
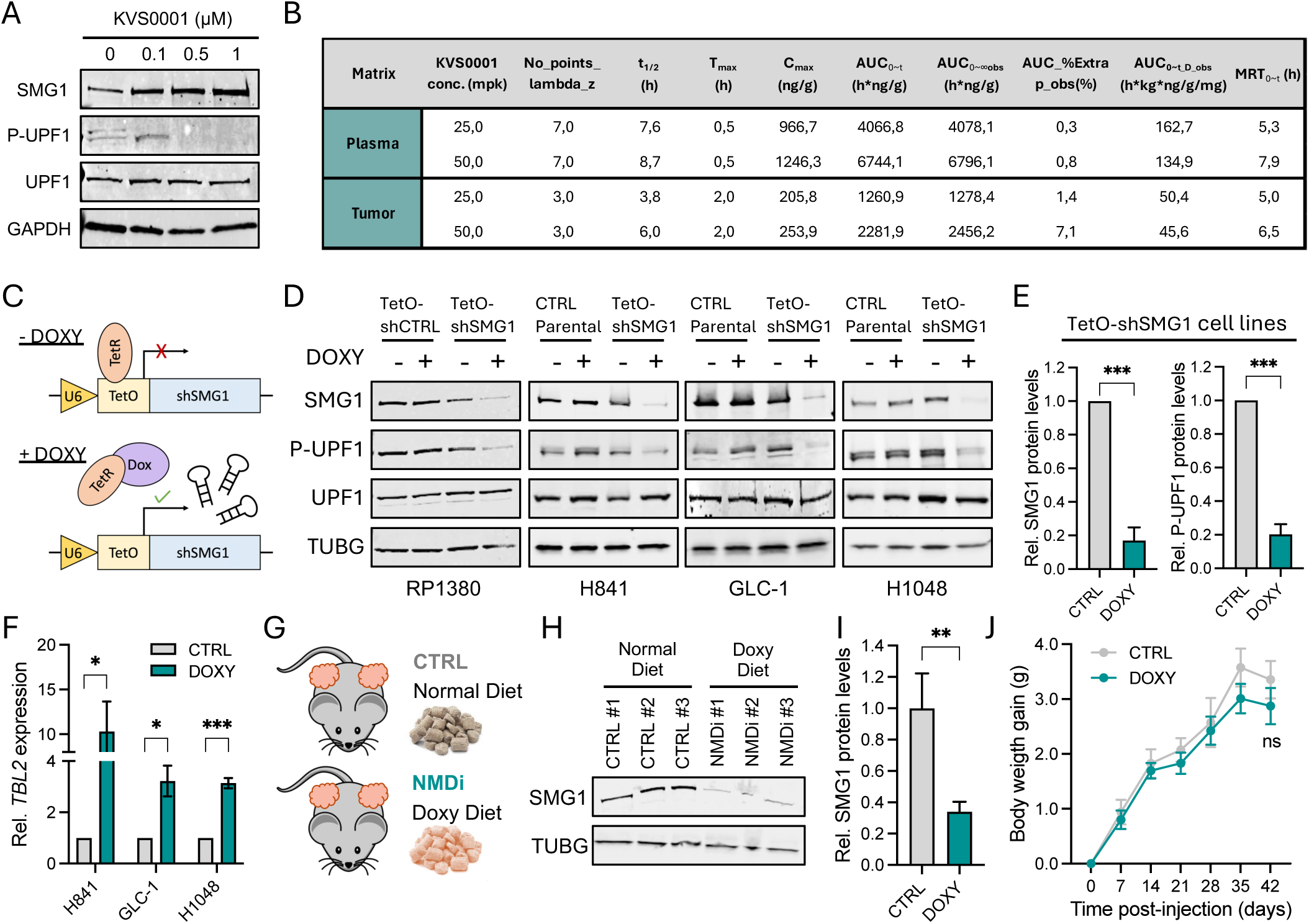
Approaches for *in vivo* inhibition of NMD: pharmacological systemic inhibition via KVS0001 treatment and genetic tumor-targeted inhibition via doxycycline-inducible SMG1-KD. **A)** Western blot on the human COR-L88 SCLC cell line (NE^High^, ASCL1+) treated *in vitro* with increasing concentrations of the SMG1i-derivate compound KVS0001 for 24 h. **B)** Pharmacokinetic parameters for KVS0001 compound detected by mass spectrometry in plasma and tumors of NSG mice after a single intraperitoneal injection with 25 or 50 milligram per kg (mpk) at timepoints post-injection indicated in **Fig. 4C-D. C)** Schematic representation for the doxycycline-inducible SMG1-KD operating within TetO-shSMG1 models, which include the murine SCLC cell line RP1380, the human SMARCA-UT cell line H841, the human NE^High^ SCLC cell line GLC-1 and the human NE^Low^ SCLC cell line H1048. **D)** Western blots for TetO-shSMG1 cell lines probing for downregulation of SMG1 and consequently decreased P-UPF1 levels upon adding 2 ug/ml doxycycline to the cell culture for 72h. For human TetO-shSMG1 cell lines, the corresponding parental cell lines served as control to account for the effects of doxycycline treatment. For the murine SCLC cell line RP1380, a TetO-shCTRL cell line was generated to exclude immunogenic effects of the TetO system in subsequent experiments shown in Fig. 6. **E)** Band densitometry quantification for SMG1 (relative to TUBG) and P-UPF1 (relative to UPF1/TUBG) protein levels in TetO-shSMG1 cell lines referring to the data in panel **D**. **F)** qPCR quantification of the *bona fide* NMD target *TBL2* in human TetO-shSMG1 cell lines after 72h of *in vitro* doxycycline treatment. **G)** Scheme for *in vivo* induction of SMG1-KD through the administration of doxycycline-containing diet to either NSG (Fig. 4) or C57BL/6J (Fig. 6) mice subcutaneously implanted with TetO-shSMG1 cells. **H)** Representative blot for SMG1 protein levels from allograft RP1380 TetO-shSMG1 subcutaneous tumors derived from C57BL/6J mice fed with normal or doxycycline diet. **I)** Densitometry quantification of protein levels determined in western blots for data in panel H (n ≥ 14 tumors grown *in vivo*). **I)** Mouse weight changes over time in mixed NSG and C57BL/6J mice receiving normal or doxycycline diet (n = 16).

**Figure S9 – related to Figure 5.**
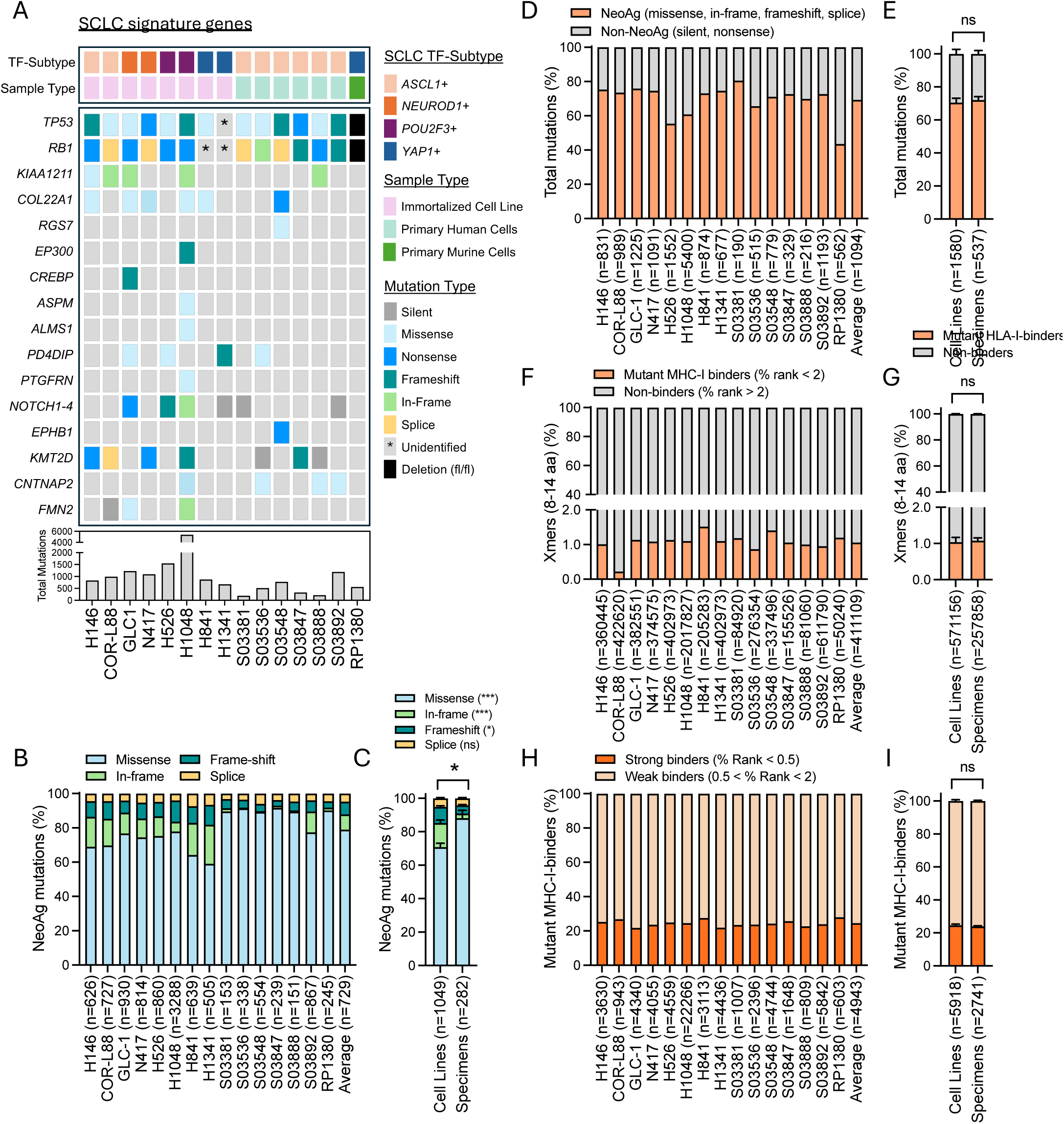
Whole exome sequencing followed by MHC-I-binding modeling predicts thousands of neoantigens in SCLC samples. **A)** SCLC-relevant genome alterations^3,5^ determined for the indicated SCLC cell lines and patient-derived tumor specimens by whole exome sequencing processed with our in-house pipeline. **B-C)** Proportion of potentially immunogenic neoantigen (NeoAg) mutations (i. e. missense, in-frame, frameshift and splice) for the indicated SCLC samples (**B**) and comparing cell line *vs* patient data (**C**). Nonstop mutations are included in the frameshift mutation count, and intron-exon mutations are included in the splice mutation count. **D-E)** Proportion of NeoAg and non-NeoAg mutations (silent and nonsense) for the indicated SCLC samples (**D**) and comparing cell line *vs* patient data (**E**). **F-G)** Proportion of MHC-I binders and non-binders predicted within all tested overlapping Xmers (8mers-14mers) for the indicated SCLC samples (**F**) and comparing cell line *vs* patient data (**G**). **H-I)** Proportion of strong and weak MHC-I-binders for the indicated SCLC samples (**H**) and comparing cell line *vs* patient data (**I**). Every graph indicates proportion (%), with absolute numbers (n = abs) indicated for each sample. Average values (n = abs_mean) are also indicated for all samples, for all cell lines and for all patient specimens. Error bars represent the SEM. ***P-value < 0.005; *P-value < 0.05; ns = non-significant (Unpaired t-tests:). See also **Table S5A-D.**

**Figure S10 – related to Figure 5.**
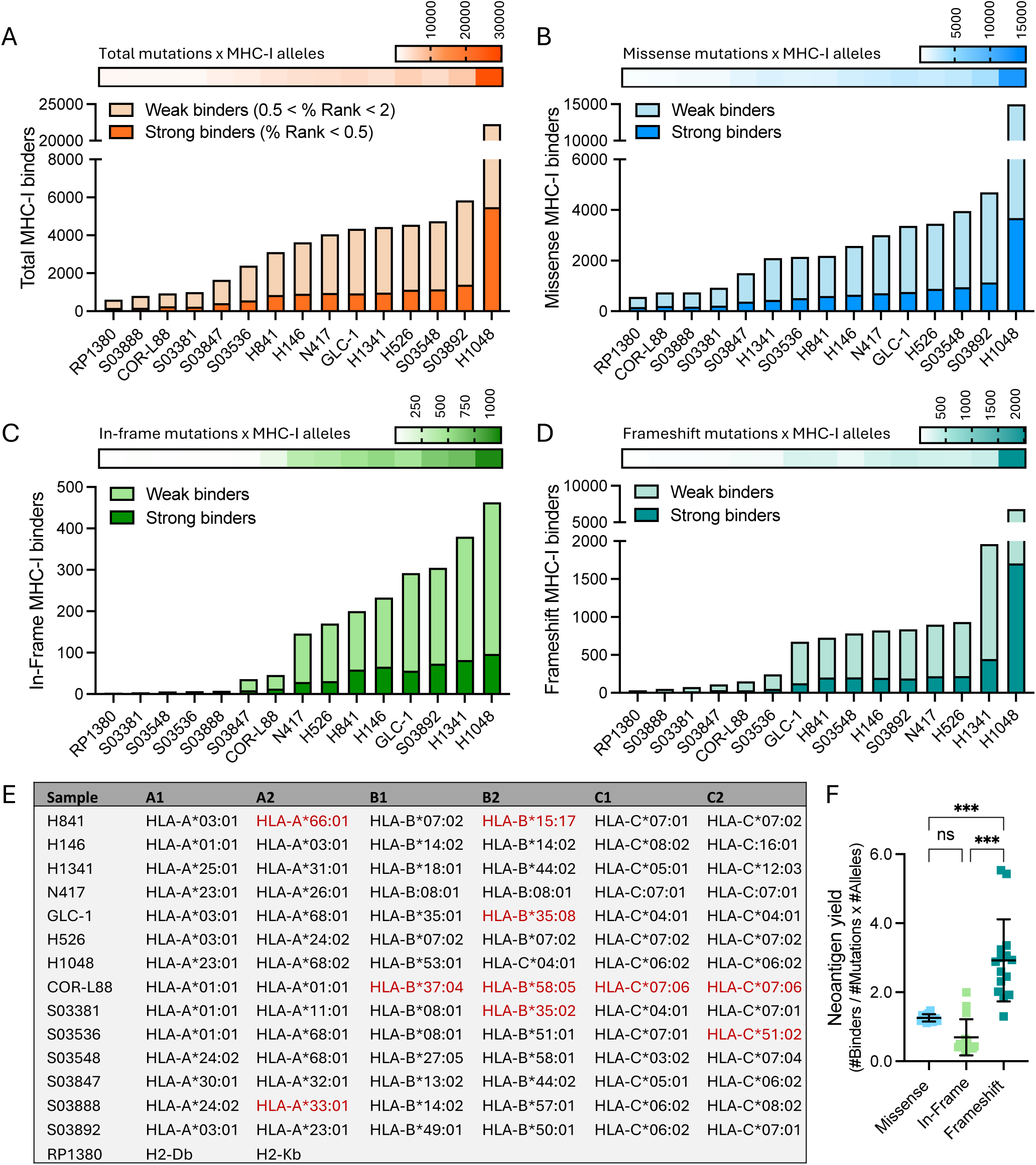
Mutation type-specific neoantigen prediction reveals a higher neoantigen yield of frameshift mutations. **A-D)** Number of predicted MHC-I binders identified for the indicated SCLC samples for all missense, in-frame and frameshift mutations together (**A**), or independently for missense (**B**), in-frame (**C**) and frameshift mutations (**D**). The number of MHC-I-binders identified in each case highly depends on the number of input mutations and the number of input MHC-I alleles (shown in **E**) included for the prediction, as shown by the upper heatmap panels in A-D. E) *MHC-I* locus genotyping for all samples included in the MHC-I binding prediction analysis. Alleles marked in red were not available for selection in the IEDB online *MHC-I binding tool* and therefore binders for those alleles could not be predicted. **F)** Neoantigen yield of different mutation types, which considers the number of predicted neoantigens (MHC-I-binders) per mutation and per MHC-I allele included in the prediction. Error bars represent the SD. ***P-value < 0.005; **P-value < 0.01; *P-value < 0.05 (One-way ANOVA). See also Fig. 5A**-B** and **Table S5A-D.**

**Figure S11 – related to Figure 5.**
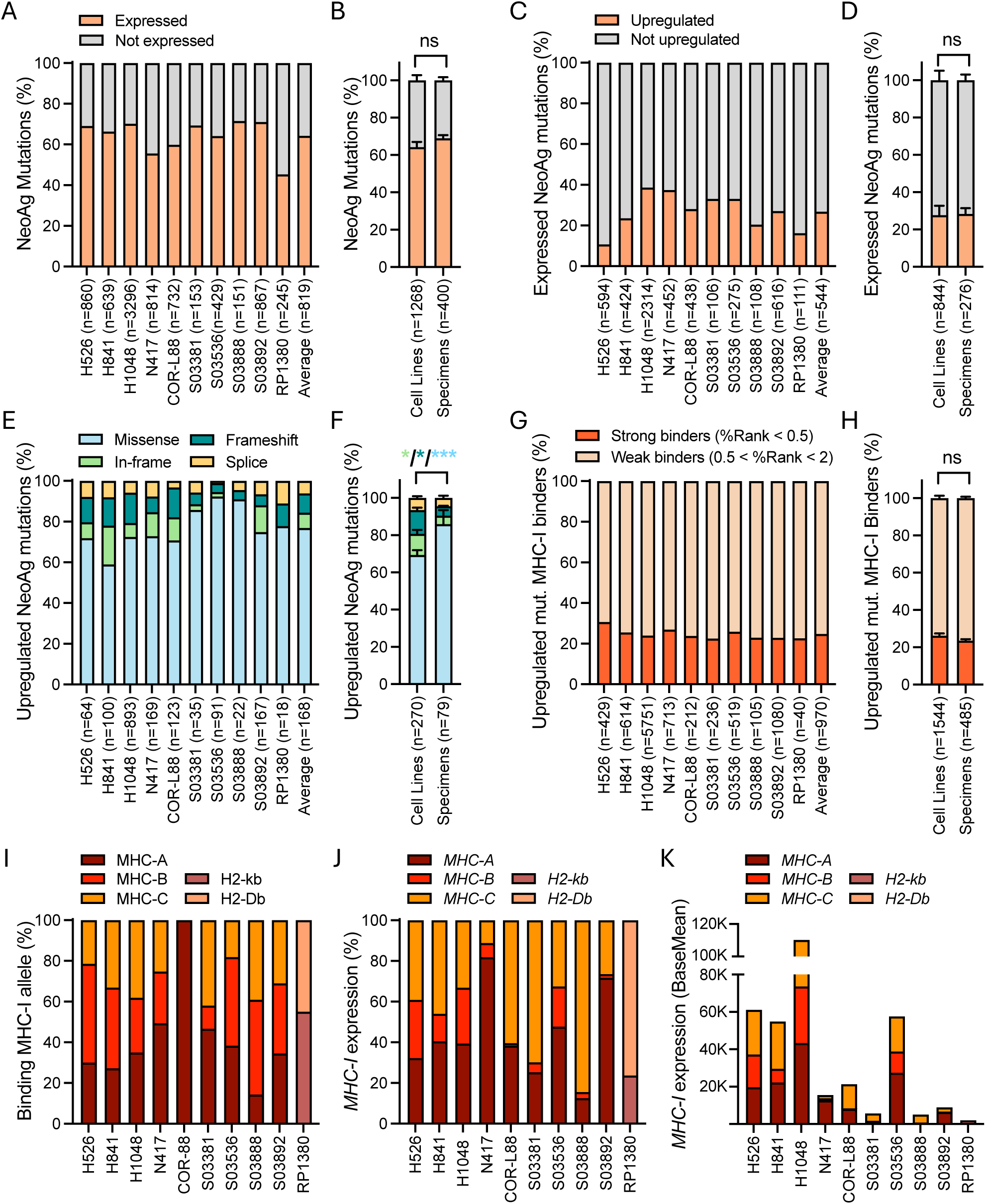
Inhibition of NMD in SCLC samples upregulates hundreds of predicted neoantigens derived from all mutation types. **A-B)** Proportion of mutated genes identified as expressed by transcriptomic sequencing for the indicated SCLC samples (**A**) and comparing of cell line *vs* patient data (**B**). Only potentially NeoAg mutations (missense, in-frame, frameshift and splice) were considered. **C-D)** Proportion of expressed mutated genes found to be upregulated upon SMG1i treatment in the indicated SCLC samples (**C**) and comparing cell line *vs* patient data (**D**). Only potentially NeoAg mutations were considered. **E-F)** Proportion of upregulated NeoAg mutations of different types (**E**) and comparing cell line *vs* patient data (**F**). **G-H)** Mutant MHC-I-binders predicted for upregulated NeoAg-encoding transcripts derived from missense, in-frame and frameshift mutations in the indicated SCLC samples (**G**) and comparing cell line *vs* patient data (**H**). **A-H** indicate proportion (%), with absolute numbers (n = abs) indicated for each sample. Average values (n = abs_mean) are also indicated for all samples, for all cell lines and for all patient specimens. **I)** Proportion of MHC-A/B/C molecules predicted to bind upregulated neoantigens from **G-H**. Note that not all six alleles per sample were available for the binding prediction (**Fig. S10E**). **J)** Proportion of *MHC-A/B/C* alleles expression determined by transcriptomic sequencing calculated from BaseMean expression values. K) *MHC-A/B/C* alleles expression (BaseMean) determined by transcriptomic sequencing. Error bars represent the SEM. ***P-value < 0.001; *P-value < 0.05; (Unpaired t-tests). See also Fig. 5A, C and **Table S5E.**

**Figure S12 – related to Figure 5.**
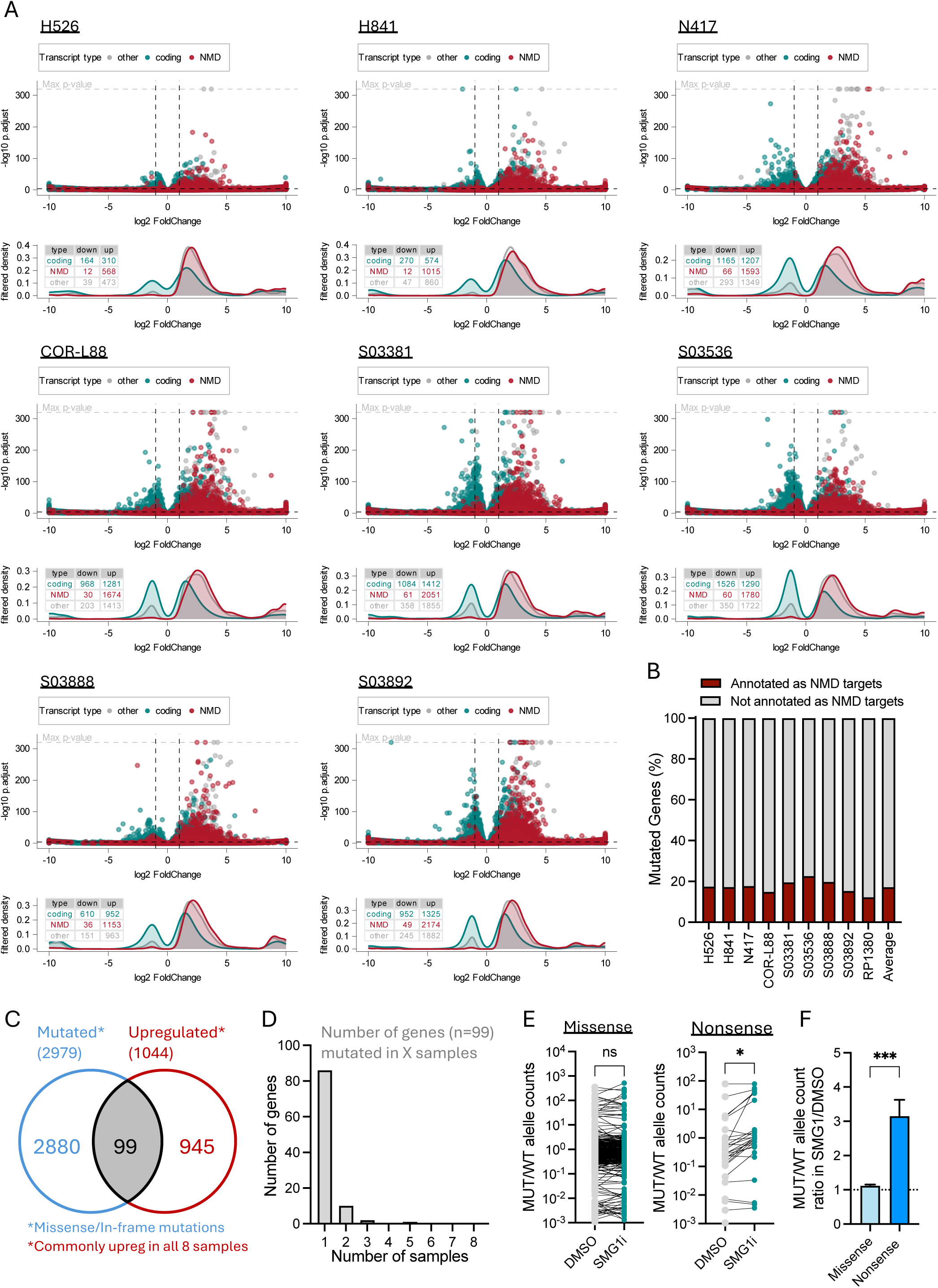
NMD-mediated regulation of missense and in-frame mutated transcripts. **A)** Volcano plots accompanied by filtered densities showing significantly up or downregulated transcripts (Log2FC > |1| and Padj < 0.001) predicted as NMD targets (*NMD*, based on transcript GENCODE annotation as *nonsense-mediated decay* owed to the presence of a PTC at > 50 bp upstream of the next splice site) in comparison to transcripts annotated as *protein-coding* or *non-coding*. Note that transcripts annotated as *nonsense-mediated decay* can also be either protein-coding or non-coding, but the *nonsense-mediated decay* annotation prevails. **B)** Proportion of mutated genes for each sample annotated as NMD targets. **C)** Overlap between of genes containing missense or in-frame mutations in at least one of the 8 tested samples (shown in **A-B**) and genes commonly upregulated in all 8 tested samples upon SMG1i treatment **D)** Number of samples in which overlapping genes identified in panel **C** are found to have a specific missense or in-frame mutation. **E-F)** Split allele count analysis in 8 samples (shown in **A-B**) using matched exome and transcriptome data. The expression ratio of mutated to wild-type alleles (MUT/WT) is shown for all upregulated missense (non-PTC mutation) and nonsense (PTC-mutation) mutated mRNAs in DMSO and SMG1i conditions **(E)** and for the SMG1i/DMSO ratio to show the differential regulation of MUT alleles between missense and nonsense mutations.

**Figure S13 – related to Figure 5.**
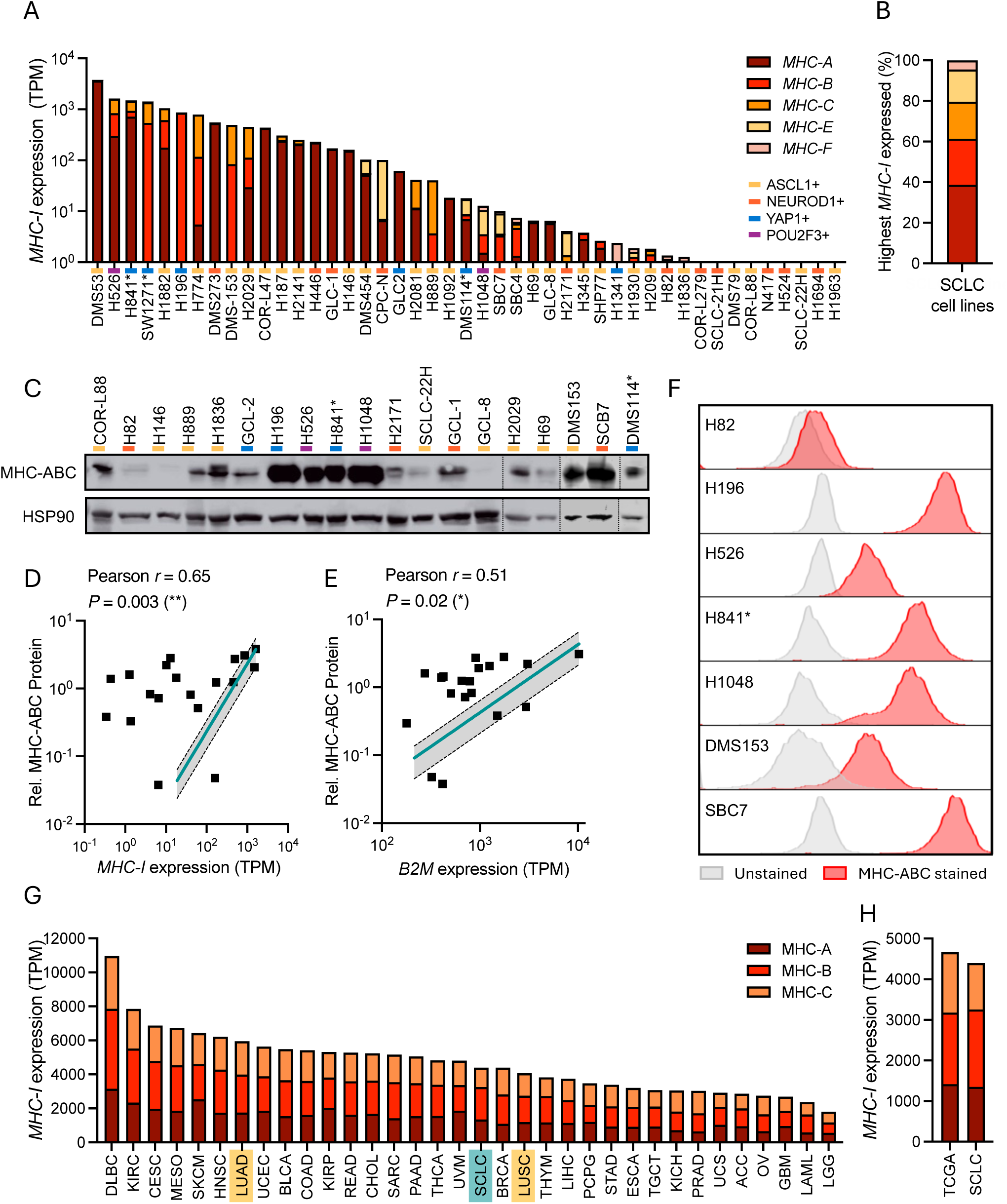
Human SCLC cell lines express MHC-I. **A)** *MHC-I* expression (sum of *MHC-A/B/C/E/F*) determined by transcriptome sequencing of 45 human SCLC cell lines under steady state conditions (with cell lines recently reclassified as SMARCA-UT marked with * and neuroendocrine phenotype indicated: ASCL1+ and NEUROD1+ (NE^High^); POU2F3 and YAP1+ (NE^Low^)). **B)** Proportion of most abundant *MHC-I* genes referring to the highest expressed *MHC-I* for each cell line determined by transcriptome sequencing. 80% of cell lines express mostly classical MHC-A/B/C molecules. Data is displayed following the color code depicted in panel A. **C)** Western blot showing protein levels of MHC-ABC in human SCLC cell lines under steady state conditions (n=19; including recently reclassified SMARCA4-UT as indicated by * and neuroendocrine phenotype as described in **A**). **D-E)** Pearson correlation of *MHC-I* mRNA expression (**D**) or *B2M* mRNA expression (**E**), both derived from transcriptome sequencing data, with MHC -ABC protein levels relative to HSP90 protein levels quantified by densitometry analysis for western blots shown in panel **C**. **F)** Flow cytometry histograms for MHC-ABC surface expression in n=7 human SCLC cell lines. Recent re-classifications as SMARCA4-UT are designated with *. H82 was chosen as a low MHC-ABC-expressing cell line and all other cell lines as high MHC-ABC, confirming expression levels and correlations depicted in previous panels **A-E**. **G)** MHC-ABC expression for SCLC and all cancers within the TCGA database. **H)** MHC-ABC expression in SCLC vs the average of all cancers within the TCGA database.

**Figure S14 – related to Figure 5.**
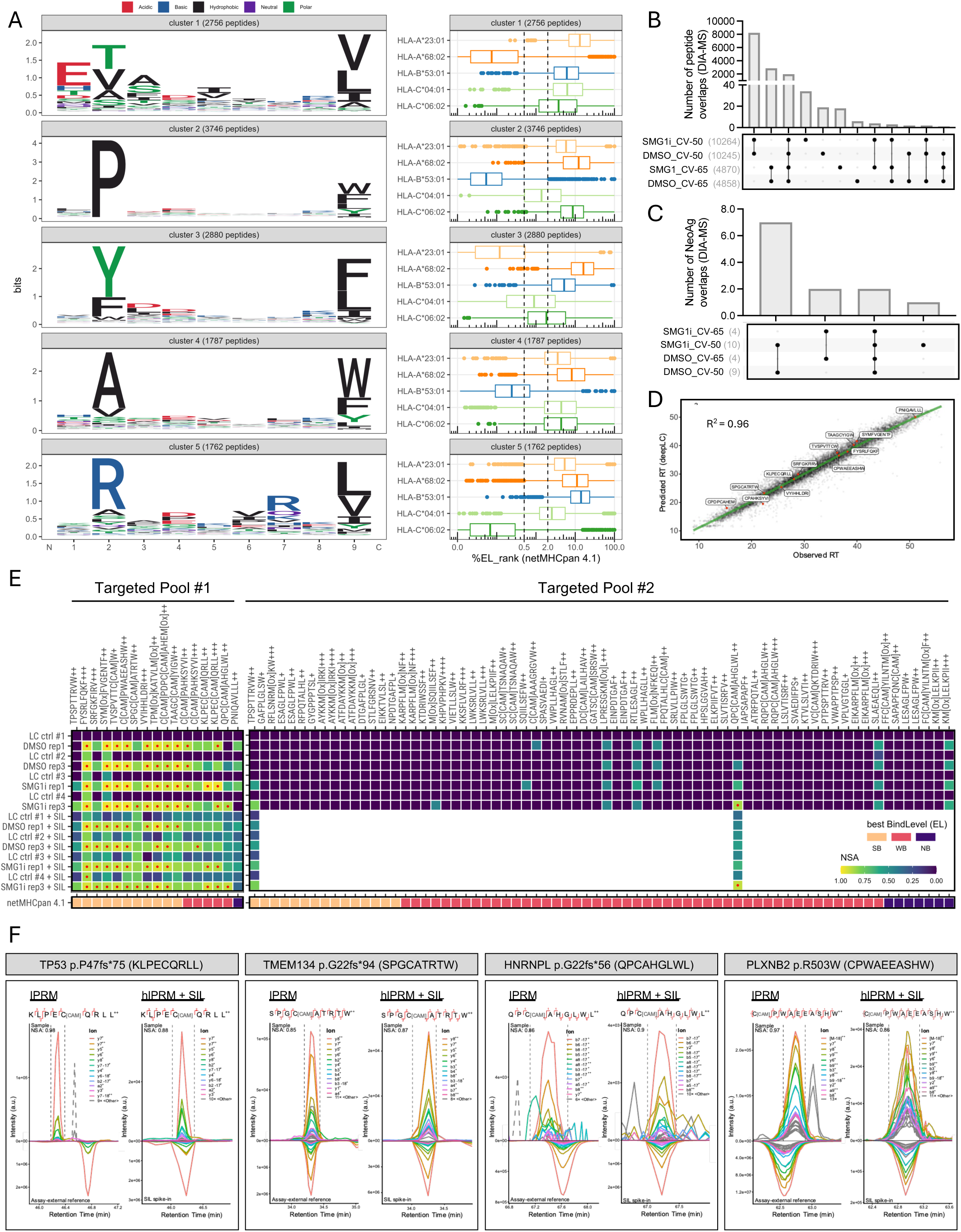
MHC-I immunopeptidomics and targeted proteomics reveal upregulation of presented neoantigens upon NMD inhibition in SCLC. **A)** Analysis for sequence motif enrichment (**left**) for MHC-I binders identified by MHC-I immunopeptidomics via data-independent acquisition mass spectrometry (DIA-MS) and NetMHCpan EL 4.1 binding predictions of peptides constituting the respective motif (**right**). **B-C)** UpSet plots showing number and overlaps of overall peptides (**B**) and *bona fide* neoantigens (**C**) captured by MHC-I immunopeptidomics DIA-MS using two different compensation voltages (CV, -50 and -65). **D**) Correlation of observed and predicted retention times (RT) for all peptides captured by MHC-I immunopeptidomics via DIA-MS, with *bona fide* neoantigens highlighted. **E)** Targeted immunopeptidomics on MHC-IP samples to validate *bona fide* neoantigens identified via DIA-MS (n = 13) and to assess novel neoantigen discovery of predicted neoantigens (n = 58, see **Methods**). NSA = Normalized Spectral Angle; SB = strong binders, WB = weak binders and NB = non-binder based on MHC-I binding predictions via NetMHCpan EL 4.1. **F)** Extracted ion chromatograms from targeted immunopeptidomics (**top**) and comparison with assay-external reference signal (IPRM) or SIL spike-in signal (hIPRM+SIL) for three SMG1i-upregulated neoantigens identified via MHC-IP-DIA-MS and a frameshift neoantigens newly discovered by the targeted approach. See also **Table S5F-G.**

**Figure S15 – related to Figure 6.**
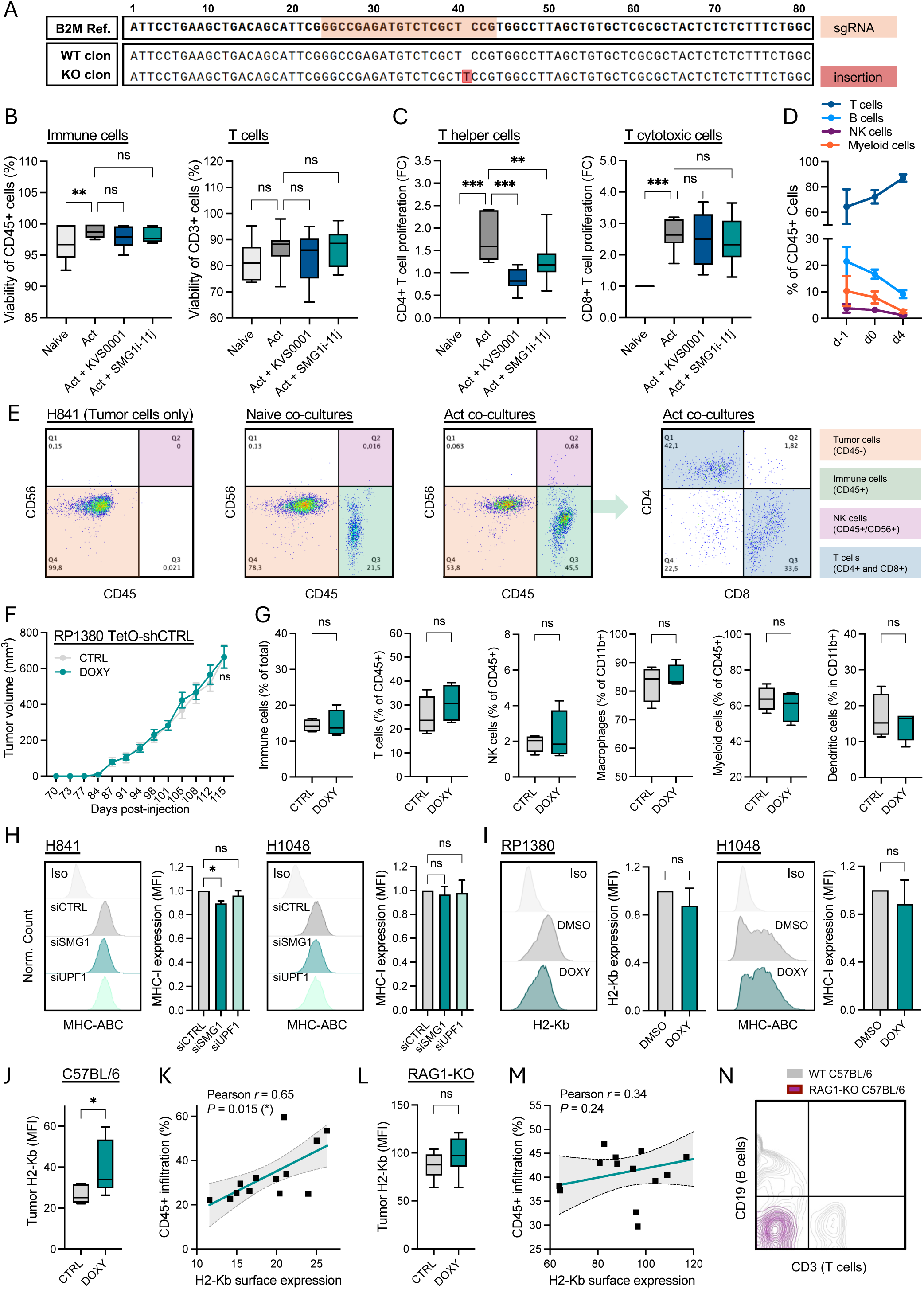
Inhibition of NMD enhances SCLC immunogenicity *in vitro* and *in vivo*. **A)** DNA Sanger sequencing of the amplified B2M locus in representative H841 CRISPR/Cas9 WT and B2M-KO clones aligned to the reference sequence (NCBI, NG_012920.2) with sgRNA and the identified insertion mutation in H841 B2M-KO cells highlighted. **B-D)** Pharmacological NMD inhibition with either KVS0001 or SMG1i-11j compounds in healthy PBMC donors upon T cell artificial activation. (**B**) Proportion of viable CD45+ cells (left) and T cells (right) within PBMC populations after 4 days treatment with a T cell artificial activation cocktail (Act = CD3+CD28+IL-2) *vs* unstimulated conditions (Naive). (**C**) Proliferation of CD4+ (left) and CD8+ (right) T cells derived from T cell counts expressed as fold change (FC) for activated (Act) conditions relative to the unstimulated (Naive) control. (**D**) Proportion of T cells (CD3+), B cells (CD19+), NK cells (CD56+) and Myeloid cells (CD11b+) within PBMC populations (CD45+) from healthy donors directly after thawing (d-1), at the beginning of stimulation (CD3+CD28+IL-2) (d0) 4 days post-stimulation (d4). Graphs represent mean + SEM (n = 7). ***P<0,001; **P<0.01; *P<0.05; ns=non-significant (One-way ANOVA). **E)** Representative flow cytometry dot plots for the staining of CD45 *vs* CD56 (for NK cells) and CD8 *vs* CD4 (both for T cells) following 4 days co-culture of healthy donor PBMCs with WT H841 tumor cells. **F)** *In vivo* tumor growth in immunocompetent C57BL/6J mice transplanted with murine RP1380 TetO-shCTRL fed with normal or doxycycline-containing diet (n ≥ 4). **G)** Flow cytometry immunophenotyping of RP1380 TetO-shCTRL tumors harvested at the end of experiments shown in panel F (n ≥ 4). **H-I)** MHC-I surface expression quantified by flow cytometry in the indicated human and murine cell lines following genetic NMD inhibition via siRNA-mediated SMG1-KD and UPF1-KD (**H**) or doxycycline-inducible SMG1-KD (**I**). **J)** *In vivo* assessment of MHC-I (H2-Kb) surface expression in control vs NMD-inhibited murine RP1380 TetO-shSMG1 tumors (following doxycycline diet, DOXY) grown subcutaneously in C57BL/6 mice. **K)** Flow cytometry data for murine RP1380 TetO-shSMG1 allograft models grown in C57BL/6 mice significantly correlating tumor-specific H2-Kb surface expression with levels of immune cell infiltration (CD45+ infiltration). **L)** *In vivo* levels of tumor-specific MHC-I (H2-Kb) surface expression in control vs NMD-inhibited RP1380 TetO-shSMG1 tumors (DOXY) grown subcutaneously in RAG1-KO C57BL/6 mice. **K)** Flow cytometry data for murine RP1380 TetO-shSMG1 allograft models grown in RAG1-KO C57BL/6 mice correlating tumor-specific H2-Kb surface expression with levels of immune cell (CD45+) infiltration. **L)** Representative flow cytometry contour plot showing T cells (CD45+/CD3+) and B cells (CD45+/CD19+) in the blood from WT C57BL/6 and RAG1-KO C57BL/6 mice.

## Materials and Methods

### Cell culture and SCLC tumor models

All our studies were performed with established immortalized cell lines (ATCC), as well as with SCLC primary tumor cells derived from patient-derived xenotransplant models or genetically engineered mouse models for SCLC. Patient tumors were obtained from biopsies or resections and engrafted subcutaneously onto immunocompromised NOD scid gamma (NSG™) mice to establish patient-derived xenograft models (PDX). Tumors from xenograft mice were then harvested at a maximum volume of 1500 mm^3^, digested to single cells employing an enzymatic tumor dissociation kit (Miltenyi, #130-095-929) and maintained in HITES media (DMEM:F12 containing 10 nM hydrocortisone, 0.005 mg/ml insulin, 0.01 mg/ml transferrin, 30 nM sodium selenite, and 10 nM β-estradiol) supplemented with 10% FBS and 1% Penicillin–Streptomycin antibiotic solution. All patient-derived xenotransplant models were pathologically inspected via H&E stainings and confirmed to have a SCLC histology by at least two independent expert pathologists. All patient-derived samples were handled according to ethical regulations and their use was approved by the institutional review board of the University of Cologne (IRB-ID 19-1164).

Murine SCLC RP and RPM cell lines were established from lung tumors grown in RP (*Rb1^fl/fl^/Tp53^fl/fl^)* and RPM (*Rb1^fl/fl^/Tp53^fl/fl^/Msh2^fl/fl^*) mice, respectively, which developed following adenoviral inhalation of CMV-Cre recombinase. Lung tumors were dissected and digested to single cells by incubation at 37 °C for 2 h while shaking in DMEM media supplemented with 100 U/ml Collagenase IV and 15 mM HEPES^113^. Single cell suspensions were then filtered through a 70 μm strainer, washed once in PBS and cultured in ultra-low attachment flasks with complete HITES media. All immortalized cell lines were obtained from ATCC and their identity was confirmed by STR genotyping. All cells were grown at 37°C under a controlled atmosphere with 5% CO_2_ and 95% relative humidity and with growth media specifications described by ATCC (https://www.atcc.org). Specifically, RP1380 cell line was developed in a previous study^129^. A list for all cell lines and primary cell models employed in this study can be found in **Table S1C**.

### Generation of CRISPR/Cas9*-*knockout cell lines

MHC-I-deficient *B2M-knockout* (B2M-KO) H841 cells were generated via CRISPR/Cas9 targeting the *B2M* sequence 5’-GGCCGAGATGTCTCGCTCCG - 3’ and employing the ribonucleoprotein (RNP) transfection guide from IDT. Lipofectamine RNAiMAX was used to transfect pre-assembled RNP complexes containing ATTO5-labelled sgRNA (IDT, #1075927) and high-fidelity Cas9 (IDT #1081060) in H841 cells. ATTO5-positive cells were single-sorted 24 hours post-transfection (hpt) and individual clones were grown and analyzed via western blot and flow cytometry to evaluate targeted protein depletion (data not shown). Several clones were then sent for Sanger sequencing after selected amplification of the *B2M* locus. A sequence alignment for representative WT and B2M-KO clones as compared to *B2M* genomic sequence NG_012920.2 can be found in **Fig. S14A**.

### NMD pathway inhibition approaches

Successful NMD pathway inhibition was achieved by using siRNA-mediated *SMG1* and *UPF1* transient knockdown systems, doxycycline-inducible *SMG1* knockdown systems and chemical inhibition targeting the SMG1 kinase. All these approaches are further described in the following subsections.

<u>siRNA transfection for transient knockdown</u>: for transient siRNA transfection, Lipofectamine RNAiMAX reagent was used at a 10pmol RNA:2μL Lipofectamine ratio following the manufacturer’s instructions. Genetic siRNA-mediated inhibition of NMD was achieved 72 hpt (as shown by respective protein levels downregulation via western blot analysis as well as NMD target upregulation measured by qPCR) with the following siRNAs: siCTRL (Origene #SR30004), siSMG1 5’-GUGUAUGUGCGCCAAAGUAUUTT - 3’ (Eurofins)^66^ and siUPF1 5’-GAUGCAGUUCCGCUCCAUUUUTT - 3’ (Eurofins)^66^.

Lentiviral transduction for the generation of doxycycline-inducible *SMG1* knockdown cells: We employed a doxycycline-inducible shRNA-based SMG1 knockdown system using constitutive expression of the *E. coli* TetR dimer (thus lacking viral components which could be immunogenic). In this system, TetR represses shRNA transcription in the absence of doxycycline/tetracycline and permits expression upon doxycycline-binding. HEK293T cells were co-transfected with the lentiviral vector pLKO-TetO-shRNA-Puro (Addgene #21915) and the required lentiviral packaging vectors (psPAX2, Addgene #12260 and pMD2.G, Addgene #12259) using lipofectamine2000, and viral supernatant was collected 48 hpt and filtered through a 45 μM syringe strainer. Filtered supernatant was then added diluted in culture media 1:1 to the cells of interest and after 24 h, successfully transduced cells were selected in fresh media containing 1.5-3 μg/ml Puromycin for 4-7 days. pTIG-Tet-shCTRL and pTIG-Tet-shSMG1 vectors^26^ were kindly provided by Prof. Fernando Pastor and shRNA inserts were subcloned into pLKO-(TetO)-shRNA-Puro vectors using *AgeI/EcoRI* restriction sites. The shSMG1 construct was compatible for targeting both human and murine SMG1 genes and targeted a different SMG1 region than the previously employed siRNA. shCTRL inserts were used as controls to determine shSMG1-specific effects. Stably pLKO-TetO-shRNA transduced cells were cultured in 2 μg/ml doxycycline-hyclate (Sigma #D9891) for at least 72 h to induce genetic NMD inhibition *in vitro* via specific SMG1 knockdown. Cell lines modified to express a doxycycline-inducible shRNA targeting *SMG1* (regarded in this study as *TetO-shSMG1 cells*) were transplanted subcutaneously in mouse flanks to create allograft or xenograft tumor models for *in vivo* assays. In this case, tumor-targeted NMD inhibition *in vivo* was achieved by feeding mice with sterilized doxycycline-containing diet (Sniff #A153D71004).

<u>Chemical NMD inhibition:</u> inhibition of the SMG1 kinase activity was achieved upon addition of 0.5 μM SMG1i (compound 11j^60^) to the cell culture media for 24 h, unless indicated otherwise in text, figures or figure legends. This compound was a kind donation from the Cystic Fibrosis Foundation (CFF, https://www.cff.org) as part of their Chemical Compound Distribution Program. The inhibition exerted by this compound was superior to that of other widely used inhibitors reported to target the NMD pathway, namely NMDI14 (50 μM for 48 h), which disrupts UPF1-SMG7 interaction^69^, and amlexanox (AMLX, 100 μM for 48 h), which is an FDA-approved PTC-readthrough drug^68^. Since none of these other compounds yielded sufficient NMD inhibition in SCLC cells, only SMG1i was use in subsequent functional assays. A derivative of compound SMG1i-11j with enhanced solubility called KVS0001^28^ (MedChem #HY-161111) was employed for pharmacological NMD inhibition *in vivo* at 50 mg/kg body weight (mpk).

### Data retrieval from the COSMIC database

To compare the mutational load in different cancer types, the COSMIC database Cancer Browser (https://cancer.sanger.ac.uk/cosmic/browse/tissue; v100) was employed. In the analysis shown in **Fig. 1A** and **Table S1A**, we chose COSMIC over TCGA because COSMIC reports data for two types of neuroendocrine lung tumors SCLC and LCNEC (including mutation calls from our previous studies^2,5^). Within the COSMIC Cancer Browser, for every tissue of interest, the option *include all* was selected for filters *Sub-tissue selection*, *Histology selection* and *Sub-histology selection*, with exception of lung tissue, for which the *Sub-histology selection* was used to extract individual mutational data for lung adenocarcinoma (LUAD), large cell neuroendocrine carcinoma (LCNEC), lung squamous cell carcinoma (LUSC) and small cell carcinoma (SCLC). Only cancer types with more than n = 50 samples were included in the analysis. Additionally, we included data from our own SCLC cohort (*SCLC_George*; n = 447, **Table S1B**).

### Measurement of NMD activity via cell-based luciferase reporter assays

To measure NMD pathway activity, we adapted a previously described and well-validated reporter assay which is based on the regulation of a β-Globin construct containing a premature-termination codon (PTC) by the NMD pathway^56–58^. Plasmids containing a Firefly luciferase (FIR) fused to the β-Globin gene (HBB) in its wildtype form (WT) or with a premature termination codon (PTC) in position 39 were acquired from Addgene (#112084 and #112085) and 3xFLAG-FIR-HBB-WT/PTC inserts were subcloned into a dual luciferase psiCHECK-2 vector (Promega #C802A) using *SacI/XbaI* restriction sites. The psiCHECK2 vector had been previously modified by silent mutagenesis to contain a *SacI* restriction site upstream of the original Firefly ORF. Therefore, the newly generated psiCHECK-2 vector allowed for transcription under the constitutive HSV TK promoter of either HBB-WT or HBB-PTC transcripts N-terminally fused to a firefly luciferase (FIR), and for the expression of a Renilla luciferase (REN) under the control of a separated constitutive SV40 promoter (see **Fig. 1C**). The FIR-HBB-PTC reporter is subjected to NMD regulation owed to the presence of the PTC and an exon-intron structure that allows for an exon-junction complex to sit downstream of the PTC. Therefore, the firefly signal (FIR) of the FIR-HBB-PTC reporter is NMD-sensitive. In contrast, the signal of the FIR-HBB-WT is NMD-insensitive due to the lack of a PTC and is therefore used as a normalization control to account for any PTC-independent regulation of the construct. Additionally, the Renilla luciferase (REN) signal allows to control for transfection efficiency and for cell viability (see **Fig. 1C**).

Multiple SCLC, NSCLC and non-neoplastic cell lines (see all cell lines in **Table 1C**) as well as HEK293 cells were transiently transfected with either the FIR-HBB-PTC reporter plasmid (NMD-sensitive, in short *PTC-reporter*) or the FIR-HBB-WT reporter plasmid (NMD-insensitive, in short *WT-reporter*) using the Lipofectamine2000 reagent at a ratio of 1μg DNA:1μl Lipofectamine following the manufacturer’s instructions. Cells were then harvested 48 hpt, and FIR and REN luminescence signals were consecutively measured within each lysate employing a luminescence plate reader (Tecan i-control, 2.0.10.0) and the Dual Luciferase Reporter Assay Kit (Promega #E1910) following the manufacturer’s instructions. Specifically, measurements were done in white 96-well plates integrating luminescent signals for 500 milliseconds after adding 40 μl of the FIR substrate and then 40 μl of REN Stop&Glo substrate. For each sample, the FIR/REN ratio was calculated to normalize for transfection efficiency and viability differences. Then, the FIR/REN ratio of the PTC-reporter was normalized to the FIR/REN ratio of the WT-reporter to subtract any PTC-independent regulation of the construct. The resulting value showing *relative light units* (RLU) was indicated as “*Luminescence (RLU)”* and represents the normalized expression of the PTC-reporter construct, which is inversely proportional to the NMD pathway activity. Thus, a high NMD activity is reported by a low PTC-reporter expression and *vice versa*. In short: *Luminescence (RLU) = (FIR/REN)_PTC_ / (FIR/REN)_WT._* Data from luciferase reporter assays are shown in **Figs. 1C-G, S1E-F, S2F-G,** and **S3D.**

For luciferase assays upon siRNA-mediated knockdowns, luciferase vectors were transfected 24 h post siRNA transfections. For luciferase assays upon chemical NMD inhibition, luciferase vectors were transfected 24 h prior to SMG1i treatment. In all assays, luciferase signals were always measured 24 h post chemical treatment, 48 h post luciferase transfection and 72 h post siRNA transfection.

### Estimation of NMD activity using a transcriptome-based NMD activity score metric

To estimate NMD pathway activity across different patient-derived samples and SCLC cell lines, we employed a previously published RNA-Seq-derived metric based on the normalized expression of 50 NMD-sensitive genes^43^. We referred to that metric as the *NMD activity score*, which was calculated either for individual samples by plotting normalized expression of all 50 genes per sample, or for a group of related samples such as all patients of a specific cancer type by plotting the NMD activity scores in reference to all 50 genes in all samples (i.e. patients). We employed this metric to estimate NMD activity in publicly available lung cancer datasets^2,5,54,55^, in adult healthy tissue data from the GTEx database (https://www.gtexportal.org/home/; v8), in cancer cell line data from the *Cancer Cell Line Encyclopedia (CCLE)* database (https://sites.broadinstitute.org/ccle/; v22Q2) and in cancer patient data from The Cancer Genome Atlas (TCGA) database (https://www.cancer.gov/tcga; v40). These data can be found across **Fig. 1** and **Fig. S1**.

Specifically, to calculate this metric for all tumors within the TCGA database (indicated in **Table S1C**) while integrating the SCLC_George cohort into the analysis, the following steps were conducted. First, TCGA data was downloaded using the GDC application programming interface. All available cases were selected for presence of data concerning Copy Number Variation (CNV), gene expression and mutations, using as filters *data_category == Copy Number Variation*, *experimental_strategy == RNA-Seq* and *experimental_strategy == WXS*. Subsequently, all available files of the three data types were downloaded for each case. For CNV data, all files with the term *Copy Number Segment* in *data_type* were downloaded. For downloading of the expression data, *workflow_type == STAR – Counts*, and *data_type == Gene Expression Quantification* were used as filters. For downloading of the mutation data, *workflow_type == Aliquot Ensemble Somatic Variant Merging and Masking*, and *data_type == Masked Somatic Mutation* were used as filters.

All files were selected for the sample types *Primary Tumor*, and *Primary Blood Derived Cancer - Peripheral Blood*, the latter corresponding to the TCGA-LAML project. The SCLC mutation and expression data were extracted from our previous work^5^, specifically from Supplementary Tables 3 and 10, respectively. We employed the TCGA expression data in FPKM values in order to allow for comparisons with the SCLC expression data. The combined harmonized dataset comprised 9429 expression and 10154 mutation files. Next, TCGA and SCLC expression data were intersected to contain only mutually present genes (n = 17086) using the Hugo Symbol for selection. For entries sharing the same Hugo Symbol, the average expression was calculated with zero values being excluded. Expression data was only considered if corresponding mutation data was available for the same patient. If, in rare cases, multiple expression data were available for primary tumors from the same patient, expression values were averaged with zero values being excluded. If several mutation files were available for a single patient, the mutation counts were averaged. To estimate the mutational burden depicted in **Fig. 1H**, only exome mutation types were considered filtering for *Missense, Nonsense, Nonstop, In_Frame_Del, In_Frame_Ins, Frame_Shift_Del, Frame_Shift_Ins, Splice_Site* and *Silent_Mutation*.

For the calculation of the *NMD activity score*, each *NMD-sensitive gene* expression value was normalized to the median expression of selected *non-targets* for the same patient. Selected NMD-sensitive genes and selected non-targets were extracted from the original work developing this metric^43^. Finally, the NMD activity score was plotted for each sample, with each dot representing the NMD activity score of each considered NMD-sensitive gene (**Fig. S1C**), or for each cancer type, with each dot representing the NMD activity score of each donor (GTEx, **Fig. S1B**); patient (Lung datasets, **Fig. 1B**; TCGA, **Fig. 1H-J**); cell line (CCLE, **Fig. 1K-M**).

### *In vitro* enzymatic phosphorylation assays

A biochemical enzymatic assay on purified components employing 436 recombinant kinases was performed by Eurofins service KinaseProfiler upon treatment with 10 nM, 100 nM and 1000 nM SMG1i. These assays included radiometric protein kinase assays and HTRF® lipid and atypical kinase assays in mostly human kinases and some murine kinases. Each assay was individually optimized for the specific kinase regarding buffer conditions, required cofactors and substrates as well as optimal amount of ATP. A detailed protocol employed for each kinase assay can be obtained upon request at Eurofins. This data can be found in **Figs. 2D-E** and **S3B**, and **Table S2A**.

Additionally, an inhibitory curve was performed for full length purified SMG1 incubated with eight increasing SMG1i concentrations (including 10, 100 and 1000 nM) and the absolute IC50 was interpolated from a non-linear adjustment using Prism (GraphPad). Specifically, per well, 12 ng of purified full length SMG1 protein was incubated with 30 nM GST-cMyc-p53 as a phosphorylation substrate and 10 um ATP for 30 min at RT in reaction buffer (25 mM HEPES pH 8.0, 0.01% Brij-35, 1% Glycerol and 10 mM MnCl2). D2-labelled anti-GST monoclonal antibody + Europium-labelled anti-phospho-P53 Ser15 antibody were used for detection via homogeneous time resolved fluorescence (HTRF) using a ratio of 10000x (Em665nm/Em620nm). This data can be found in **Fig. S3A**. This assay was later repeated to compare the IC50 of both SMG1 kinase inhibitors SMG1i and KVS0001 and the data is found in **Fig. 4A**.

### Cellular SMG1i on/off-target inhibitory assay

To estimate the IC50 for SMG1i-mediated on-target NMD inhibition in cellular assays, luciferase PTC-reporter assays were conducted. Cells were first transfected with the luciferase HBB-PTC reporter construct, harvested 24 hpt and replicate-plated into several wells for their treatment with increasing concentrations of SMG1i for the next 24 h. FIR and REN luciferase signals readout was conducted 24 h post-treatment and 48 h post-transfection as described in a previous section to estimate the degree of NMD inhibition under each concentration, which was inversely proportional to the HBB-PTC reporter signal. An inhibitory curve could be then used to interpolate the absolute IC50 via a non-linear adjustment using Prism (GraphPad). This data is shown in **Fig. S3D**.

For the evaluation of the apparent affinity of SMG1i on the PIP5K2C/PIP4K2C kinases (possible off-target inhibition of PIP5Kg), NanoBRET^TM^ assays were conducted at Eurofins. These assays measured the competition of the NanoBRET^TM^ tracer in HEK293-H cells transiently transfected with NanoLuc® luciferase-kinase Fusion Vector using the NanoBRET^TM^ detection method designed to measure the molecular proximity in living cells. Such NanoBRET^TM^ Target Engagement (TE) Intracellular Kinase Assays were performed according to the Promega Adherent Format Assay Protocol using a NanoBRET^TM^ Tracer. HEK293-H cells were transiently transfected with a NanoLuc® Fusion Vector, seeded in 384-well plates, and cultivated overnight. Target engagement of the compound was measured by treating the cells with the SMG1i compound in presence of a fluorescent tracer at a fixed concentration or in presence of DMSO for background corrections (no tracer control); the reaction was performed for 120 minutes at 37°C under 5% CO2 followed by 15 minutes at RT for temperature equilibration. Substrate of Nanoluc® was then added. After 2 minutes, the donor emission (NanoLuc®) and acceptor emission (fluorescent tracer) were measured at respectively λex=450 nm and λem=610 nm using a microplate reader (Envision, Perkin Elmer). Bioluminescence Resonance Energy Transfer (BRET) ratio was determined by dividing the acceptor emission value by the donor emission value. Background correction was performed by subtracting the measured BRET ratio in absence of a tracer (no-tracer control) from the BRET ratio for each sample. The standard inhibitory reference compound (Foretinib) was tested in each experiment at several concentrations to obtain an inhibition curve to determine the IC50 value. This data is shown in **Fig. S3E-F**.

### Proliferation assays

For proliferation assays, 1x10^5^ cells were stained with 5 μM of proliferation dye eFluor670 (Invitrogen #65-0840-85), diluted in PBS (1 ml of PBS per 1x10^6^ cells) and plated in duplicate wells for treatment with either DMSO as a control or 0.5 μM SMG1i for NMD inhibition. eFluor670 fluorescence intensity of single living cells was initially measured at day 0 and then after 4-6 days by flow cytometry using the APC channel (see respective section). The number of cell divisions was estimated assuming a half-decrease of the mean fluorescence intensity (MFI) per division with the following formula: Division Number = log_2_ [(MFI_day0_ – MFI_unstained_)/(MFI_dayX_ – MFI_unstained_)]. The average cycling time was calculated by dividing the number of cell divisions by the experimental end time (typically 6 days unless indicated otherwise). This data is shown in **Figs. 2I** and **S4**.

### Cell death assays

For cell death assays, cell growth (confluence) and cell death were monitored via live-cell imaging for 4 days every 6 hours using an Incucyte S3 device (Sartorius). Cells were treated with either DMSO or 0.5 μM SMG1i in the presence of several cell death inhibitors, namely 10 μM of the pan-caspase inhibitor QvD for apoptosis inhibition (MCE #HY-12305), 5 μM Ferrostatin-1 (Fer-1) for ferroptosis inhibition (MCE #HY-100579) or 20 μM of the RIPK1 inhibitor necrostatin-1s (Nec1s) for necroptosis inhibition (MCE #HY-14622A). Additionally, the dye DRAQ7 (Invitrogen #D15106) was added to stain and quantify dead cells over time. Specifically, 5x10^4^ cells per well were plated in a 48-well plate, each condition was plated in triplicates and 4 photos per well were taken at the indicated times. At the end of the experiment, confluence over time was normalized to the initial confluence to account for cell seeding fluctuations and was expressed as confluence ratio, and cell death over time was quantified based on DRAQ7 dye intensity and normalized to the respective confluence for each given well and timepoint. This data is shown in **Figs. 2J-P** and **S5.**

### Cell viability assays

The decrease in cell viability upon SMG1i treatment was monitored employing the CellTiter-Glo® (CTG) Luminescent Cell Viability Assay (Promega #G7570). For generation of viability curves, 5x10^3^ cells per well were plated in flat-white 96-well plates with serial increasing concentrations of SMG1i (triplicate wells per concentration) and incubated under cell culture standard conditions. After 24, 48 and 72 h, 100 μl of CTG substrate diluted 1:5 in PBS were added per well, plates were incubated for 10 minutes at RT in the dark and luminescence signal was measure in a luminescence plate reader (Tecan i-control, 2.0.10.0). Signals were converted to cell viability (%) by normalization to the untreated control, and viability curves were used to estimate GI50 values via non-linear adjustment using Prism (GraphPad). These data is shown in **Figs. 2Q, 2U** and **S6**.

Similarly, the CTG assay was used to determine cell viability in several cell lines and patient-derived cells after 3 days (not shown) and 6 days of chemical (SMG1i) or genetic (DOXY) NMD inhibition. In this case, cells were treated in 12 well plates, with media and compound replacement every 3 days. For the readout, half of the cells were harvested with the other half remaining under treatment. Harvested cells were thoroughly resuspended in 500 μl media and 3x 100 μl of each cell suspension was transferred to a flat-white 96-well plate to proceed as specified above. For viability assays in doxycycline inducible SMG1-KD systems, the viability of TetO-shSMG1 cells was normalized to the viability of parental control cells to account for any doxycycline-associated toxicity. This data can be found in **Figs. 2V**, **3F** and **4H-I.**

### Flow cytometry

To prepare samples for flow cytometry, cells were washed once with PBS to then generate single cell suspensions with Trypsin-EDTA. Cells were washed once in FACS buffer (PBS + 2% FCS + 2 mM EDTA) before addition of FcR blocking reagent (Miltenyi, human #130-059-901, murine #130-092-575) and incubation for 10 min on ice. Cells were stained with the desired antibodies for 15 min on ice in the dark, washed once with FACS buffer and resuspended in FACS buffer containing 0.25 μg/ml DAPI for flow cytometric analysis with a MACSquant16 Analyzer (Miltenyi). The device was compensated and calibrated as specified by the manufacturer using compensation beads (#130-104-693) and calibration beads (#130-093-607), respectively. Flow cytometry data was analyzed with Flowjo software. Human primary antibodies used were B2M (Molecular probes #A15770), MHC-ABC (Invitrogen #11998342), CD45 (Miltenyi #130110636), CD3 (Miltenyi #130113142), CD4 (Biolegend #357410), CD8 (Miltenyi #130110678), CD56 (Miltenyi #130113310), CD11b (Miltenyi #130110556). Murine primary antibodies used were CD45 (Miltenyi #130110665), CD3 (Miltenyi #130119793), Nkp46 (BD Biosciences #3122669), CD19 (Miltenyi #130112037), CD11b (Miltenyi #1301138063), CD14 (Miltenyi #130115559), H2-Kb (Miltenyi #130115586), IFN-γ (ThermoFisher #48731182), KI-67 (Miltenyi #130120418). Flow cytometry data can be found in **Fig. 6B**, **6F**, **6H-I**, **6K-L**, **S15B-E** and **S15G-N.**

### Cytosolic calcium measurements

For the detection of cytosolic calcium increase during ER stress-related cell death, cells were treated for 72 h with 0.5 μM SMG1i or 1-10 μM tunicamycin (TUNIC), which served as a positive control for ER stress-related cell death. Cells were then stained for cytosolic calcium by adding for 1 h at 37 °C 1 μM of the calcium indicator Fluo-4AM (ThermoFisher #F14201) to the cell culture media. Afterwards, cells were harvested in PBS, washed once in FACS buffer, stained with DAPI and subsequently analysed by flow cytometry. Calcium staining was detected in the FITC channel by gating on yet-living DAPI-negative single cells. This data can be found in **Fig. 3E**.

### Viability rescue by inhibition of the ER stress response pathway

To rescue SMG1i-induced cell death, H1048 and GLC-1 (not shown) cells were treated for 72 h with 0.5 μM SMG1i; timepoint at which death became apparent. Co-treatment (72 h) with increasing concentrations of several ER stress sensors inhibitors were used to rescue the observed cell death phenotype. The following compounds were used: PERK inhibitors GSK2606414 (PERKi, MCE #HY-18072) and ISRIB Trans-Isomer (T-ISRIB, MCE #HY-12495); and the IRE1 inhibitor MKC-3946 (IREi, MCE #HY-19710). Concentration ranges employed for each compound are specified in **Fig. 3H**.

### Protein extraction and western blotting (WB)

Cell pellets were lysed in RIPA buffer (20 mM Tris–HCl pH 7.5, 150 mM NaCl, 1 mM Na2EDTA, 1 mM EGTA, 1% NP-40, 1 mM Na3VO4, 1% sodium deoxycholate, 2.5 mM sodium pyrophosphate, 1 mM β-glycerophosphate) supplemented with phosphatase inhibitors (Roche tablets #04906837001) and protease inhibitors (Roche tablets #11836170001) for 20 min on ice and lysates were pre-cleared by centrifugation at maximal speed and 4°C. Protein concentrations were quantified using the Pierce BCA assay kit (Thermo Scientific #23225) according to the manufacturer’s instructions. Protein lysates were denatured in Lämmli Buffer (60 nM Tris pH 6.8, 10 % glycerol, 2 % SDS, 5 % β-mercaptoethanol, 0.003% bromophenol blue) for 10 min at 95°C and run in 4-20% gradient Tris-Glycine gels (Invitrogen, #15531242) and 1X SDS running buffer (25 mM Tris, 192 mM glycine, 0.1% SDS) for 2h at 120 V. Proteins were then transferred to a PVDF membrane using an iBlot3 transfer system (Invitrogen) at 25V for 7 min and low cooling and membranes were blocked with 5% milk powder dissolved in 1X TBST (50 mM Tris–HCl pH 7.6, 150 mM NaCl, 0.05% Tween-20) before incubating them overnight at 4°C with the desired primary antibodies under rotation. Membranes were then washed three times for 5-10 min with 1X TBST and incubated with a suitable fluorophore-coupled secondary antibody for 1 h at RT in the dark followed by three 5-10 min washes with 1X TBST. Membranes were imaged in an Odyssey® DLx Imaging System (LI-COR) and densitometry analysis was conducted with the Image Studio Lite software v5.0.21 (LI-COR).

Primary antibodies used were FIREFLY 1:500 (Abcam #ab21176), SMG1 1:500 (CST #4993), P-UPF1-Ser1127 1:500 (Sigma-Aldrich #07-1016), UPF1 1:1000 (CST #9435), UPF1 1:500 (Bio-Rad #VMA00627), RB1 (Abcam AB181616), P53 (CST #2524), MSH2 (Thermo Fisher #33-7900), NCAM1/CD56 (Proteintech #14255-1-AP), BiP (CST #3177), p62 1:500 (CST #5114), ATF4 1:1000 (CST #11815), CHOP 1:500 (CST #2895), BAK1 1:1000 (CST #3814), STAT3 1:1000 (CST #9132), P-STAT3-Tyr705 1:1000 (CST #9145), AKT 1:1000 (CST #2920), P-AKT-Ser473 1:1000 (CST #4060), 4EBP 1:1000 (CST #9644), P-4EBP-Thr37/46 1:1000 (CST #2855), HLA-ABC 1:1000 (Abcam #ab703228), B2M 1:500 (Abcam #ab75853), HSP90 1:1000 (CST #4877), GAPDH 1:2000 (CST #2118), TUBG 1:1000 (Abcam #ab7291), ACTB 1:1000 (Santa Cruz #sc-47778). Secondary antibodies were anti-Rabbit 680 1:5000 (CST #5366), anti-Rabbit 800 1:10000 (CST #5151), anti-mouse 680 1:5000 (CST #5470) and anti-mouse 800 1:10000 (CST #5257). WB data is found in **Figs. 1C, 2S, 3D, 3G, 5E, 6C, S1F, S2B-E, S3C, S7C, S8A, S8D-E, S8H-I** and **S13C**.

### RNA extractions

To prepare samples for qPCR studies, frozen cell pellets were used to extract RNA employing the Quiazol Lysis reagent (Qiagen #79306) following the manufacturer’s instructions. RNA pellets were then dissolved in RNase-free water and quantified with the use of a Nanodrop 200C spectrophotometer (Thermo Scientific).

To prepare samples for transcriptome sequencing, RNA extractions were performed with the Qiagen RNAeasy Mini Kit according to the instructions of the manufacturer. Specifically, RNA extractions from cell pellets were performed with the RNeasy Mini Kit (Qiagen #74104) following the provided protocol “Purification of Total RNA from Animal Cells using Spin”, and RNA extraction from fresh-frozen tissues processed with Tissue-Lyser LT (Qiagen #85600) was done with RNeasy Micro Kit RNA (Qiagen #74004) following the provided protocol “Tissue-Lyser LT from Qiagen”. RNA quality and concentration were assessed with the TapeStation (Agilent #50675576), and samples with an RNA integrity number (RIN) of at least 7 were subjected to RNA sequencing.

### qPCR

500 ng of RNA were subjected to cDNA synthesis using the HiScript III 1st Strand cDNA Synthesis Kit (NeoBiotech #NB-54-0182-02). The cDNA was then diluted 1:5 in nuclease-free water and specific gene amplification was performed in a QuantStudio5 thermocycler (Applied Biosystems) using specific primer pairs in ChamQ Universal SYBR-Green qPCR master mix (Vazyme #Q711). Relative quantification of gene expression was calculated using the ΔΔCt method and β-Actin (ACTB) was used as a housekeeping gene for normalization. The following primers were used for relative quantification of NMD targets: *ATF3* For 5’-CACTGGTGTTTGAGGATTTTGCTAA-3’, Rev 5’-GCAGCTGCAATCTTATTTCTTTCTC-3’; *SMG5* For 5’-GACCTGAGTGAAGGCTTTGAATCG-3’, Rev 5’-GCCATTGAGGGAATCGGGAGC-3’; *TBL2* For 5’-GCAGTCATTTACCACATGC-3’, Rev 5’-TATTGTTTCTGCTTCTTGGAT-3’; *ACTB* For 5’-ATCAAGATCATTGCTCCTCCTGAG-3’, Rev 5’-CTGCTTGCTGATCCACATCTG - 3’; *GAPDH* For 3’-AGTCAGCCGCATCTTCTTTTGC- 5’, Rev 3’-GTTAAAAGCAGCCCTGGTGACC -5’. For size restriction reasons, this study shows only results for *TBL2* as one representative *bona fide* NMD target, but in every case qPCR quantifications were conducted for all NMD targets indicated above (data not shown). NMD targets and non-targets were additionally validated at NMDRHT*^67^* (https://nmdrht.uni-koeln.de). qPCR data can be found **in Fig. 4E, 5D, S2H-I** and **S8F**.

### Transcriptome sequencing and data processing

For bulk RNA sequencing, extracted RNA was used to generate cDNA libraries of Poly-A-selected mRNA with the Illumina TruSeq protocol for messenger RNA. Then, short read paired-end (2x 100 bp) sequencing was performed employing an Illumina HiSeq or NovaSeq platform and aiming at 50 Mio reads to achieve a sequencing depth comparable to that of previous studies in SCLC^2–5^ and typically TCGA datasets, a depth which also enables identifying low-expression transcripts. Reads were aligned against the human genome (GRCh38/h38, GENCODE release 42 transcript annotations^130^ supplemented with SIRVomeERCCome annotations from Lexogen) using the STAR read aligner (version 2.7.10b)^131^. Transcript abundance estimates were computed with Salmon (version 1.9.0)^132^ in mapping-based mode using a decoy-aware transcriptome (GENCODE release 42) with the options -- numGibbsSamples30 --useVBOpt -- gcBias -- seqBias. After the import of transcript abundances in R (version 4.3.0) using tximport (version 1.28.0)^133,134^differential transcript expression (DTE) analysis was performed with the DESeq2 R package (version 1.40.1)^134^ including batch information in the design formula. Transcripts with less than 10 counts in half the analyzed samples were pre-filtered and discarded. The DESeq2 Log2FoldChange (Log2FC) estimates were shrunk using the apeglm R package (version 1.22.1)^135^. Processed transcriptome sequencing data for all samples can be found in **Tables S2 (Fig. S1A** and **Fig. 2F-H)** and **S3 (Fig. S1A** and **Fig. 3A-C).**

General significance cut-offs were Log2FC > |1| & Padj < 0.05 unless indicated otherwise. Significantly upregulated and downregulated transcripts were selected for each sample and then compared across all samples (which included a total of 8 different samples, four immortalized cell lines - one per transcription factor-based SCLC subtype - and four patient-derived primary samples) using the multiple list comparator of Molbiotools (https://molbiotools.com/listcompare.php) to evaluate overlaps. The UpSetR package (version 1.4.0)^136^ was used to plot overlaps between samples (**Fig. S3A**).

Genes which transcripts were found significantly up/downregulated upon SMG1i treatment in at least 6 out of 8 samples were used for pathway enrichment analysis using gProfiler (https://biit.cs.ut.ee/gprofiler/gost) and including Gene Ontology Biological Processes (GOBP), KEGG pathways, Wikipathways (WP) and REACTOME pathways annotations. To avoid well-known redundancies as part of this analysis, all genes associated with significantly enriched cell cycle-, apoptosis- or ER stress-related terms were manually collected under a single topic term and significantly deregulated transcript expression for those genes was plotted (**Fig. 3B**); provided that such transcripts were annotated as *protein-coding* and were expressed in at least 7 out of 8 samples. Overlaps and pathway enrichment analyses can be found in **Table S4.**

Significantly de-regulated transcripts annotated by GENCODE as *nonsense-mediated decay* or as *intron retention* (*transcript type* column in **Table S3**) were considered *PTC-transcripts* (**Fig. 2H** and **Fig. 3C**). For each sample, corresponding expression and mutation data were intersected using base R (version 4.3.0) to evaluate upregulation of neoantigen-encoding transcripts, considering only one mutated upregulated transcript per gene to avoid counting redundancies (**Fig. 5C** and **Table S5D**).

### DNA extraction

For patient-derived material, DNA was extracted from fresh-frozen tumor material previously propagated as xenotransplant models in immunocompromised NSG mice. All tumor samples derived from murine xenotransplant models showed a tumor content of at least 95% in H&E staining. When available, DNA was also extracted from patient-matched fresh-frozen blood which served as a normalization control to filter out germline variants. For cell lines, DNA was extracted from frozen pellets and matched controls for normalization were not available. DNA extractions were then conducted with the Puregene DNA extraction kits (Qiagen) for blood (#158023), cells (#158043) and tissues (#158063) following the manufacturer’s protocol.

### Whole exome sequencing

For whole-exome sequencing, the Twist Human Core Exome kit (Twist Bioscience) was used to prepare exon-enriched libraries which were then subjected to 2x100 bp paired-end sequencing at a mean insert size of 200 base pairs (bp) on the Illumina NovaSeq platform aiming for a 150X coverage. Sequencing reads of human samples derived from xenotransplant models were aligned to a combined human and murine reference genome (GRCh37/hg19 and GRCm38/mm10) and data analyses were performed on reads mapped to the human genome to thus exclude murine sequencing reads. For human cells sequencing reads were mapped to the human genome h19 and for murine cells to the murine genome mm10. Further analysis for tumor purity, tumor ploidy, somatic mutations and copy number alterations were performed with our in-house pipeline *Peiflyne*^3–5^ (http://www.uni-koeln.de/med-fak/peiflyne/peiflyne.tgz). Processed exome sequencing results for all samples can be found in **Suppl. Table 5A**. Data derived from exome sequencing can be found in **Fig. 1A, 1H-J, 1K-M, 2P, 2T, 4G, 5A-E, 5H-I, S9-11** and **S12B-F.**

Split analysis for wildtype and mutant allele counts was conducted for nonsense and missense mutations to compare the number of mutant (MUT) alleles to the respective wildtype (WT) allele in DMSO *vs* SMG1i-treated cells. To this end, the frequencies of single-nucleotide variants (SNVs) were examined in 48 samples, comprising n=3 RNA sequencing data files per condition from four cell lines (H526, H841, H1048, COR-L88) and four SCLC patient-derived primary cells (S03381, S03536, S03888 and S03892). Mapping to the reference genome GRCh37/hg19 was performed using the RSEM tool (https://github.com/deweylab/RSEM, version 1.3.1). The obtained BAM files and corresponding processed whole-exome sequencing data, filtered to contain only missense or nonsense mutations identified in transcripts found upregulated upon NMD inhibition, were subsequently analysed using the Vartrace module of our in-house developed tool Peiflyne. The resulting MUT and WT allele counts were averaged for each for the three replicates of each condition, and MUT/WT ratios were compared between DMSO and SMG1i conditions. These results can be found in **Fig. S12E-F**.

### MHC-I binding prediction

Mutational information derived from whole exome sequencing was used for the *in-silico* generation of sample-specific mutant peptide libraries using an in-house pipeline. The analysis specifically focused on specific types of coding mutations affecting the exome of the human or murine genome. Specifically, missense mutations, in-frame indels and frameshift indels were considered, while silent, nonsense and splice mutations were not considered. Mutation coordinates and mutation nomenclatures derived from exome data were used to retrieve wild-type sequences for mutated genes and to incorporate determined mutations in specified locations. Full length mutated nucleotide (nt) sequences were then translated to amino acid (aa) sequences and trimmed for subsequent analyses. For missense and in-frame mutations, peptide sequences were trimmed to include the mutated amino acid(s) in a centered position preceded by 13 aa upstream and followed by 13 aa downstream of the mutated site. Frameshift and nonstop mutant sequences were trimmed only 13 aa upstream of the mutation start while preserving downstream the altered peptide sequence following the frameshift. The employed trimming lengths enabled prediction of MHC-I binders for all possible sizes fitting the MHC-I peptide pocket (8mers-14mers = Xmers). *MHC-I* locus genotyping (**Fig. S10E**) was conducted at high resolution using a Luminex 200^TH^ device (Luminex Corporation)^137^ and the LABType SSO Class I A/B/C Locus Typing Test kits (#RSSO1A, #RSSO1B and #RSSO1C, One Lambda, Thermo Fisher Scientific) by the Institute of Hematology and Transfusion Medicine at the University of Bonn.

Sample-specific peptide libraries and MHC-I-genotyping information were then used to predict the binding of mutant peptide-derived Xmers (choosing all sizes, 8-14mers) to sample-specific MHC-I molecules with the online MHC-I-binding tool NetMHCpan 4.1 from the immune epitope data base (IEDB, http://tools.iedb.org/mhci/)^85,86^. From all Xmers generated for a particular mutant peptide library, binders were selected based on a percentage (%) rank below 2 and categorized Into strong (%rank < 0.5) and weak binders (0.5 < %rank < 2), according to IEDB guidelines. Note that the chosen trimming strategy allows to evaluate binders of all sizes (8-14 aa) but some of the output 8-13 aa predicted binders do not contain the mutation. Sample-specific mutant peptide libraries employed for prediction of neoantigens via MHC-I-binding modeling can be found in **Table S5C** and predicted MHC-I binders can be found in **Table S5D**.

### MHC-I immunopeptidomics and targeted immunopeptidomics mass spectrometry

MHC-I immunopeptidomics was performed with H1048 cells (see MHC-I typing in **Fig. S10E**) due to their high number of somatic mutations and predicted MHC-I binders (**Fig. S9-11**) and their high levels of HLA-I surface expression (**Fig. S13C-F**), in order to maximize the likelihood of capturing neoantigens. To this end, 5×10^7^ cells were treated with 0.5 µM SMG1i or DMSO as a control for 48 h and were harvested on ice-cold PBS by gentle scrapping to preserve surface epitopes. Immunoprecipitation of MHC-I:peptide complexes was performed as previously described^138^ with in-house produced panHLA antibodies (clone W6/32 and additional steps for the forced oxidation of methionine using H2O2 and reduction and alkylation of cysteine using tris(2-carboxyethyl)phosphine (TCEP) and iodoacetamide (IAA). For quantitative purposes, three biological replicates were included in the analysis, and a fraction of one of the replicates, together with a fourth newly included replicate, were employed for hit validation via targeted immunopeptidomics.

Lyophilized peptides were dissolved in 12 µl of a 5 % ACN/ 0.1 % TFA solution and spiked with 0.5 µl of 100 fmol/µl Peptide Retention Time Calibration (PRTC) Mixture (Pierce, #88321) and 10 fmol/µl JPTRT 11 (a subset of peptides from the Retention Time Standardization Kit (JPT, #RTK-1-100), and then transferred to QuanRecovery Vials with MaxPeak HPS (Waters, #186009186).

Untargeted sample analysis was conducted with an UltiMate 3000 RSLCnano system coupled to an Orbitrap Exploris 480 equipped with a FAIMS Pro Interface (Thermo Fisher Scientific). For chromatographic separation, peptides were first loaded onto a trapping cartridge (Acclaim PepMap 100 C18 μ-Precolumn, 5μm, 300 μm i.d. x 5 mm, 100 Å; Thermo Fisher Scientific) and then eluted and separated using a nanoEase M/Z Peptide BEH C18 130A 1.7µm, 75µm x 200mm (Waters, #186008794). The total analysis time was 120 min and separation was performed at a flow rate of 0.3 µl/min with a gradient starting from 1 % solvent B (100 % ACN, 0.1 % TFA) and 99 % solvent A (0.1 % FA in H2O) for 0.5 min. Concentration of solvent B was increased to 2.5 % in 12.5 min, to 28.6 % in 87 min and then to 38.7 % in 1.4 min. Subsequently, concentration of solvent B was increased to 80 % in 2.6 min and kept at 80 % solvent B for 5 min for washing. Finally, the column was re-equilibrated at 1 % solvent B for 11 min. The LC system was coupled on-line to the mass spectrometer using a Nanospray-Flex ion source (ThermoFisherScientific), a SimpleLink Uno liquid junction (FossilIonTech) and a CoAnn ESI Emitter (Fused Silica 20 µm ID, 365 µm OD with orifice ID 10 µm; CoAnn Technologies). The mass spectrometer was operated in positive mode and a spray voltage of 2400 V was applied for ionization with an ion transfer tube temperature of 275°C. For ion mobility separation, the FAIMS module was operated with standard resolution and a total carrier gas flow of 4.0 l/min. Each sample was injected twice using either a compensation voltage (CV) of -50 V or -65 V for maximal orthogonality and thus increased immunopeptidome coverage. Full Scan MS spectra were acquired for a range of 300 – 1650 m/z with a resolution of 120000 (RF Lens 50 %, AGC Target 300 %). MS/MS spectra were acquired in data-independent mode (DIA) using 44 previously determined dynamic mass windows optimized for MHC class I peptides with an overlap of 0.5 m/z. HCD collision energy was set to 28 % and MS/MS spectra were recorded with a resolution of 30.000 (normalized AGC target 3000 %).

FAIMS-DIA MS data was analyzed using the Spectronaut software (version 17.6; Biognosys)^139^. Raw files recorded with different CVs were searched separately against the UniProtKB/Swiss-Prot database using the directDIA approach (retrieved: 21.10.2021, 20387 entries) as well as against a database containing the H1048-specific mutant peptide library generated as specified above from H1048 whole exome data (see Table S5B). Search parameters were set to non-specific digestion and a peptide length of 7-15 amino acids. Carbamidomethylation of cysteine and oxidation of methionine were included as variable modifications. Identification results were reported with 1 % FDR at the peptide level. All peptides identified by Spectronaut were further analyzed by NetMHCpan 4.1 binding predictions^86^, Gibbs 2.0 clustering of peptide sequences^140^, and retention time prediction by DeepLC^141^. Quantification of HLA class I-presented peptides was performed as described before^15^ using the raw output at the MS2 level from Spectronaut with cross-run normalization disabled. Peptides with a fold change in abundance > 2 and with an FDR ≤ 0.05 were defined as “hits” while peptides with a fold change ≥ 1.5 and an FDR ≤ 0.2 were defined as “candidates”. Additionally, we implemented a hybrid imputation method using knn imputation for values with “missing at randomness”, e.g. peptides detected in 2 out of 3 replicates anda MinDet imputation for peptides which were not detected in any of the replicates of a condition due to their abundance under the limit of detection. Of note, the TP53 frameshift peptide presented in **Fig. 5H-I** was only found in SMG1i-treated samples and its values were imputed using the MinDet method for DMSO-treated control samples.

MHC-I immunopeptidomics data was intersected with matched transcriptome and genome data to assign peptides which 1) derived from de-regulated mRNAs (**Fig. 5H, left panel**) and 2) derived from mutated genes (**Fig. 5H, right panel**). In addition, a tumor associated antigen (TAA) *in silico* library was compiled by integrating information from four different tumor antigens and cancer testis antigens databases, namely CEDAR (https://cedar.iedb.org), CAPED (https://caped.icp.ucl.ac.be), caAtlas (https://www.zhang-lab.org/caatlas/) and TANTIGEN (https://projects.met-hilab.org/tadb/). Using this library, significantly up/downregulated peptides were filtered for candidate TAA. Next, assigned TAA peptides ever found presented in human adult healthy tissues were filtered out using the PCI database (https://pci-db.org), which contains information on > 10 Million MHC-I/II-presented peptides obtained from > 3000 samples derived from > 40 different healthy human adult tissues^142^.

Targeted immunopeptidomics validation of mutation-derived peptides was performed using optiPRM, a parallel reaction monitoring method with increased sensitivity due to per-precursor collision energy optimization^143^. Briefly, all candidates identified by FAIMS-DIA as well as 58 additional frameshift mutation-derived predicted H1048 HLA-A/B binders (see more details about the selection of these peptides in the next section) were obtained as stable isotope labelled (SIL) synthetic peptides (JPT Peptide Technologies GmbH). PRM acquisition was performed using the following parameters: MS data were recorded at a resolution of 90,000 at 200 m/z, over the mass range from 205 to 1285 m/z, with 300 % AGC target and 25 ms maximum IT. MS2 data were acquired with PRM scans at a resolution of 120000 at 200 m/z. The target precursor list was provided with preselected charge states, the corresponding m/z and optimized CE values and narrow isolation window (≤1 m/z) tuned per precursor, and their expected retention time (±1.5 min.) predefined with SIL peptides and indexed to a set of retention time standard peptides. The normalized AGC target was set to 1000 %. The maximum injection time mode was set to dynamic, aiming for a coverage of at least five points across the chromatographic peak. The dynamic RT feature using the PRTC mixture was active. Protonated polycyclodimethylsiloxane (a background ion originating from ambient air) at 445.12 m/z served as a lock mass. For spike-in experiments, 10 fmol of SILs were added. The target list was reduced to peptides with evidence from the initial measurement. MS2 data of target peptides was acquired with a resolution of 480000 at 200 m/z while additional MS2 data of SILs was acquired with a resolution of 45000 at 200 m/z, normalized AGC target of 200 % and a maximum injection time of 350 ms using the optimized CE values, narrow isolation windows and predefined and indexed expected retention time (±1.5 – 5 min). Data were manually analyzed using Skyline and R as described before^143^ by comparing spectra recorded from SIL peptides with spectra recorded from the sample. Normalized spectral angles (NSAs) were calculated as described before^144^.

### Neoantigen immunogenicity evaluation via Fluorospot assays

In order to evaluate the immunogenicity of identified presented neoantigens found upregulated in SMG1i-treated H1048 cells (NeoAg pool#1, n=3) as well as of several predicted frameshift neoantigens (NeoAg pool#2, n=136) (**Table S5G**), FluoroSpot assays were conducted. In brief, healthy donor PBMCs are mixed with peptides at high concentrations which can then bind to cell surface MHC-I molecules of antigen presenting cells subsequently leading to the recognition by autologous T cells within the same PBMC population. Upon neoantigen recognition, T cells become activated and secrete IFN-γ, which can be detected in an ELISA-like fashion as the reaction is performed with antibody-coated plates for the desired secreted cytokine.

Healthy-donor PBMCs were thawed as specified in a previous section, washed once in PBS, dissolved in pre-warmed RPMI media supplemented with 20% FBS and incubated for 1 h at 37 °C. In the meantime, FluoroSpot plates (Mabtech kit #FSP-0102-2, all required antibodies provided) were washed 3x with 200 µl PBS per well and then blocked with 100 µl per well of RPMI 10% FCS for 30 min at RT. Afterwards, 2x10^5^ PBMCs in 50 µl reaction media per well supplemented with 0.1µg/ml anti-CD28 were seeded and 50 µl of reaction media containing the desired stimuli were added, analyzing triplicates for each condition. Conditions included a negative control (0.1µg/ml anti-CD28 only), a positive control (0.1ug/ml anti-CD28 + 0.1ug/ml anti-CD3), a human actin peptide pool control (Peptides & Elephants #LB02233), a viral peptide pool control (Peptides & Elephants #LB01359), and the H1048-specific neoantigen peptide pool #1 and pool #2. All peptide pools were used at 1 ug of each peptide/ml concentration. Plates were incubated at 37°C for 18-20 h before discarding the supernatant, followed by 5 washing steps with 200 µl PBS per well and 2h incubation at 37 °C with the primary antibody solution (2.5 ug/ml anti-IFN-γ #7-B6-1-FS-BAM diluted 1:200 in 0.1% BSA/PBS buffer). The supernatant was then discarded, and plates were washed 5 times with 200 µl PBS per well followed by 1h incubation at RT with the secondary antibody solution (1:200 anti-BAM-490 diluted 1:200 in 0,1% BSA/PBS buffer) and 5 more washing steps with 200 µl PBS per well. The plates were then incubated with 50 µl per well of fluorescence enhancer and incubated for 15 min at RT in the dark. The solution was then discarded, the protection underdrain was removed, and plates were dried at RT in the dark for at least 24 h until the color of the grey membranes turned to white. The plate fluorescence was imaged and quantified using an AID iSpot Spectrum Reader (AID, Germany).

### Peptide pools rational design

Synthetic peptides were used in this study for **1)** neoantigen validation and discovery via targeted proteomics (peptides purchased from JPT) and **2)** neoantigen immunogenicity testing via FluoroSpot assays (peptides purchased from Peptides & Elephants).

For targeted proteomics, the three bona fide neoantigens identified via MHC-I immunopeptidomics which presentation was upregulated in SMG1-treated samples (**Fig. 5H-I**) were obtained as stable isotope labeled (SIL) reference peptides. together with the other 10 MS-identified bona fide neoantigens, which all derived from missense mutations (Targeted Pool#1, **Table S5G**). Additionally, 58 peptides were obtained as SIL reference peptides and screened by optiPRM with the aim of capturing predicted neoantigens. Here, we selected only frameshift mutations present in H1048 cells from genes for which mRNAs levels were found upregulated in SMG1i-treated samples, and for which corresponding wild type peptides had been identified via MHC-I immunopeptidomics as upregulated in SMG1-treated samples (n = 23 frameshift mutant genes). All unique 9mers predicted to bind any of the HLA-A or HLA-B alleles expressed by H1048 (HLA-C is very low expressed in this cell line, **Fig. S13A**) were selected for screening (Targeted Pool #2, **Table S5G**).

The immunogenicity of the three bona fide neoantigens identified by HLA-I immunopeptidomics for which presentation was upregulated in SMG1-treated samples was tested via Fluorospot assays (NeoAg Pool #1, **Table S5G**). The immunogenicity of 136 additional peptides was also evaluated (NeoAg Pool #2, **Table S5G**). For the design of this pool, we selected only frameshift mutations identified in H1048 cells from genes which mRNAs were found upregulated in SMG1i-treated samples, and for which corresponding wild type peptides had been identified by HLA-I immunopeptidomics also upregulated in SMG1-treated samples (n = 23 frameshift mutant genes). In silico translated amino acid sequences for those selected frameshift mutations were then used to predict MHC-I 9mer binders using NetMHCpan EL 4.1, since MHC-I has a remarkably higher preference to bind 9 aa peptides over all other sizes (**Fig. 5F**). Because these peptides were analyzed in FluoroSpot assays using healthy donor PBMCs with unknown HLA-genotype, the MHC-I binding prediction was conducted for the top three European MHC-A/B/C alleles (HLA-A*02:01/01:01/03:01; HLA-B*07:02/08:01/15:01; HLA-C*04:01/03:04/07:01) according to the Stefan-Morsch-Foundation database (611746 donors) in order to maximize the likelihood of random matches.

### *In vitro* tumor and immune cell co-cultures

The human cell line H841 was chosen for our *in vitro* co-cultures because of its high tumor mutational burden leading to a high number of potential neoantigens, its high transfectability, its high MHC-I expression (as we showed in **Fig. S13**) and more importantly because four of its MHC-A/B/C alleles, namely MHC-A*03:01, MHC-B*07:02, MHC-C*07:01 and MHC-C*07:02, are all among the top three most frequent MHC-A/B/C alleles – each within it class respectively - within the European population (http://allelefrequencies.net/top10freqsc.asp). Using this H841 cell line thus increases the likelihood of random matches. Indeed, HLA-I typing of two randomly chosen PBMC donors revealed that each of them shared two HLA-I alleles with H841 (data not shown). PBMCs were isolated from the blood of healthy donors via Ficoll centrifugation gradient and frozen in FCS supplemented with 20% DMSO. Osmotic careful thawing was conducted by placing vials for 5 min at 37°C in a bead bath, carefully transferring the cell solution to a 50 ml falcon and slowly adding dropwise pre-warmed T cell media (RPMI 1640 supplemented with 10% FBS, 1% Pen-strep, 10 mM HEPES, 1 mM sodium pyruvate, 2 mm L-Glutamine and 50 μM β-mercaptoethanol), before centrifuging the cells for 3 min at 500 g and resuspending pellets in T cell media followed by an overnight culture at 37°C to allow cells for recovery. On the same day, H841 WT or *B2M*-KO cells were transfected with siCTRL/siSMG1/siUPF1, and medium was replaced 6 hpt to avoid toxicity. The next day, PBMCs were prepared as naive (unstimulated), primed with 1000U/ml hIL-2 (Peprotech #200-02) and 1 μg/ml anti-CD28 (BioLegend #302934) or artificially activated (Act) by stimulation with 1000U/ml hIL-2, 1 μg/ml anti-CD28 and 1 μg/ml anti-CD3 (BioLegend #317326) right before their co-culture with H841 cells in a 10:1 ratio in 96-well plates in T cell media (1x10^4^ H841 cells and 1x10^5^ PBMCs per well). After 4 days, co-cultured cells were harvested and stained with anti-CD45 and DAPI (0.25 μg/ml) to quantify single living tumor cells (CD45-) and T cells (CD45+) via flow cytometry (see respective section). The generation of *B2M*-KO cells is described in a previous section. Co-culture data is shown in **Fig. 6A-D** and **Fig. S15E**. Healthy donor PBMCs were also used for studying the effects of SMG1i and KVS0001 treatment on the activation of T cells (shown in **Fig. S15B-C**). In this case 0.5 μM of SMG1i or KVS0001 were added to the activation cocktail specified above (CD3+CD28+IL-2) and living CD4+ and CD8+ T cell counts in naive *vs* activated conditions were employed to determine T cell proliferation levels. Healthy donor PBMCs were also used for the evaluation of immune cell populations present at day -1 (thawing), day 0 (stimulation starts) and day 4 (naive vs activated) by flow cytometry as specified in a previous section. This data is shown in **Fig. 15D**.

### *In vivo* experiments and mouse lines

For *in vivo* experiments in full immunocompromised mice, NOD scid gamma (NSG) mice (Charles River) were injected with 1.5-3x 10^6^ cells per flank for the desired models (parental control cells and TetO-shSMG1 cells) while inhaling isoflurane as narcosis. Mice were grouped to receive a specific treatment once tumors reached 100 mm^3^ to evaluate a possible regression of established tumors. For systemic pharmacological NMD inhibition, NSG mice were injected intraperitoneally (ip) with 100 ul of either vehicle control solution (10%DMSO; 40%PEG; 5% Tween-80; 45%PBS) or the SMG1 inhibitor compound KVS0001 at 50 mg/kg (mpk) for maximum three weeks in a 5-days-on-two-days-off regimen. For tumor-targeted genetic NMD inhibition using doxycycline-inducible SMG1-KD models (TetO-shSMG1 cells), mice were fed with either a normal diet or a doxycycline-containing diet (Sniff #A153D71004) for the duration of the experiment. Mice weight and tumor growth were monitored twice a week. Following ethical regulations, mice were sacrificed when at least one of its tumors reached a volume of 1000 mm^3^ or a tumor length of 15 mm. Once all control mice had been terminated, mice receiving treatment within the same experiment were also terminated. This data is found in **Fig. 4**.

For *in vivo* experiments in immunocompetent mice, C57BL/6J mice (Janvier) were injected subcutaneously with 1.5-3x10^6^ RP1380 TetO-shCTRL or RP1380 TetO-shSMG1 cells per flank. Following engraftment of the cells, mice were immediately grouped to receive either a normal diet or a doxycycline-containing diet to induce the targeted knockdown of *SMG1* in RP1380 TetO-shSMG1 cells. This data is found in **Figs. 6E-F**, **6M-N** and **S15F-G**. To exclude the influence of the immune system in the observed phenotypes, these experiments were repeated in fully immunocompromised NSG mice (Charles River) (**Fig. 4K**) and partially immunocompromised RAG1-KO C57BL/6J mice lacking B and T lymphocytes^110^ (**Fig. 6G-I** and **Fig. S15J-N**). Mice weight and tumor growth were monitored at least once a week. For these experiments, all mice were sacrificed simultaneously for subsequent tumor analysis as soon as the average tumor volume reached the termination criteria in the control group.

Immune checkpoint blockade treatment with anti-PD-1 (BioXCell, *InVivo*MAb clone RMP1-14) was conducted on RP1380 TetO-shSMG1 allograft C57BL/6J models receiving doxycycline-diet directly after tumor cell injection to induce steady-state SMG1-KD. These mice received either 100 ul PBS or anti-PD-1/PBS (200 ug/mice), injected intraperitoneally three times per week for four weeks, and treatment started for each mouse independently once tumors reached a size of 50-100 mm^3^. Mice were sacrificed after completing four weeks of anti-PD-1 treatment. In this case, tumor volumes were measured twice per week and normalized to the exact initial tumor volume at treatment start to show tumor growth changes under treatment as volume fold change. Related data is shown in **Fig. 6J-L**.

In all cases, tumor volumes (V) were calculated following the elliptical model as V = 0.5 × L × W^2^, where L is the tumor length and W is the tumor width. All graphs for *in vivo* tumor growth (shown in **Fig. 4** and **Fig. 6**) show the end timepoints indicated in the *x*-axes (days post-injection) are the actual experimental end time points at which mice were sacrificed according to the ethical guidelines or experimental plan and tumors were extracted for further analysis.

All animal experiments were approved by and conduced in accordance with the regulations of the State Agency for Nature, Environment and Consumer Protection of the State of North Rhine-Westphalia (LANUV, *Landesamt für Natur, umwelt und Verbraucherschutz Nordrhein-Westfalen*, AZ 81-02.04.2018.A002 and AZ 81-02.04.2022.A394).

### *In vivo* pharmacokinetics for the SMG1 inhibitor KVS0001

The *in vivo* pharmacokinetics (PK) of KVS0001 was serviced by KYINNO BIOTECHNOLOGY (Study #E0273-S2401). In this case, the SCLC cell line H82 was implanted subcutaneously in female NSG mice and once tumors reached 300 mm^3^, mice received a single ip dose of KVS0001 either at 25 or 50 mpk. Following KVS0001 administration blood and tumors were collected at various timepoints (as indicated in **Fig. 4C-D**) and processed to measure free the presence of KVS0001 compound via mass spectrometry (LC-MS/MS). A full report of methods and results provided by KYINNO BIOTECHNOLOGY are available upon request. Relevant PK parameters are shown in **Fig. S8B.**

### Single Cell TCR sequencing

Single cell TCR sequencing was performed for tumors grown *in vivo* in immunocompetent C57BL/6J mice. Tumors were extracted and enzymatically digested to single cells in HBSS solution containing 0.1 g Collagenase IV (Sigma Aldrich #C0130), 10 mg Hyaluronidase (Sigma Aldrich #H6254) and 2000 U DNase IV (Sigma Aldrich #D5025). Tumor digestions were immunophenotyped via flow cytometry (see respective section below) to quantify different intra-tumoral immune cell populations. A fraction of the tumor digestions was stained with anti-CD45 and subjected to FACS for CD45+ cell sorting in a BD FACSARIA Fusion device (BD Biosciences). CD45+ cells as single cell suspensions were prepared with a targeted cell recovery of 10.000 cells per sample on a 10X-Chromium X/iX device (10X Genomics) for 5’ TCR sequencing with the Chromium Single Cell Mouse TCR Amplification Kit (10X Genomics #PN-1000254) following the instructions of the manufacturer. Sequencing was performed on a NovaSeq aiming for 50.000 read pairs per cell for each sample; and for TCR enriched libraries covering a sequencing depth of 5.000 read pairs per sample. Data processing was performed with CellRanger software (v7.0.0, 10x Genomics)^145^ to perform V(D)J transcript sequence assembly, alignment to a reference genome (mm10), filtering for low quality data and obtaining the count matrix. Then, *CellRanger multi* pipeline was implemented to infer TCR clonality. TCR sequencing data can be found in **Fig. 6M-N** and **Table S5H.**

## Data availability

The raw sequencing data is available at the European Genome-Phenome Archive and will be made available upon publication of this work (preliminary registration ID: #1176). The mass spectrometry proteomics data have been deposited to the ProteomeXchange Consortium via the PRIDE partner repository^146^ with the dataset identifier PXD066995.

## Supplemental information

<u>Table S1 (Related to Fig. 1) – NMD activity analysis across cancer types</u>

S1A. Mutation count in cancers from the COSMIC database.
S1B. Mutation count in the SCLC_George cohort.
S1C. List of all cellular models employed experimentally in our study.
S1D. TCGA cancer types and abbreviations.

<u>Table S2 (Related to Fig. 2) – Specificity and reproducibility NMD inhibition methods</u>

S2A. Full kinome assay for SMG1i compound.
S2B. DESeq2 DTE analysis for H841 treated with DMSO vs 0.5 um SMG1i for 24 h.
S2C. DESeq2 DTE analysis for H841 treated with siCTRL/siSMG1/siUPF1 for 72 h.
S2D. Overlaps in upregulated transcriptome for all samples in S2B and S2C.
S2E. Overlaps in downregulated transcriptome for all samples in S2B and S2C.

<u>Table S3 (Relative to Fig. 3 and Fig. 5) – DESeq2 DTE analysis for SMG1i/DMSO samples</u> including cell lines (H526, H841, H1048, N417, COR-L88) and SCLC patient-derived primary cells (S03381, S03536, S03888 and S03892).

<u>Table S4 (Related to Fig. 3) – Cellular processes affected upon NMD inhibition in SCLC</u>

S4A. Overlaps in the upregulated transcriptome for all samples in S3.
S4B. Overlaps in the downregulated transcriptome for all samples in S3.
S4C. Pathway enrichment analysis for commonly upregulated genes from S4A.
S4D. Pathway enrichment analysis for commonly downregulated genes from S4B.

<u>Table S5 (Related to Fig. 5 and Fig. 6) – Inhibition of NMD increases SCLC immunogenicity</u>

- S5A. Exome sequencing data for several human cell lines and SCLC specimens.
- S5B. Absolute numbers of mutations and predicted MHC-I binders per sample.
- S5C. Sample-specific *in silico*-generated mutant peptide libraries.
- S5D. Sample-specific predicted mutant MHC-I binders (predicted neoantigens).
- S5E. Exome-transcriptome data intersection: expression of mutated genes.
- S5F. MHC-I immunopeptidomics: presented peptides in H1048 (SMG1i/DMSO).
- S5G. Peptide pools employed in targeted proteomics and FluoroSpot assays.
- S5H. TCR sequencing.

